# A Universal Language for Finding Mass Spectrometry Data Patterns

**DOI:** 10.1101/2022.08.06.503000

**Authors:** Alan K. Jarmusch, Allegra T. Aron, Daniel Petras, Vanessa V. Phelan, Wout Bittremieux, Deepa D. Acharya, Mohammed M. A. Ahmed, Anelize Bauermeister, Matthew J. Bertin, Paul D. Boudreau, Ricardo M. Borges, Benjamin P. Bowen, Christopher J. Brown, Fernanda O. Chagas, Kenneth D. Clevenger, Mario S. P. Correia, William J. Crandall, Max Crüsemann, Tito Damiani, Oliver Fiehn, Neha Garg, William H Gerwick, Jeffrey R. Gilbert, Daniel Globisch, Paulo Wender P. Gomes, Steffen Heuckeroth, C. Andrew James, Scott A. Jarmusch, Sarvar A. Kakhkhorov, Kyo Bin Kang, Roland D Kersten, Hyunwoo Kim, Riley D. Kirk, Oliver Kohlbacher, Eftychia E. Kontou, Ken Liu, Itzel Lizama-Chamu, Gordon T. Luu, Tal Luzzatto Knaan, Michael T. Marty, Andrew C. McAvoy, Laura-Isobel McCall, Osama G. Mohamed, Omri Nahor, Timo H.J. Niedermeyer, Trent R. Northen, Kirsten E. Overdahl, Tomáš Pluskal, Johannes Rainer, Raphael Reher, Elys Rodriguez, Timo T. Sachsenberg, Laura M. Sanchez, Robin Schmid, Cole Stevens, Zhenyu Tian, Ashootosh Tripathi, Hiroshi Tsugawa, Kozo Nishida, Yuki Matsuzawa, Justin J.J. van der Hooft, Andrea Vicini, Axel Walter, Tilmann Weber, Quanbo Xiong, Tao Xu, Haoqi Nina Zhao, Pieter C. Dorrestein, Mingxun Wang

**Author notes:** These authors contributed equally to the work (Alan K. Jarmusch, Allegra T. Aron).

## Abstract

Even though raw mass spectrometry data is information rich, the vast majority of the data is underutilized. The ability to interrogate these rich datasets is handicapped by the limited capability and flexibility of existing software. We introduce the Mass Spec Query Language (MassQL) that addresses these issues by enabling an expressive set of mass spectrometry patterns to be queried directly from raw data. MassQL is an open-source mass spectrometry query language for flexible and mass spectrometer manufacturer-independent mining of MS data. We envision the flexibility, scalability, and ease of use of MassQL will empower the mass spectrometry community to take fuller advantage of their mass spectrometry data and accelerate discoveries.

## Main Text

Despite the widespread use of mass spectrometry (MS) in science to characterize proteins, peptides, polymers, small molecules, and nucleic acids, it remains difficult for scientists to search for known patterns of chemical classes within and across MS data sets. The variety of applications of MS and the diversity of class-specific chemical patterns makes automation difficult. Searches for specific chemicals or specific chemical classes within MS data are performed manually or using specialized software tools. These tools are generally designed to search for a specific pattern, to search data within a single research domain^1^, or were created by computational scientists for their own use that present a high barrier for reuse^2^. This inability of the wider MS userbase to mine MS data rapidly across different MS datasets has left potential discoveries hidden in the data. To address the need for universal searching of MS data, we created MassQL, an open-source MS query language for flexible and mass spectrometer manufacturer-independent mining of MS data.

Based upon the concept that MS data captures unique characteristics of chemical structures, such as isotopic patterns (e.g., bromination), diagnostic fragmentation (e.g., product ion of sulfur trioxide), and neutral loss (e.g., loss of sugar moieties), MassQL implements common MS terminology to build a consensus vocabulary to search for MS patterns in a single mass spectrometry run up to entire data repositories (**Fig. 1a, 1b**). The MassQL language encompasses formal definition of common MS terms, including MS1 patterns, such as precursor ion *m/z* or isotopic patterns, and MS/MS fragmentation patterns (including support for data-dependent acquisition and data-independent acquisition, e.g., SWATH and MS^e^), as well as terms for separation methods, including retention time and ion mobility drift time. Since this terminology is agnostic to the type of mass spectrometry data acquisition used, the MassQL querying language is compatible with all MS data. Additionally, MassQL query options include parameters for setting user-defined tolerances, such as ion intensities and mass accuracies, and boolean conjunctions, such as AND/OR, can be used to create more complex pattern queries marking inclusion or exclusion criteria (**Fig. 1b**). To facilitate adoption of MassQL, we made use of community input to establish commonly used terms required for a succinct language that could be readily shared and reused and new terms can be defined, which enables grammar and syntax evolution of MassQL to maintain compatibility of queries to advancing MS technologies. The resulting MassQL language provides users the flexibility and expressiveness to query simple and complex MS patterns within their data and across public data regardless of their expertise in computational MS. Community members have written and applied MassQL queries to their own research (**SI Notes 1.1 - 5.2**) using MS/MS spectral information (**SI Notes 1.1 - 2.8**), precursor isotopic patterns (**SI Notes 3.1 - 3.9**), drift time (**SI Notes 4.1 - 4.2**), and other parameters (**SI Notes 5.1 - 5.2**) for querying MS data sets for chemically and biologically relevant molecules, such as identification of iron-binding compounds (**SI Note 3.3.1 and 3.3.2)** and distinguishing glycoconjugates (**SI Note 2.8**), among others.

**Figure 1.**
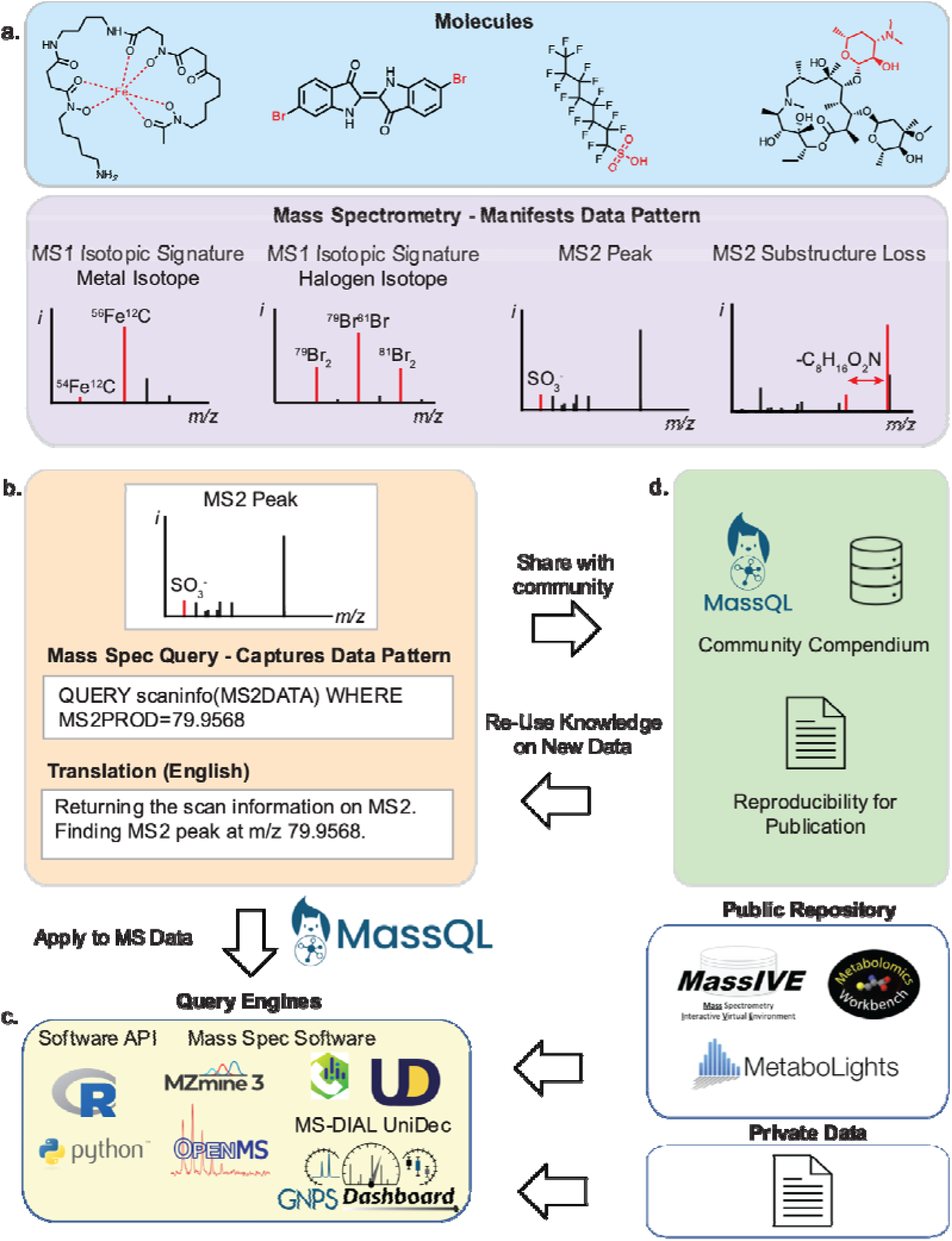
a) Examples of molecules that produce distinctive data patterns when measured by mass spectrometry. b) MassQL query representing MS/MS fragmentation patterns that encapsulates a characteristic mass loss. The query can be translated to 9 languages for enhanced accessibility. c) MassQL is a universal tool to query MS data. MassQL enables data searching in a single file to entire mass spectrometry repositories. MassQL has also been incorporated into a wide range of mass spectrometry software. d) MassQL queries are shared and reused via the Community Compendium, which increases reproducibility and knowledge dissemination.

MassQL is supported natively in a variety of community supported MS software, including MZmine^3^, pyOpenMS^4^, MS-DIAL^5^, UniDec^6^, GNPS^7^, and the GNPS Dashboard^8^ (**Fig. 1c**). To spur more widespread integration of MassQL into other platforms, MassQL is available as Python and R libraries and as a web API (for integration into software tools using a programming language without official libraries, including Java, Scala, C#). Further, a standalone command line tool and portable scalable computational workflow^9^ are available to the community to run on their own compute clusters or in the cloud at GNPS^7^. To help users learn how to write and perform MassQL queries, we have created documentation (<u>Link</u>), instructional videos (<u>Link</u>), and an interactive MassQL sandbox (https://msql.ucsd.edu/). The MassQL sandbox enables users to interactively write and apply queries on demonstration data, including public MS/MS spectral libraries. As the research community is global, each MassQL query within the sandbox includes an automated translation into English, Portuguese, Spanish, German, French, Chinese, Japanese, Korean, and Russian, which can be included in manuscripts and grants to ensure reproducibility. From the MassQL sandbox, users can implement their desired search parameters and with a single click, apply the query to their own data in GNPS.

Community members have written and applied MassQL and contributed to a wiki-like community compendium using 35 applications of MassQL (<u>Link</u>, **Fig 1d**). The MassQL query compendium will function as an app store to provide a centralized location for MassQL query deposition. The compendium provides an opportunity for users to reuse successful MassQL queries to search for the same or similar classes of compounds in other MS datasets. Uniquely, MassQL is scalable and able to query individual files and across hundreds of thousands of data files from thousands of public projects in multiple repositories, including MassIVE/GNPS^7^, Metabolomics Workbench^10^, and MetaboLights^11^. This capability has thus far aided in the discovery of uncharacterized analogs of different chemical classes in a wide variety of fields, including environmental chemistry, (**SI Note 1.1.1 - Unexpected Organophosphate Compounds in the Environment**), bioinorganic chemistry (**SI Note 3.3.1 - Discovering Iron Binding Molecules from Fungi**), and natural product discovery (**SI Note 3.5.1 - Pentabrominated Natural Products**). Due to the unique scalability of MassQL, it is possible to focus queries towards a specific structural class across an entire repository of data. To enable users to efficiently explore the potentially large chemical diversity present in all public data, MassQL’s output is interoperable with existing software tools such as molecular networking. For the first time, it is possible to use a straightforward language as a filter in conjunction with molecular networking to focus on a structural class across an entire data set or repository and visualize that class’ full chemical diversity (**SI Notes 1.1.2, 1.5, 1.7, 1.9, 1.12, 3.8)**. MassQL derives strength from the users in the community as an open-source, flexible, shareable, instrument agnostic, and scalable data analysis tool. We envision the flexibility of MassQL and further development of the language and ecosystem over time to meet the scientific community’s growing needs in mining MS data, a nearly-indispensable chemical analysis solution for science.

## Code Availability

Reference Engine Implementation (Python), language formal grammar, GNPS Workflow, NextFlow Workflow, and interactive web interface can be found here: https://github.com/mwang87/MassQueryLanguage

Also available in Pypi: https://pypi.org/project/massql/

R API can be found here: https://github.com/rformassspectrometry/SpectraQL

Mzmine: https://github.com/mzmine/mzmine3

OpenMS: https://pyopenms.readthedocs.io/en/latest/massql.html

MS-DIAL 5: http://prime.psc.riken.jp/compms/index.html

UniDec: https://github.com/michaelmarty/UniDec

Language Documentation: https://mwang87.github.io/MassQueryLanguage_Documentation/

## Acknowledgements

We thank Alan Leung for initial discussions and guidance on language design. This research was supported in part by the BBSRC-NSF award 2152526, the Intramural Research Program of National Institute of Environmental Health Sciences of the NIH (ES103363-01, Jarmusch), (ES030158, Fiehn), the National Institute of General Medical Sciences of the NIH (R01 GM125943, Sanchez; R01 GM107550, Gerwick/Dorrestein; R35 GM128690, Phelan), the National Institute of Allergy and Infectious Diseases of the NIH (R21AI156669, McCall), (R15AI137996, Stevens), National Science Foundation (2128044, Sanchez), (CHE-1845230, Marty), NSF CAREER Award (2047235 Garg), the Burroughs Wellcome Fund (1021280, McCall), Fundação de Amparo à Pesquisa do Estado de São Paulo (2018/24865-4), Fundação de Amparo à Pesquisa do Estado do Rio de Janeiro (E-26/201.260/2021, Borges; E-26/211.314/2019, Chagas), Biological Sciences Scholars Program at the University of Michigan (Kersten), the National Research Foundation of Korea (NRF-2020R1C1C1004046, Kang), the German Research Foundation (EXC 2124, Petras), the Swedish Research Council (VR 2020-04707, Globisch), Fund for Financing Science and Innovation Support under the Ministry of Innovative Development of the Republic of Uzbekistan (Kakhkhorov), the German Research Foundation (DFG) TRR 261 (project 398967434, Walter), FOR2372 (project 290827466, Crüsemann) the German Ministry for Education and Research (de.NBI, BMBF FKZ031 A 534A) (and EPIC-XS, project number 823839, funded by the Horizon 2020 programme of the European Union (Kohlbacher, Sachsenberg), the Czech Science Foundation (21-11563M, Pluskal), U.S. Department of Energy Joint Genome Institute (https://ror.org/04xm1d337; a DOE Office of Science User Facility, is supported by the Office of Science of the U.S. Department of Energy operated under Contract No. DE-AC02-05CH11231 (Northen and Bowen) and Subcontract NO. 7601660, Wang. JSPS KAKENHI (21K18216, H.T.), the National Cancer Center Research and Development Fund (2020-A-9, H.T.), JST ERATO Grant (JPMJER2101, H.T.), AMED Japan Program for Infectious Diseases Research and Infrastructure (21wm0325036h0001, H.T.), JST National Bioscience Database Center (NBDC, H.T.), the Novo Nordisk Foundation, Denmark (NNF20CC0035580, NNF16OC0021746, Weber), University of Michigan Biological Science Initiative (UM-BSI, A.T.), Betty and Gordon Moore Foundation (ATA)

## Author Contributions

MW conceived the project. MW and AKJ designed the language. MW, RS, JR, AV, HT, MTM, and TTS developed the software. MW, PCD supervised the development of the project. DP, ATA, AKJ, AB, MTM, WB, NG, VVP and PCD provided feedback. AB, JJJvdH, KBK, LIM, RS, MTM, SAK, TP, WB, RMB, FOC, QX tested the software. AB, JJJvdH, KBK, KL, LIM, WC, TD, TP, MSPC, DG, ACM, NG, MC, MJB, RMB, FOC, OGM, AT, EEK, TW, MMAA, PDB, SH, QX, ATA contributed a use case. MW, ATA, DP wrote the documentation. MW, ATA, AKJ, VVP, PCD wrote the manuscript. All authors edited and approved of the manuscript.

