## Supplemental Information for "A Universal Language for Finding Mass Spectrometry Data Patterns"

#### Table of Contents


**1 MassQL Query using MS/MS Spectral Information** 3

1.1.1 MassQL Query for Organophosphate Compounds in the Environment 3

1.1.2 MassQL Query for Organophosphate Compounds in the Environment at Data Repository Scale 5

1.2 Detection of Fusaricidin Depsipeptides in Microbial Extracts 8

1.3 Impact of Leishmania major Infection on Glycerophosphocholine Lipids 10

1.4 Metabolism of Antibiotic Trimethoprim by Burkholderia cenocepacia 12

1.5 Investigating Cannabidiol Degradation Products using MassQL 14

1.6 Finding Drug Metabolites from Public Fecal Datasets 16

1.7 Using MassQL to Identify New Albicidin Derivatives via Pseudo Precursor Ion, Pseudo Neutral Loss, and Sequence Tag Scan 19

1.8 Investigation of Horizontally Acquired Quorum Signal Synthases from Predatory Myxobacteria 24

1.9 Looking for Potential New Antibiotics by Searching Aminoglycosidic Compounds in Natural Products using MassQL 25

1.10 Gallic acid derivatives as potential anti-inflammatory agents in Plants 27

1.11.1 Finding Lipid Tail Length Analogs with MassQL 29

1.11.2 Finding Lipid Tail Length Analogs with MassQL: Short Tail Variant 31

1.11.3 Finding Lipid Tail Length Analogs with MassQL: Longer Tail Variant 32

1.12 Natural Product Peptidogenomics 34

1.13 Exploring in vitro Liver Metabolism of Xenobiotics using MassQL 37

1.14 MassQL Query for Xenobiotic Conjugation of Malonyl Glucose Conjugates 39

1.15 Sulfatome analysis of urine samples 41

1.16 Data exploration for glycosylated macrolides produced in Actinomycetes 43

1.17 Identification of cyclic peptide analytes in plant metabolomes 45

**2 MassQL Query using MS/MS Spectral Information and Multiple Queries** 51

2.1 Tracking The Biosynthesis of Isotopically Labeled Phenylpropanoids 51

2.2.1 Discovery of Novel Piperamides in Piperaceae plants using MassQL 54

2.2.2 Discovery of Novel Piperamides in Piperaceae plants using MassQL: Diagnostic Ions Refined 56
2.3 Searching for Glycoalkaloids from Solanum species using MassQL 58
2.4 Search for Sources of Compounds of Interest for SARS-CoV-2 61
2.5 Homoserine Lactone 63
2.6 Screening Chemical Diversity of Peptide Natural Products 66
2.7 MassQL to Assess Mass2Motif Substructure Patterns and Annotate Mass2Motif Substructure Patterns: Acylcarnitines as a Case Study 68
2.8 - MassQL for the Distinction of C-hexoglycosides, O-hexoglycosides and Pentoglycosides 75

**3 MassQL Query using MS/MS Spectral Information and MS Isotope Pattern** 77
3.1 Manual Exploration versus MassQL for Processing Untargeted Metabolomics Data: Data Exploration for Sulfur-containing Perfluorinated Compounds in Human Plasma (NIST Standard Reference Material 1950) 77
3.2.1 Exploring Drug Metabolism in Human Plasma via MassQL 82
3.2.2 Exploring Drug Metabolism in Human Plasma via MassQL: Product and Neutral Loss 84
3.3.1 Discovering Iron Binding Molecules from Fungi: 54Fe peak 85
3.3.2 Discovering Iron Binding Molecules from Fungi: 54Fe peak, 13C peak, and apo peak 87
3.4 Discovering Putative Iron Binding Molecules in Repository Search 89
3.5.1 Polybrominated analogs and putative biosynthetic precursors of the “eagle killer toxin”, aetokthonotoxin 90
3.5.2 Repository Scale MassQL Search to find Pentabrominated Natural Products 98
3.6.1 Chlorinated Compounds in Lichen Thalli Extracts: Monochlorinated Compounds 103
3.6.3 Chlorinated Compounds in Lichen Thalli Extracts: Trichlorinated Compounds 106
3.7.1 Halogenated Compounds: Chloride 108
3.8 MassQL Query for Stable Isotope Labeling and Compound Specific Fragment Analysis for Relevant Xenobiotic Metabolites 111
3.9 Application of MassQL for Mass Defect Filtering - Searching for Acylphloroglucinolated Catechins from Agrimonia pilosa 113

**4 MassQL Query and Ion Mobility - Mass Spectrometry Data** 116
4.1 MassQL to Search for Perfluoroalkyl and Polyfluoroalkyl Substances (PFAS) with Ion Mobility in Parallel Accumulation and Serial Fragmentation (PASEF) Data 116
4.2 Detecting Fungal Metabolites of Interest in Microbial Extracts from Isobars Separated by Trapped Ion Mobility Spectrometry (TIMS) 120

**5 MassQL Query and Gas Chromatography Mass Spectrometry** 123
5.1 Detection of fatty acid ethyl esters (FAEEs), biomarkers for excessive alcohol consumption, using MassQL to query GC-Orbitrap-EI data 123

### 1 MassQL Query using MS/MS Spectral Information

##### 1.1.1 MassQL Query for Organophosphate Compounds in the Environment

**Author/s: Nina Zhao**

| |
|---|
| **MassQL Query**<br>`QUERY scansum(MS2DATA) WHERE`<br>`MS2PROD=98.9847:TOLERANCEPPM=50:INTENSITYPERCENT=50 FILTER MS2PROD=98.9847` |
| **MassQL Translation**<br>`Returning the summed scan information on MS2.`<br>`The following conditions are applied to find scans in the mass spec data.`<br>`Finding MS2 peak at m/z 98.9847 with a 50.0 PPM tolerance and a minimum percent intensity relative to base peak of 50.0%.`<br>`Finding MS2 peak at m/z 98.9847.` |
| **MassQL Query Link**<br>https://proteomics2.ucsd.edu/ProteoSAFe/status.jsp?task=e3504aff6ee14a619b19a3ffa3625fb6 |
| **Additional Data Analysis**<br>**GNPS MS/MS Library Search**<br>https://gnps.ucsd.edu/ProteoSAFe/status.jsp?task=1b0382c15423412cba012ed4162d1ebf |
| **Data Availability**<br>MSV000087187 |

Organophosphate esters (OPEs) are widely used as flame retardants and plasticizers in consumer and industrial products. The environmental hazard of OPEs is of concern due to their increasing usage as the replacements of traditional brominated flame retardants and their toxicity effects, which include fecundity decrease, thyroid endocrine disruption, and developmental toxicity in fish. Most of the environmental investigations on OPEs employ targeted analysis with mass spectrometry, but recent studies utilizing non targeted mass spectrometry have revealed novel OPEs. Therefore, screening strategies for OPEs in environmental samples are needed.

OPEs have the general structure $O{=}P(OR)_3$, a central phosphate group with three alkyl or aromatic moieties. Previous studies have reported the phosphate ion ($H_4O_4P^+$, *m/z* 98.9842) as the diagnostic fragment of this class of chemicals. We previously identified several OPEs in the marine water samples from Puget Sound. Here, we used MassQL to search for the phosphate ion fragment in the same dataset.

MassQL returned 771 spectra hits; among them, ~22% (182, belonging to 4 major precursors *m/z*) were noise spectra which required further filters on percent of total intensity to be ruled out. The rest of the spectra hits (589) belong to ~60 unique molecular features and show a true fragment of *m/z* 98.98. A subsequent library search identified four OPEs, tributyl phosphate, tris(chloroethyl) phosphate, tris(chloroisopropyl) phosphate, and tirs(butoxyethyl) phosphate, explaining ~56% (331) of the non-noise spectra hits. Tributyl phosphate was not identified in our original publication based on traditional fold-change non targeted analysis. This example demonstrated the possibility to screen for environmental contaminants in MS data without a predefined suspect list.

##### 1.1.2 MassQL Query for Organophosphate Compounds in the Environment at Data Repository Scale

**Author/s: Nina Zhao**

| |
|---|
| **MassQL Query**<br>`QUERY scansum(MS2DATA) WHERE`<br>`MS2PROD=98.9847:TOLERANCEPPM=50:INTENSITYPERCENT=50 FILTER MS2PROD=98.9847` |
| **MassQL Translation**<br>`Returning the summed scan information on MS2.`<br>`The following conditions are applied to find scans in the mass spec data.`<br>`Finding MS2 peak at m/z 98.9847 a 50.0 PPM tolerance and a minimum percent intensity relative to base peak of 50.0%.`<br>`Finding MS2 peak at m/z 98.9847.` |
| **MassQL Query Link (repository-scale)**<br>https://proteomics2.ucsd.edu/ProteoSAFe/result.jsp?task=f5094e83a4f042e88f0d423dcb52b11c&view=extract_results |
| **Additional Data Analysis**<br>**Falcon Cluster, EPS = 0.3**<br>https://proteomics2.ucsd.edu/ProteoSAFe/status.jsp?task=f1a4a4c3496645e8b643f8d41c601d03<br>**GNPS Molecular Networking**<br>https://proteomics2.ucsd.edu/ProteoSAFe/status.jsp?task=cf943d6808ed4d7da441184f531a62c3 |
| **Data Availability**<br>All public data from Q Exactive on GNPS |

To discover novel organophosphate compounds, we ran the phosphate ion MassQL query developed in Use Case 14 over all Q Exactive data on the GNPS repository. The MassQL query returned 338,439 spectra hits in total. Based on a comprehensive list of organophosphate compounds (n = 95) compiled by Ye et al.[1] (2021), only 15% (51,310) of the spectra hits could be explained by these known organophosphate compounds (based on precursor *m/z* match with 20 ppm mass error). Top matches included tris(chloropropyl) phosphate (31,039 matches), tributyl phosphate and isomers (12,967 matches), tri(2-ethylhexyl) phosphate (or structural isomers; 3511 matches), and triethyl phosphate (2171 matches). The remaining 85% matches represent a candidate pool for new organophosphate compounds.

We further analyzed the query results using molecular networking. MS-Cluster, the clustering tool for classical molecular networking on GNPS, failed to handle such a large amount of spectra and returned many repetitive clusters (example: link). We explored a newly developed clustering tool, Falcon[2], which very effectively reduced the number of repetitive clusters. Sending the Falcon output to molecular networking returned 169 spectra families. Mining into these spectra families led to new compound identification. For example, **SI Figure 1.1.2 - 1a** demonstrated a spectra family of organophosphate compounds with alkyl substituents. The *m/z* 267.17 represented tributyl phosphate with dimers at *m/z* 533.34, and the *m/z* 435.36 represented trioctyl phosphate (or structural isomers) with dimers at *m/z* 869.71 and trimers at *m/z* 1305.07. Notably, the group of *m/z* 211.03 matched the molar mass of dibutyl phosphate, which was not discovered by library search. **SI Figure 1.1.2 - 1b** demonstrated a spectra family of chlorinated organophosphate compounds. The m/z 428.89 matched tris(dichloro-propyl) phosphate, and *m/z* 327.01 matched tris(chloro-propryl) phosphate compounds. Such findings highlighted the capacity of MassQL coupled to classical molecular networking to connect related chemicals on a repository scale and to identify new chemicals.

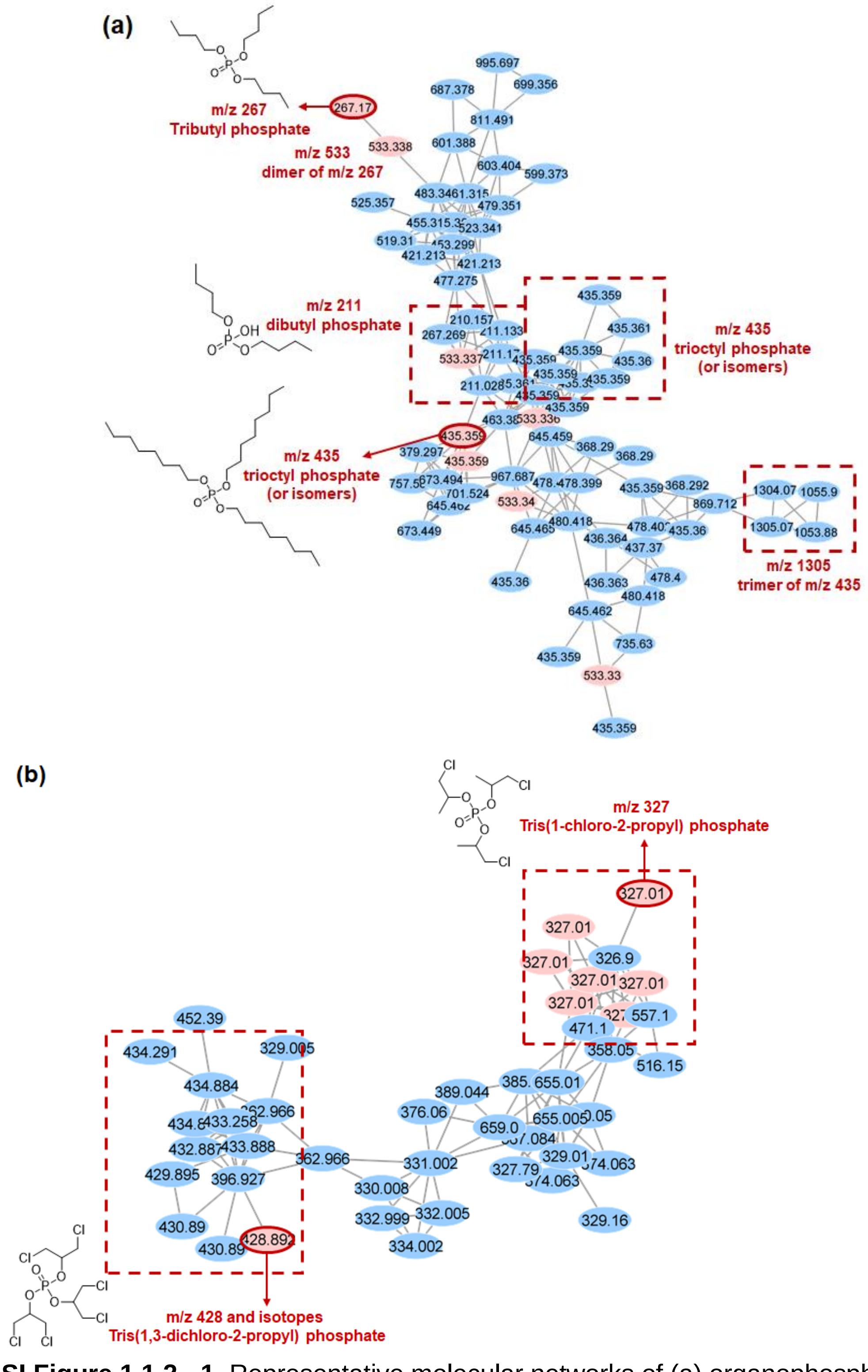


**SI Figure 1.1.2 - 1.** Representative molecular networks of (a) organophosphate compounds with alkyl substituents and (b) with chlorinated substituents identified through MassQL search. Red and blue nodes represented identified and unidentified spectra through GNPS library search, respectively.

#### 1.2 Detection of Fusaricidin Depsipeptides in Microbial Extracts

**Author/s: Osama G. Mohamed and Ashootosh Tripathi**

| |
|---|
| **MassQL Query**<br>`QUERY scaninfo(MS2DATA) WHERE`<br>`MS2NL=255.23: TOLERANCEMZ=0.02 AND MS2PROD=256.23:TOLERANCEMZ=0.02` |
| **MassQL Translation**<br>`Returning the scan information on MS2.`<br>`The following conditions are applied to find scans in the mass spec data.`<br>`Finding MS2 neutral loss peak at m/z 255.23 a 0.02 m/z tolerance.`<br>`Finding MS2 peak at m/z 256.23 a 0.02 m/z tolerance.` |
| **MassQL Query Link**<br>https://proteomics2.ucsd.edu/ProteoSAFe/status.jsp?task=3afe33c186a44bfe945a74d21c06f51e<br>**MassQL Query Link (large scale)**<br>https://proteomics2.ucsd.edu/ProteoSAFe/status.jsp?task=7fc6e39e34f84a159cb171b31ca03b09 |
| **Additional Data Analysis**<br>**GNPS Molecular Networking**<br>https://gnps.ucsd.edu/ProteoSAFe/status.jsp?task=9ef11868efa94bde892d27101bb579a4 |

Fusaricidins are a class of cyclic lipopeptides isolated from *Paenibacillus* sp. Their structures consist of two moieties, a cyclic polypeptide that consists of six amino acids and a guanidino-3-hydroxypentadecanoic acid fatty acid moiety (GHPD). This class of metabolites has shown *in vitro* antimicrobial activity against Gram-positive bacteria and *Fusarium* fungi. Fusaricidins MS2 fragmentation spectra are characterized with the fragment product ion *m/z* 256.2389 (GHPD) and neutral loss of 255.23 (residual cyclopeptide moiety, MH-GHPD) (**SI Figure 1.2 - 1**).

The use of MassQL enabled us to quickly detect and group fusaricidins in a microbial extract. The current resources of University of Michigan Natural Products Discovery Core (UM-NPDC) include a > 50,000 sample natural product extracts (NPE) library collected and curated from across the world, including Costa Rica, Papua New Guinea, Israel, Nepal, Panama, Peru, and the United States. Up to date, ~3000 NPEs have been chemically profiled using UHPLC-qTOF and are still growing. This tool can foster the discovery process of new congeners of interesting metabolites with characteristic tandem MS spectra.

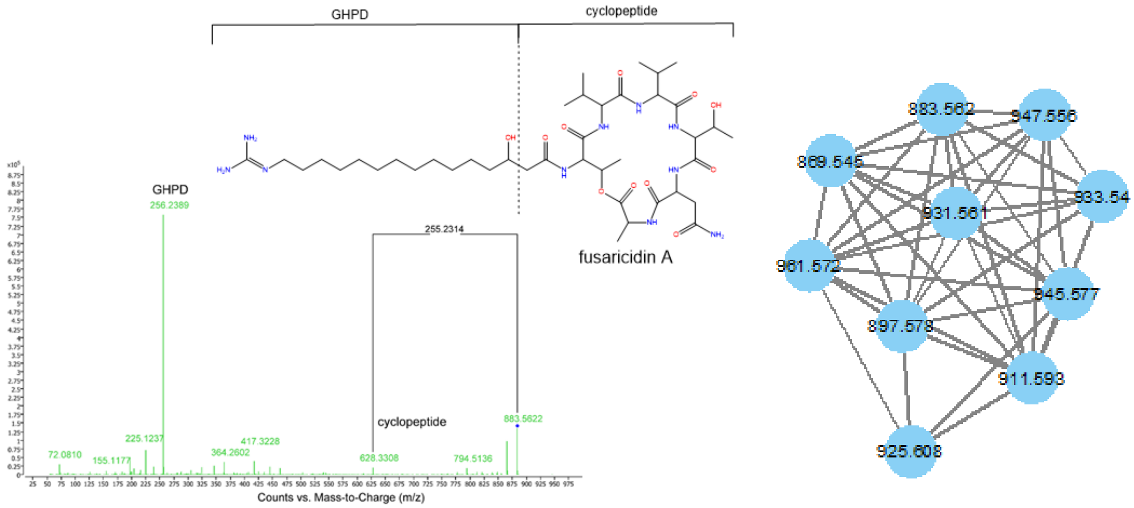


**SI Figure 1.2 - 1.** Experimental MS2 fragmentation of fuscaridin A and resulting molecular network of fuscaridins from MassQL query.

#### 1.3 Impact of *Leishmania major* Infection on Glycerophosphocholine Lipids

**Author/s: Laura-Isobel McCall**

| **MassQL Query** |
| --- |
| `QUERY scaninfo(MS2DATA) WHERE MS2PROD=184.0739:TOLERANCEMZ=0.01:INTENSITYPERCENT=30:INTENSITYVALUE=500 AND MS2PROD=125.0004:TOLERANCEMZ=0.01:INTENSITYPERCENT=10:INTENSITYVALUE=1500 AND MS2PROD=104.1075:TOLERANCEMZ=0.01 AND MS2PROD=86.09697:TOLERANCEMZ=0.01:INTENSITYPERCENT=10:INTENSITYVALUE=2000` |
| **MassQL Translation**<br>Returning the scan information on MS2.<br>The following conditions are applied to find scans in the mass spec data.<br>Finding MS2 peak at m/z 184.0739 a 0.01 m/z tolerance and a minimum percent intensity relative to base peak of 30.0% and a minimum intensity value 500.0.<br>Finding MS2 peak at m/z 125.0004 a 0.01 m/z tolerance and a minimum percent intensity relative to base peak of 10.0% and a minimum intensity value 1500.0.<br>Finding MS2 peak at m/z 104.1075 a 0.01 m/z tolerance.<br>Finding MS2 peak at m/z 86.09697 a 0.01 m/z tolerance and a minimum percent intensity relative to base peak of 10.0% and a minimum intensity value 2000.0. |
| **MassQL Query Link**<br>https://proteomics2.ucsd.edu/ProteoSAFe/status.jsp?task=abb71fc649a945fc950ad7fd684eefeb |
| **Additional Data Analysis**<br>**GNPS Feature-based Molecular Networking**<br>https://gnps.ucsd.edu/ProteoSAFe/status.jsp?task=451754c383de461e9e4abdf6eb3199d2 |
| **Data Availability**<br>MSV000081004 |

Current treatments for neglected parasitic diseases suffer from low efficacy, high rates of adverse effects, and emergence of drug resistance. Identifying metabolic pathways altered by infection *in vivo* may lead to the discovery of new antiparasitic treatment strategies, which are sorely needed. Studies of infection-modulated metabolites have revealed infection-induced metabolic perturbations in multiple members of the glycerophosphocholine family of phospholipids[3,4], necessitating a detailed investigation of this family of metabolites under infection conditions. Glycerophosphocholines have four diagnostic MS2 fragmentation peaks in positive mode: *m/z* 184.0739 (phosphocholine), *m/z* 125.0004 (2,2-Dihydroxy-1,3,2-dioxaphospholan-2-

ium), *m/z* 104.1075 (choline) and *m/z* 86.09697 (N,N,N-Trimethylethenaminium) from the phospholipid head group. We therefore built a MassQL query to search a dataset of *Leishmania major*-infected and uninfected mouse ear tissue.

MassQL query returned all of the glycerophosphocholines that had previously been annotated via molecular networking and manually inspected for the presence of the four MS2 glycerophosphocholine diagnostic peaks[3]. Prior manual analysis had taken >4 h, whereas MassQL run took 2.5 min, followed by ~15 min of results exploration (**SI Figure 1.3 - 1**). In addition, this query returned four more features (*m/z* 518.3235 retention time 203.635 sec, *m/z* 552.3299 retention time 172.307 sec, *m/z* 600.403 retention time 224.992 sec, and *m/z* 768.5895 retention time 366.716 sec). Using LIPIDMAPS[5] and based on precursor mass, we annotate *m/z* 518.3235 as LPC 18:3, *m/z* 552.3299 as PC 19:0, *m/z* 600.403 as LPC 24:4, LPC O-24:5;O or PC O-24:4, and *m/z* 768.5895 as PC O-36:4. These features had been missed in our prior analyses because they were singleton network nodes and had not returned a library match using our prior networking parameters.

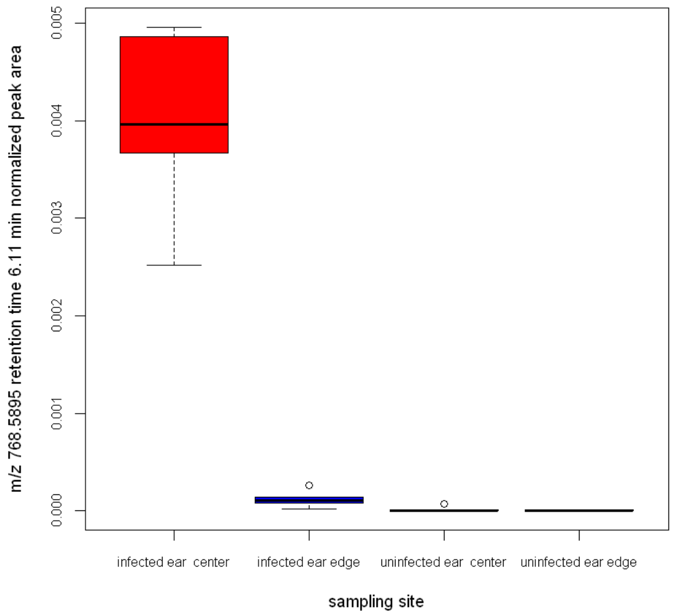


**Figure 1.3 - 1.** Box and whisker plot of normalized peak area for *m/z* 768.5895, putatively identified as PC O-36:4, a glycerophosphocholine result using MassQL. Normalized peak area is elevated by *Leishmania major* infection at lesion sites (infected ear center and infected ear edge) compared to uninfected samples.

#### 1.4 Metabolism of Antibiotic Trimethoprim by *Burkholderia cenocepacia*

**Author/s: Neha Garg and Andrew McAvoy**

| |
|---|
| **MassQL Query**<br>`QUERY scaninfo(MS2DATA) WHERE`<br>`MS2PROD=291.1458 AND MS2PROD=123.0665` |
| **MassQL Translation**<br>`Returning the scan information on MS2.`<br>`The following conditions are applied to find scans in the mass spec data.`<br>`Finding MS2 peak at m/z 291.1458.`<br>`Finding MS2 peak at m/z 123.0665.` |
| **MassQL Query Link**<br>https://proteomics2.ucsd.edu/ProteoSAFe/result.jsp?task=db21c5c316034b09a74668b53828b39b&view=query_results |
| **Additional Data Analysis**<br>**GNPS Feature-based Molecular Networking**<br>https://gnps.ucsd.edu/ProteoSAFe/status.jsp?task=451754c383de461e9e4abdf6eb3199d2 |
| **Data Availability**<br>MSV000084945 |

Trimethoprim when combined with sulfamethoxazole is a bacteriocidal antibiotic used by clinicians to treat bacterial infections, such as those caused by *Burkholderia cepacia* complex (Bcc) bacteria in cystic fibrosis patients. *Burkholderia* spp. are well known for their ability to metabolize xenobiotic compounds. In a previous untargeted metabolomics study, we cultured Bcc bacteria in the presence and absence of trimethoprim. In MS/MS, trimethoprim features a prominent peak with *m/z* 123.0665 generated by cleavage of a carbon-carbon bond of the methylene connecting the two rings, yielding the 2,4 diaminopyrimaidin-5-yl-methylium fragment ion (**SI Figure 1.4 - 1**). With MassQL, we are able to identify spectra containing fragment masses corresponding to trimethoprim (*m/z* 291.1452) and the 2,4 diaminopyrimaidin-5-yl-methylium fragment, allowing us to mine our dataset for metabolomic features that correspond to novel biotransformation products of trimethoprim.

This MassQL query returned a list of metabolites which display an MS2 fragmentation pattern indicative of a trimethoprim substructure, facilitating the identification of trimethoprim biotransformation products. Manual inspection revealed that 33 out of the 47 candidate features truly contained a trimethoprim substructure, which can undoubtedly be improved by optimizing the query. Previously, we relied on MS2LDA[6] to find compounds containing a trimethoprim substructure, which involves extracting Mass2Motifs from data, manually inspecting the Mass2Motifs found in trimethoprim, and searching for all other molecules which contain relevant Mass2Motifs. MassQL provided additional flexibility to refine the Mass2Motif pattern. Notably, the MassQL query correctly identified a molecule with *m/z* 549.129 as containing a trimethoprim substructure, which was not recognized by MS2LDA analysis. Although manual inspection of raw data should always be performed to verify *in silico* predictions, MassQL has the potential to greatly streamline metabolomic analysis in a manner that can be creatively applied based on the unique analysis needs of a particular dataset.

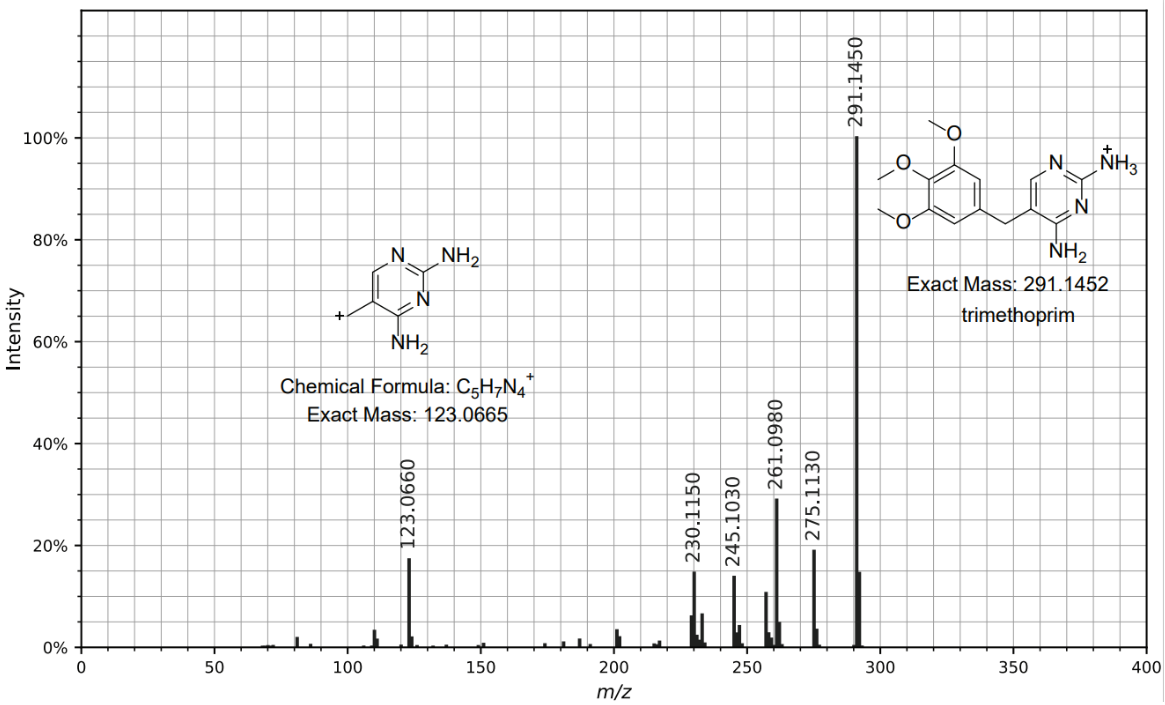


**SI Figure 1.4 - 1.** Experimental MS2 spectrum of trimethoprim, with the characteristic 2,4 diaminopyrimaidin-5-yl-methylium fragment ion (*m/z* 123.0665) indicated.

#### 1.5 Investigating Cannabidiol Degradation Products using MassQL

**Author/s: Matthew J. Bertin and Riley D. Kirk**

| |
|---|
| **MassQL Query**<br>`QUERY scaninfo(MS2DATA) WHERE`<br>`MS2PROD=259.5:TOLERANCEMZ=0.5` |
| **MassQL Translation**<br>`Returning the scan information on MS2.`<br>`The following conditions are applied to find scans in the mass spec data.`<br>`Finding MS2 peak at m/z 259.5 a 0.5 m/z tolerance.` |
| **MassQL Query Link (day 1)**<br>https://proteomics2.ucsd.edu/ProteoSAFe/result.jsp?task=d9c7673620b54e58a15f967334a5024d&view=query_results<br>**MassQL Query Link (day 4)**<br>https://proteomics2.ucsd.edu/ProteoSAFe/result.jsp?task=c80fcbe42f7048678c66ca379487f53a&view=query_results |
| **Additional Data Analysis**<br>**GNPS Molecular Networking**<br>Day 1:<br>https://gnps.ucsd.edu/ProteoSAFe/status.jsp?task=432b9dff5ef143b69b719ef42a41d9d2<br>Day 4:<br>https://gnps.ucsd.edu/ProteoSAFe/status.jsp?task=bd728f3e31ac447384118dc3003cba01 |

Cannabidiol (CBD) is a major phytocannabinoid found in Cannabis plants. The use of CBD continues to gain interest in the pharmaceutical, nutraceutical, and even the cosmeceutical industries. Detection and annotation of cannabidiol degradation products remains important for understanding the stability and quality of the now numerous CBD products available to the public. This understanding is especially critical with recent research demonstrating the biological activity (inhibition of topoisomerase IIα and β) of an oxidized cannabidiol quinone (CBDQ)[7]. We analyzed purified CBD at two time periods (t = 0 d and t = 4 d) following exposure to natural light, and the clear solution on day 0 began to take on a pale yellow color by day 4. LC-MS/MS data were acquired in data-dependent mode using a Thermo Scientific LTQ XL. We utilized mass query language to identify product ion *m/z* 259, a prominent fragment ion from cannabinoid precursors. We subsequently subjected each query analysis to molecular networking and we examined the changes in chemical space over time. At day 0, eight unique precursor *m/z* values were identified with the *m/z* 259 product ion, while at day 4, over fifty unique precursor *m/z* values were identified including a putative identification of CBD quinone (*m/z* 329)[8], and many other likely degradation products (**SI Figure 1.5 - 1**). This approach shows an effective means to mine mass spectrometry data to identify potential CBD degradation molecules for quality control as more and more consumers utilize these products.

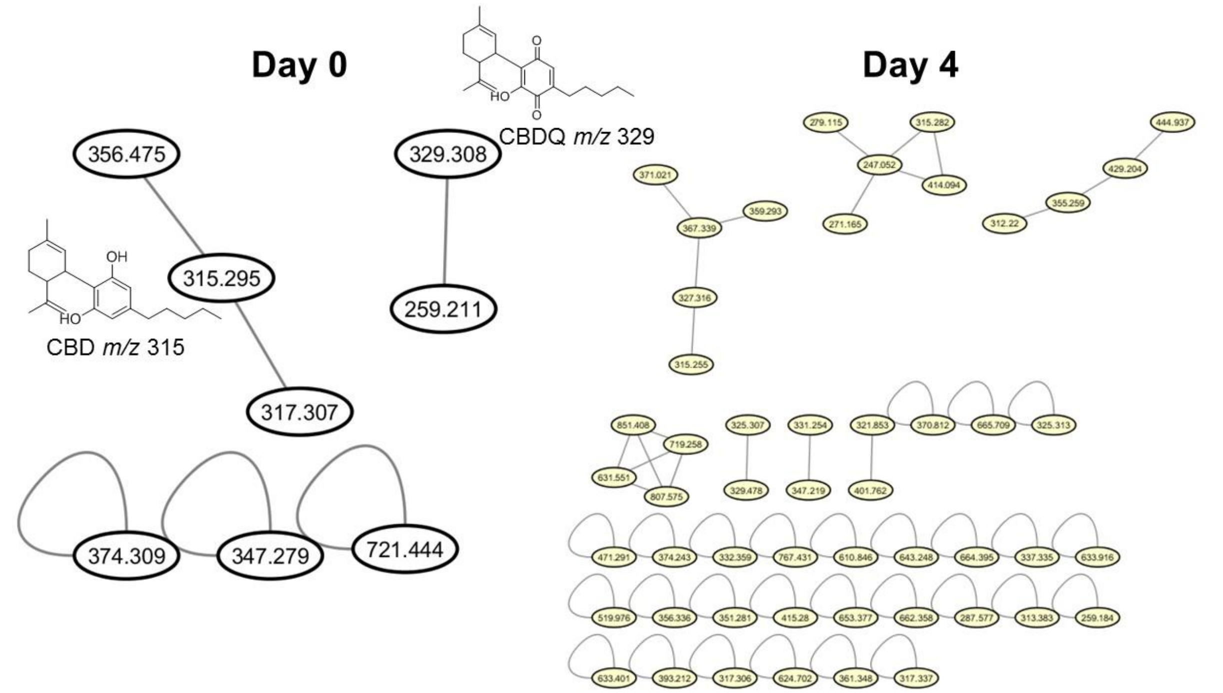


**SI Figure 1.5 - 1**. LC-MS/MS-based molecular networks following query analysis (compounds with product ion *m/z* 259) of CBD standard solution in methanol at both day 0 and day 4. Structures of CBD (*m/z* 315) and CBDQ (*m/z* 329) are shown.

#### 1.6 Finding Drug Metabolites from Public Fecal Datasets

**Author/s: Kyo Bin Kang**

| |
|---|
| **MassQL Query**<br>`QUERY scaninfo(MS2DATA) WHERE MS2PROD=311.1508:TOLERANCEMZ=0.0025:INTENSITYPERCENT>5` |
| **MassQL Translation**<br>Returning the scan information on MS2.<br>The following conditions are applied to find scans in the mass spec data.<br>Finding MS2 peak at m/z 311.1508 a 0.0025 m/z tolerance and a minimum percent intensity relative to base peak of 5.0%. |
| **MassQL Query Link**<br>https://proteomics2.ucsd.edu/ProteoSAFe/status.jsp?task=45b72be5b8e54d0d92959ec57f4f4558 |
| **Additional Data Analysis**<br>N/A |
| **Data Availability**<br>MSV000080673 and MSV000082221 |

In a previous study, we identified 68 putative metabolites of sildenafil by applying molecular networking analysis on LC-MS/MS data of *in vitro* microsomal biotransformation experiments[9]. The MASST search of these spectra revealed that at least four public data (all the data were human fecal extracts) contained intact sildenafil or its major metabolite of *m/z* 449.1971. However, other metabolites were not found in the MASST analysis, and our further investigation revealed that the matched data actually contained the other metabolites but they were not detected due to the low cosine similarity, which was mainly caused by significantly different ion intensities of fragment ions. Here, we queried a common fragment of sildenafil metabolites, *m/z* 311.1508, and this query revealed the occurrence of other sildenafil metabolites in four matched data (**SI Figure 1.6 - 1**).

We found 150 MS/MS scans containing a fragment ion of *m/z* 311.1508 from four public fecal extract data files. Some of them were identified as sildenafil metabolites in our previous study, by comparing their spectra with the experimental spectra acquired in our *in vitro* work. In the previous study, we already discussed that these metabolites were not found in the MASST search, because they showed low cosine similarity to the experimental spectra due to the different relative intensities between fragments. Thus, we found them from the public data by visualizing extracted ion chromatograms (EICs) of identified metabolites and comparing *m/z* values of major fragment ions manually. In this case, we successfully found drug metabolites with the characteristic fragment ion using MassQL, which means we can find out drug metabolites from public dataset more easily even when their MS/MS spectra show low cosine similarity to our reference spectra (**SI Figure 1.6 - 2**).

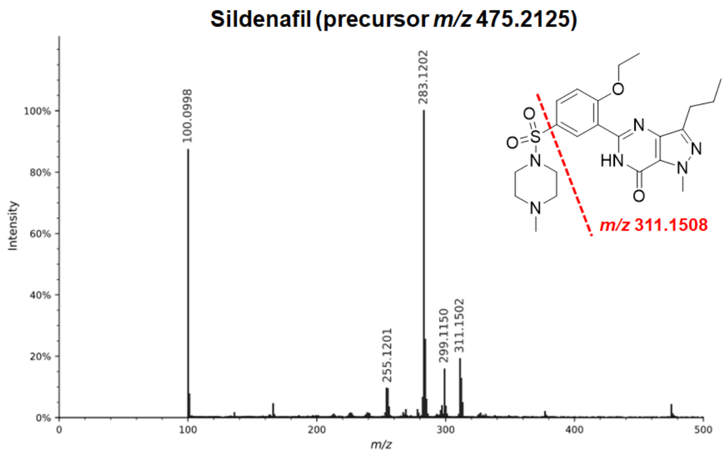


**SI Figure 1.6 - 1**. MS/MS spectrum for Sildenafil with distinctive fragmentation (*m/z* 311.1508) indicated.

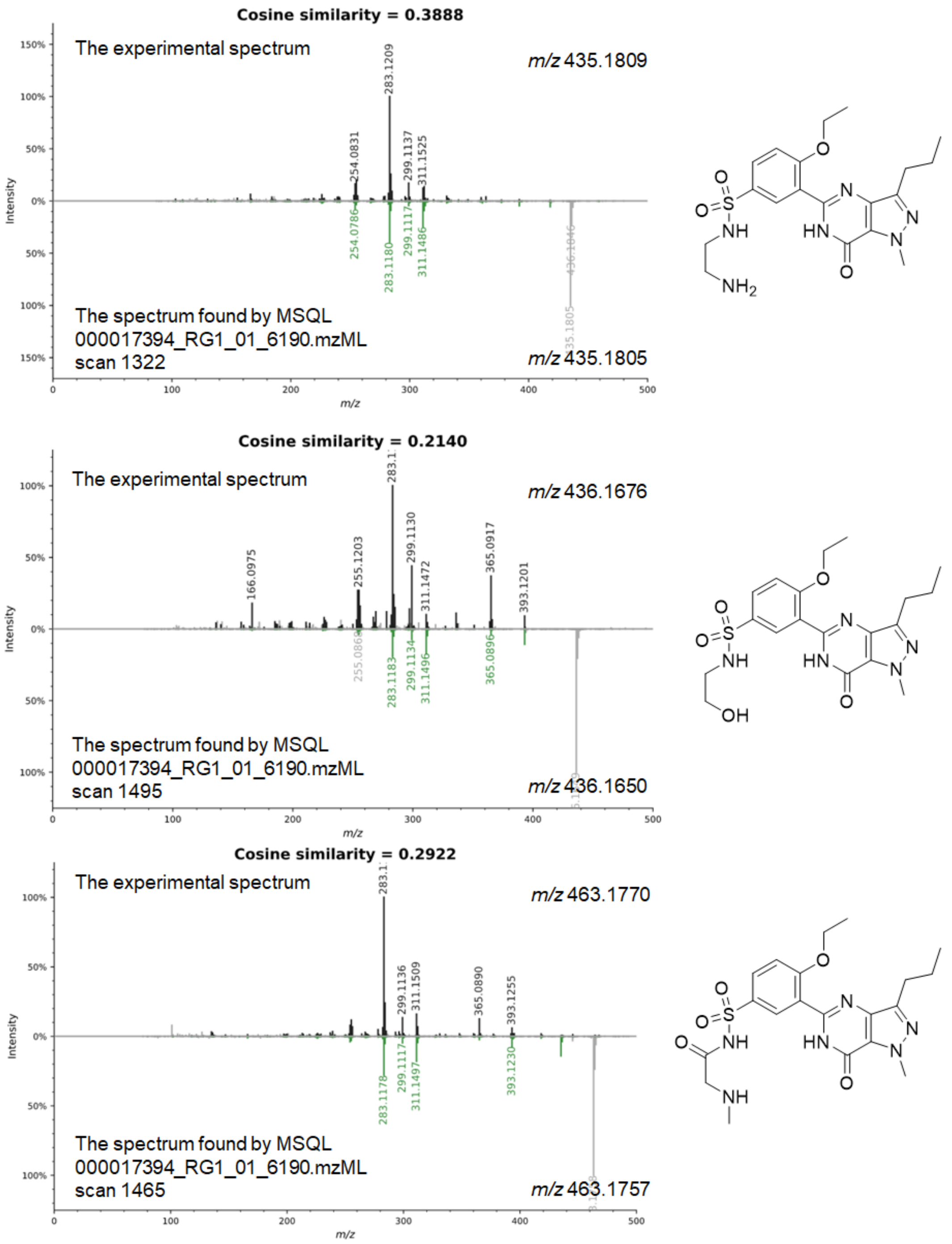


**SI Figure 1.6 - 2.** The query found sildenafil metabolites containing a fragment ion of *m/z* 311.1508 from the public datasets. Mirror plots of found spectra and experimental spectra from the *in vitro* sildenafil metabolism are shown together with suggested metabolite structures.

#### 1.7 Using MassQL to Identify New Albicidin Derivatives via Pseudo Precursor Ion, Pseudo Neutral Loss, and Sequence Tag Scan

**Author/s: Daniel Petras**

Albicidins are a family of non-ribosomal peptide poly-ketide hybrid molecules (NRP-PK) and very potent antimicrobials against a broad range of gram positive and gram negative bacteria[10]. Due to their NRP core, albicidins produce a series of characteristic b and y ions. In order to mine tandem mass spectrometry data from extracts from *Xanthomonas albilineans*, the known producer of albicin, we used MassQL to identify spectra that contain the characteristic fragment masses *m/z* 468.2 and *m/z* 660.2.

**MassQL Query**

```
QUERY scaninfo(MS2DATA) WHERE
MS2PROD=660.2:TOLERANCEMZ=0.1:INTENSITYPERCENT=1 AND
MS2PROD=468.2:TOLERANCEMZ=0.1:INTENSITYPERCENT=1
```

**MassQL Translation**

```
Returning the scan information on MS2.
The following conditions are applied to find scans in the mass spec data.
Finding MS2 peak at m/z 660.2 a 0.1 m/z tolerance and a minimum percent
intensity relative to base peak of 1.0%.
Finding MS2 peak at m/z 468.2 a 0.1 m/z tolerance and a minimum percent
intensity relative to base peak of 1.0%.
```

**MassQL Query Link**

https://proteomics2.ucsd.edu/ProteoSAFe/status.jsp?task=fbde4479225f444f9bd9fd6eef40b507

**Additional Data Analysis**

n/a

**Data Availability**

MSV000080427

With this query we identified 118 scans that fulfilled our criteria by means of containing product ions with *m/z* = 468.2 and 660.2 with a minimum relative intensity of 1%. The most abundant spectra were two MS/MS spectra with a precursor mass of 843.26, the mass of albicidin (total 46 scans). In addition to the main albicidin, our MassQL search identified spectra from 55 other nominal precursor masses and thus potential derivatives. The most frequent masses were *m/z* 875.28 which correspond to known derivative and *m/z* 845.27 which has a double bond less than the main Albcidin. As the b and y ions and thus the product ion masses could be shifted if modification at the N or C terminus are present, not all derivatives will be identified by the pseudo precursor ion scan. Thus, we next used MassQL to search for mass spectra that contain a series of neutral losses from binding blocks typical for albicidins, such an ortho-meta double substituted pABA with the neutral masses. As the double substituted pABA residue (183.096 Da) is located at the C terminus, we would expect a modification of the N terminus to shift both the precursor and the B fragments by the mass of the modification and thus query with following code.

| **MassQL Query**<br>`QUERY scaninfo(MS2DATA) WHERE`<br>`MS2NL=183.096:TOLERANCEMZ=0.1:INTENSITYPERCENT=5` |
|---|
| **MassQL Translation**<br>`Returning the scan information on MS2.`<br>`The following conditions are applied to find scans in the mass spec data.`<br>`Finding MS2 neutral loss peak at m/z 183.096 a 0.1 m/z tolerance and a minimum percent intensity relative to base peak of 5.0%.` |
| **MassQL Query Link**<br>https://proteomics2.ucsd.edu/ProteoSAFe/status.jsp?task=e2d62adc469f442581aaf69132989613 |
| **Additional Data Analysis**<br>n/a |
| **Data Availability**<br>MSV000080427 |

This query identified successfully 72 spectra with the precursor mass 843.27, corresponding to the main albicidin derivative. The higher number in comparison to the results from the precursor scan, could be explained by higher robustness towards mass deviation, as the query measures here relative mass loss between the precursor and a given fragment ion.
In the case of albicidin, several analogs with modification on both N and C terminus are known. To identify these analogs, we would ideally only look at internal neutral losses within the peptide chain. For such an experiment, which is not possible on conventional tandem mass spectrometers, one would need to run an offset for the building block specific delta between two product ions. However with DDA MS/MS data and MassQL, we can simply search for spectra that contain a primary product ions and a second product ion with a defined offset form the first one, according to the building block, in our case pABA or the double substituted pABA.
In the case of albicidin, X could be the b4 ion with *m/z* 495.3 and as X+164.9 the b5 fragment with *m/z* 660.2.

In order to expand our search to peptides that contain a amino acid at any position of the peptide chain, we formulated a new query that uses a variable delta ( e.g. F the delta mass of the double substituted pABA, 164.9 Da) mass pair which we called sequence tag search.

| **MassQL Query** |
|---|

| |
|---|
| `QUERY scaninfo(MS2DATA) WHERE MS2PROD=X:TOLERANCEMZ=0.1:INTENSITYPERCENT=5 AND MS2PROD=X+164.9:TOLERANCEMZ=0.1:INTENSITYPERCENT=5` |
| **MassQL Translation**<br>Returning the scan information on MS2.<br>The following conditions are applied to find scans in the mass spec data.<br>Finding MS2 peak at m/z X a 0.1 m/z tolerance and a minimum percent intensity relative to base peak of 5.0%.<br>Finding MS2 peak at m/z X+164.9 a 0.1 m/z tolerance and a minimum percent intensity relative to base peak of 5.0%. |
| **MassQL Query Link**<br>https://proteomics2.ucsd.edu/ProteoSAFe/status.jsp?task=f67251c63a3c41c0a6b47cdae029360d |
| **Additional Data Analysis**<br>n/a |
| **Data Availability**<br>MSV000080427 |

With this query, 4201 spectra were identified. Among them are the main Albicidin and several other known derivatives that contain a double substitute pABA.

Similar to the query before but focusing on the building block pABA we formulated the following query with the delta mass 119.1 Da. In this case, the y2 ion with *m/z* 349.2 of albicdin was picked as X and the y3 ion with *m/z* 468.3 was matched as X+119.

| |
|---|
| **MassQL Query**<br>`QUERY scaninfo(MS2DATA) WHERE MS2PROD=X:TOLERANCEMZ=0.1:INTENSITYPERCENT=5 AND MS2PROD=X+119.1:TOLERANCEMZ=0.1:INTENSITYPERCENT=5` |
| **MassQL Translation**<br>Returning the scan information on MS2.<br>The following conditions are applied to find scans in the mass spec data.<br>Finding MS2 peak at m/z X a 0.1 m/z tolerance and a minimum percent intensity relative to base peak of 5.0%.<br>Finding MS2 peak at m/z X+119.1 a 0.1 m/z tolerance and a minimum percent intensity relative to base peak of 5.0%. |
| **MassQL Query Link**<br>https://proteomics2.ucsd.edu/ProteoSAFe/status.jsp?task=f7026cb8426e4eca8d513f19e271a6ea |

| |
|---|
| **Additional Data Analysis**<br>n/a |
| **Data Availability**<br>MSV000080427 |

In total, 5,323 spectra fulfilled the requirements of the query, and in addition to the main Albicidin, several other derivatives that contain pABA were identified. Overall both sequence tag searches reveal a larger number of non specific spectra due to the wider search space. To increase specificity for the search and to yield a full sequence tag of multiple amino acids in a defined order, one can simply combine the two queries by adding the first delta to the second through the AND logic, in our case for two subsequent double substitute pABA moieties and concatenate the commands with an AND logic and formulate the following query:

**MassQL Query**

```
QUERY scaninfo(MS2DATA) WHERE MS2PROD=X:TOLERANCEMZ=0.1:INTENSITYPERCENT=5
AND MS2PROD=X+164.9:TOLERANCEMZ=0.1:INTENSITYPERCENT=5 AND
MS2PROD=X+329.8:TOLERANCEMZ=0.1:INTENSITYPERCENT=5
```

**MassQL Translation**

```
Returning the scan information on MS2.
The following conditions are applied to find scans in the mass spec data.
Finding MS2 peak at m/z X a 0.1 m/z tolerance and a minimum percent
intensity relative to base peak of 5.0%.
Finding MS2 peak at m/z X+164.9 a 0.1 m/z tolerance and a minimum percent
intensity relative to base peak of 5.0%.
Finding MS2 peak at m/z X+329.8 a 0.1 m/z tolerance and a minimum percent
intensity relative to base peak of 5.0%.
```

**MassQL Query Link**

https://proteomics2.ucsd.edu/ProteoSAFe/status.jsp?task=d7c4f1981ba14d31b02773ec34d796f1

**Additional Data Analysis**

https://gnps.ucsd.edu/ProteoSAFe/status.jsp?task=3f3244a99dd646eea93884423b0b80bc

**Data Availability**

MSV000080427

The results show that fewer spectra (2,910) where retrieved; however, among them were multiple albicidins with the given b ion series, e.g. the b4 ion with *m/z* 495.3 and as X+164.9 the b5 ion with *m/z* 660.2 and the precursor with water loss at *m/z* 825.3 as the X+329.8 (2 x 164.9), as indicated in Figure 1.7 a. To visualize the results and order and collapse redundant spectra, we generated a molecular network of the MassQL output. Focusing on the precursor mass of Albicidin, we could quickly identify a network of albicindin analogs (Figure 1.7 b) that contained several known (Figure 1.7 c) and new albicidin derivatives. These results shows how increasing the number of delta masses within a sequence tag increase the specificity of the query and can be applied to identify specific amino acid sequences containing natural products, especially non-ribosomal or ribosomal peptides with know biosynthesis clusters / precursor peptides.

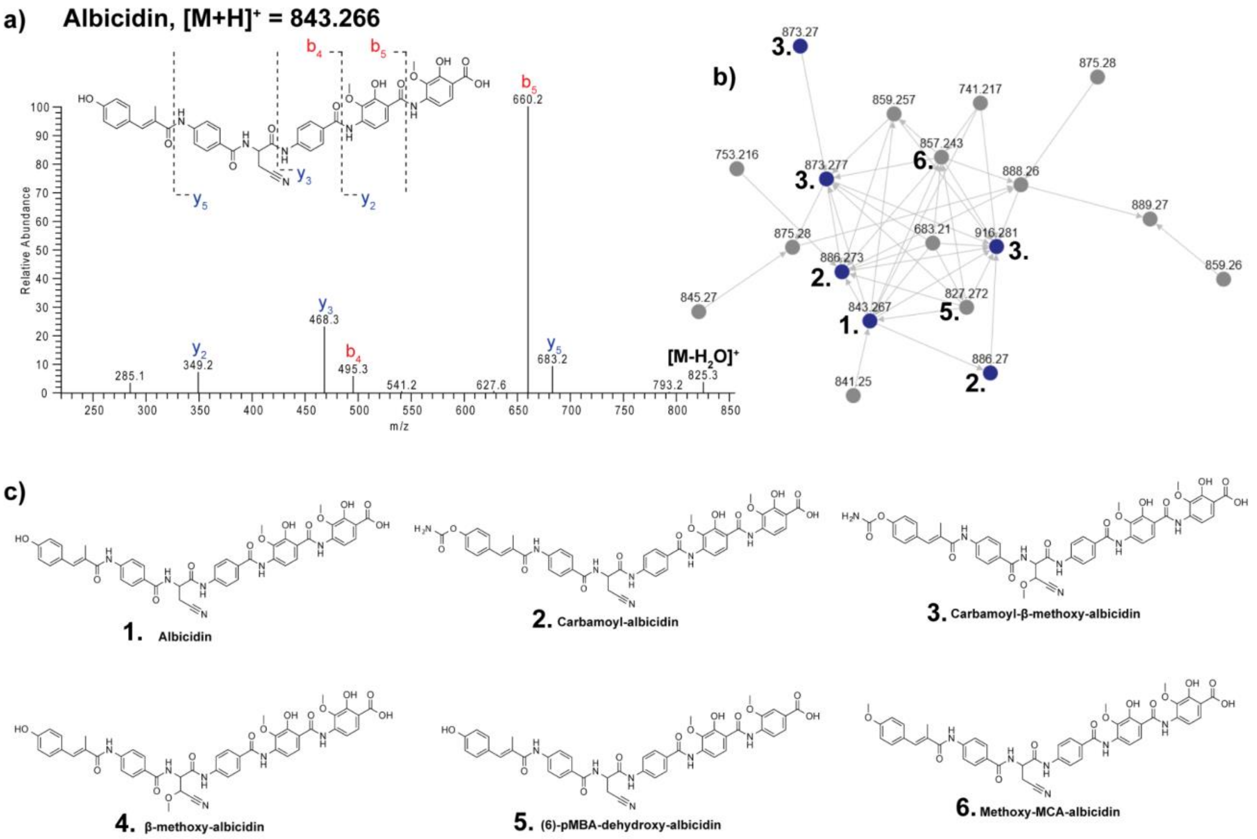


**SI Figure 1.7**: a) Example MS/MS of Albicidin MS/MS fragmentation. GNPS Dashboard link to raw data [Link](). b) Molecular Network of query spectra from Sequence Tag Search. c) Observed known Albicindin derivatives.

#### 1.8 Investigation of Horizontally Acquired Quorum Signal Synthases from Predatory Myxobacteria

**Author/s: Cole Stevens**

| **MassQL Query**<br>`QUERY scaninfo(MS2DATA) WHERE MS2PROD=102.1:TOLERANCEMZ=0.1 AND MS2PROD=74.1:TOLERANCEMZ=0.1` |
|---|
| **MassQL Translation**<br>`Returning the scan information on MS2.`<br>`The following conditions are applied to find scans in the mass spec data.`<br>`Finding MS2 peak at m/z 102.1 a 0.1 m/z tolerance.`<br>`Finding MS2 peak at m/z 74.1 a 0.1 m/z tolerance.` |
| **MassQL Query Link**<br>https://proteomics2.ucsd.edu/ProteoSAFe/status.jsp?task=f71f79b124a04c37b4f5964fd8c2aad5 |
| **Additional Data Analysis**<br>n/a |
| **Data Availability**<br>MSV000087996 |

Myxobacteria are a potential keystone taxa within soil that contribute to nutrient cycling as generalist predators. Perception of acylhomoserine lactone (AHL) signaling molecules secreted by prey bacteria or quorum signal "eavesdropping" increases the predatory capacity of the myxobacterium *Myxococcus xanthus*. Our recent discovery of horizontally acquired AHL synthases from myxobacterial genome data suggests that myxobacteria may also secrete quorum signals to influence or disrupt cooperative behaviors of prey. While we have previously demonstrated the functionality of an AHL synthase from the myxobacterium *Archangium gephyra* via heterologous expression in *Escherichia coli*, we have yet to observe production of AHLs from *A. gephyra* or any other myxobacteria that possess an AHL synthase. We suspect that variability of the *N*-acyl side chain component of AHLs or absence of environmental stimuli in axenic cultivation conditions have hindered detection of AHL biosynthesis from myxobacteria using molecular networking and traditional biosensor-based approaches. Utilizing Mass Query Language (MassQL) we are able to search untargeted mass spectrometry data from myxobacterial extracts for features that include common fragmentation patterns associated with the core homoserine lactone moiety of AHLs. This search provides the opportunity to quickly determine if AHL-like metabolites are present in myxobacterial extracts despite *N*-acyl side chain variability. Such rapid analysis also enables us to explore a variety of environmental conditions such as predator-prey interactions that may induce myxobacterial AHL biosynthesis. Ultimately, MassQL will enable analysis of extracts from myxobacteria to detect production of AHLs and facilitate future efforts to determine the utility of AHL synthases acquired from prey.

#### 1.9 Looking for Potential New Antibiotics by Searching Aminoglycosidic Compounds in Natural Products using MassQL

**Author/s: Anelize Bauermeister**

**MassQL Query**

```
QUERY scaninfo(MS2DATA) WHERE
MS2PROD=204.1:TOLERANCEMZ=0.1:INTENSITYPERCENT=10 AND
MS2PROD=186.1:TOLERANCEMZ=0.1:INTENSITYPERCENT=5 AND
MS2PROD=168.1:TOLERANCEMZ=0.1:INTENSITYPERCENT=5
```

**MassQL Translation**

```
Returning the scan information on MS2.
The following conditions are applied to find scans in the mass spec data.
Finding MS2 peak at m/z 204.1 a 0.1 m/z tolerance and a minimum percent
intensity relative to base peak of 10.0%.
Finding MS2 peak at m/z 186.1 a 0.1 m/z tolerance and a minimum percent
intensity relative to base peak of 5.0%.
Finding MS2 peak at m/z 168.1 a 0.1 m/z tolerance and a minimum percent
intensity relative to base peak of 5.0%.
```

**MassQL Query Link**

https://proteomics2.ucsd.edu/ProteoSAFe/status.jsp?task=2407b315c9c54383b7887794a2a47d7b

**Additional Data Analysis**
**GNPS Molecular Networking**

https://gnps.ucsd.edu/ProteoSAFe/status.jsp?task=7fd7e6c34cd14ec28592fee0fe92cc0f

**Data Availability**

MSV000085018

There are several drugs, including antibiotics and anticancer agents, that present as an aminoglycoside attached to the aglycone. Doxorubicin and daunorubicin, for instance, are anticancer anthracyclines that contain an anthracyclinone and a daunosamine (sugar moiety), which play a very important role in the DNA-binding stabilization of the drug[10]. Such aminoglycosides present a characteristic fragmentation pattern[11,12] and the fragment ions can be easily identified in an MS/MS spectrum. Therefore, searching for new structures that present a glycoside with an amino group attached can be a great approach in Natural Products in the searching for new potential drugs.

This example shows fragmentation and formation of fragment ions *m/z* 204, *m/z* 186 and *m/z* 168 from the aminoglycoside moiety present in the brasilicardin structure (**SI Figure 1.9 - 1**). These fragment ions were used in MassQL to search for other compounds that present the same fragment ions. The molecular networking grouped 7 different metabolites, including the brasilicardin used for the search, and also groups in a separated family the ions referring to the same compounds with double charge. These results highlight the potential of MassQL to search for compounds with aminoglycosides. This tool can be used to search any kind of aminoglycosides, in the search for new potential metabolites with antibiotics or anticancer activities in LC-MS/MS untargeted data of crude extracts.

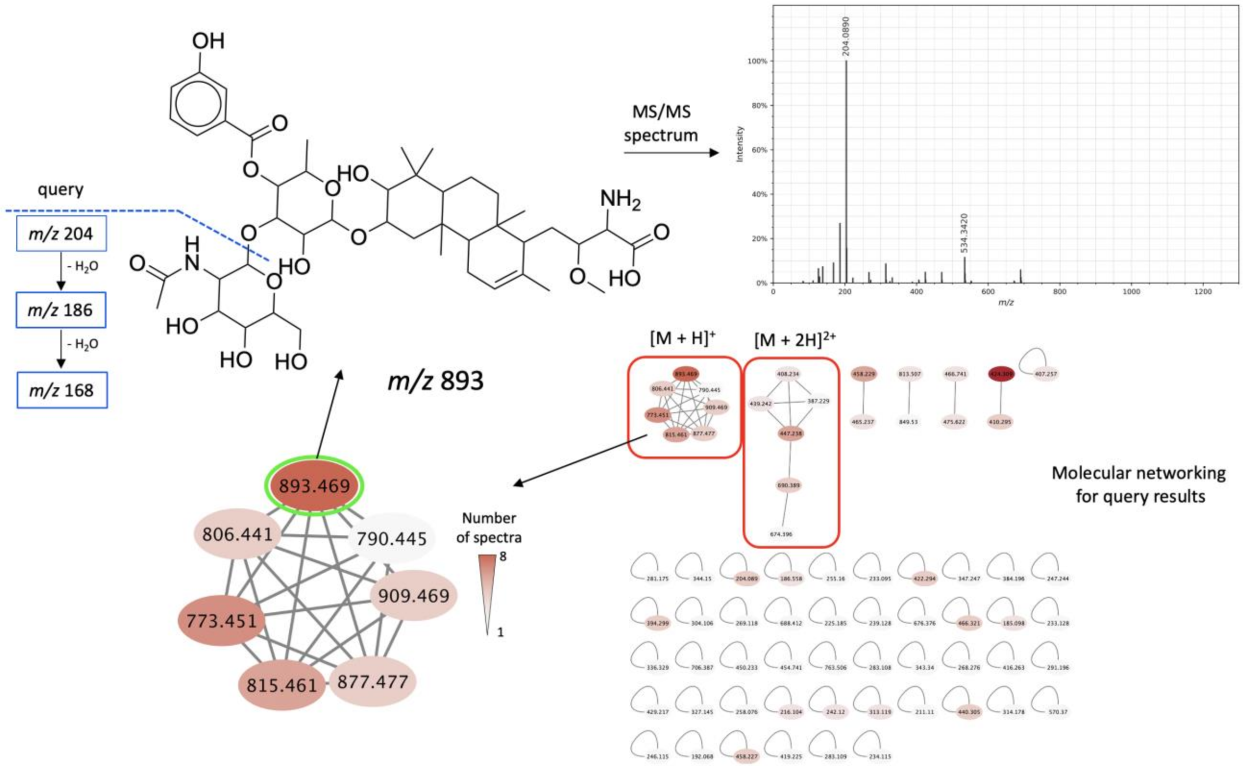


**SI Figure 1.9 - 1**. Query from brasilicardin used as an example for the search of metabolites with aminoglycosides. The molecular networking of the resulting spectra shows 7 analogs of brasilicardin that contain the same aminoglycoside.

##### 1.10 Gallic acid derivatives as potential anti-inflammatory agents in Plants

**Author/s: Wender Gomes**

| |
|---|
| **MassQL Query**<br>`QUERY scaninfo(MS2DATA) WHERE`<br>`MS2PROD=169.01:TOLERANCEMZ=0.1:INTENSITYPERCENT=30 AND`<br>`MS2PROD=125.02:TOLERANCEMZ=0.1:INTENSITYPERCENT=10` |
| **MassQL Translation**<br>`Returning the scan information on MS2.`<br>`The following conditions are applied to find scans in the mass spec data.`<br>`Finding MS2 peak at m/z 169.01 a 0.1 m/z tolerance and a minimum percent intensity relative to base peak of 30.0%.`<br>`Finding MS2 peak at m/z 125.02 a 0.1 m/z tolerance and a minimum percent intensity relative to base peak of 10.0%.` |
| **MassQL Query Link**<br>https://proteomics2.ucsd.edu/ProteoSAFe/status.jsp?task=797b825c3e7c48b182c8881b37eadd08 |
| **Additional Data Analysis**<br>n/a |
| **Data Availability**<br>MSV000088562 |

3,4,5-trihydroxybenzoic acid is generally known as gallic acid (GA), and it is a natural secondary metabolite which can be found in many natural sources, such as fruits, nuts, and plants. In recent years, that compound and derivatives has been receiving attention due to the powerful anti-inflammatory properties[13], and the galloyl group is strongly correlated to molecular action mechanisms. Herein, based on previous reports to the fragmentation pathway of gallic acid, we showed the search for derivatives of that compound using product ions reported to the galloyl group (MS2 peaks, at *m/z* 169.01 and *m/z* 125.02). For that, we used a dataset (MSV000088562) from a plant reported in the Brazilian Amazon as an anti-inflammatory natural source[14]. Figure 1.10A shows a compound annotated using GNPS library that was previously reported with inflammatory effects[15], and the examples in Figure 1.10B show compounds which have the product ions used as query. Results highlight the potential of MassQL to search for potential anti-inflammatory agents in natural products.

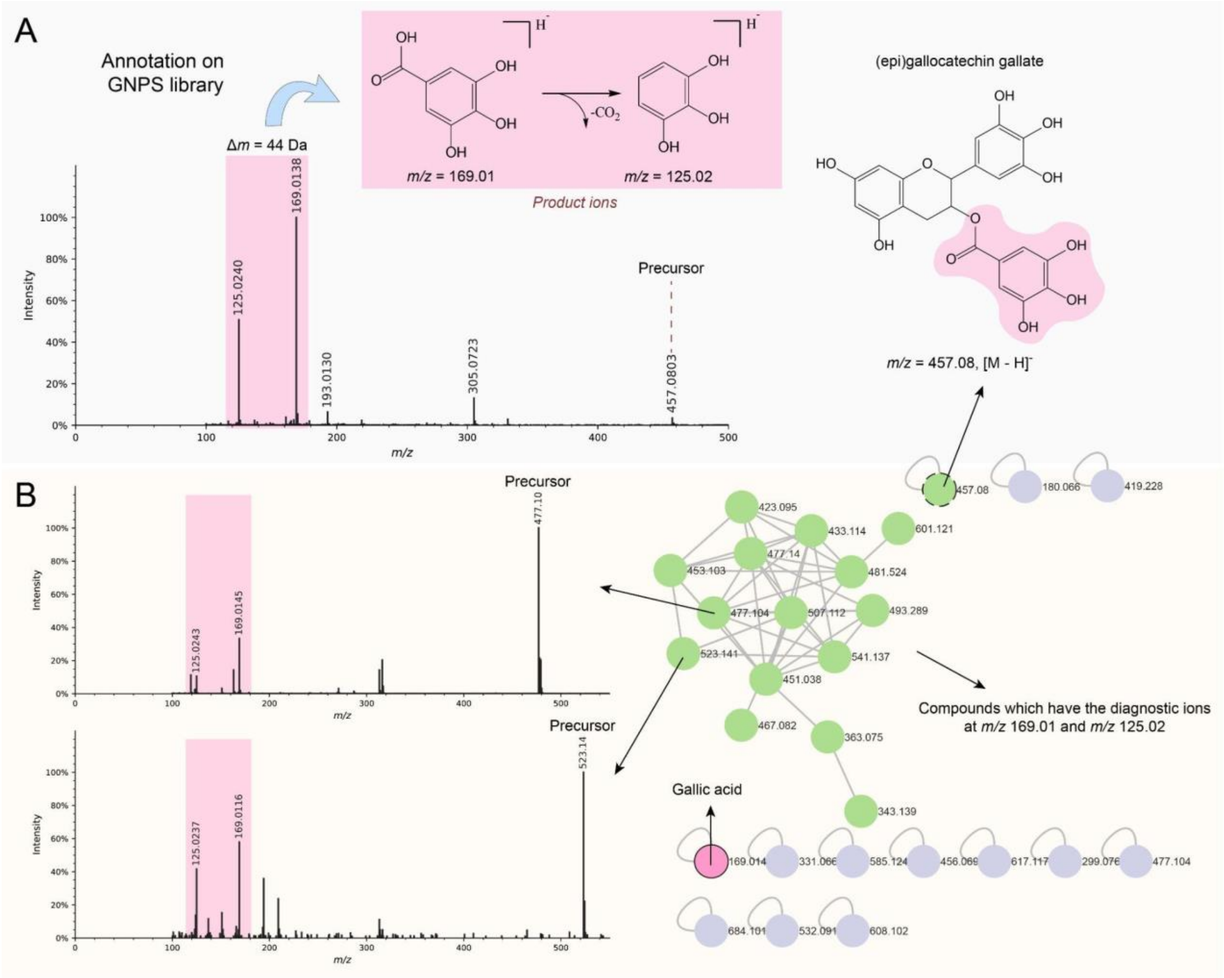


**SI Figure 1.10.1 - 1.** A) Fragmentation pathway and queries to diagnostic the Gallic acid derivatives (*m/z* 169.01 and *m/z* 125.02) detected to (epi)gallocatechin gallate, compound annotated on GNPS library; B) Network highlighted in green to compounds that have the diagnostic ions to gallic acid derivatives.

##### 1.11.1 Finding Lipid Tail Length Analogs with MassQL

**Author/s:** **Paul D. Boudreau and Mohammed M. A. Ahmed**

| |
|---|
| **MassQL Query**<br>`QUERY scaninfo(MS2DATA) WHERE`<br>`MS2PROD=523.2358:TOLERANCEPPM=10:INTENSITYPERCENT=5 AND`<br>`MS2PROD=212.1645:TOLERANCEPPM=10:INTENSITYPERCENT=5` |
| **MassQL Translation**<br>`Returning the scan information on MS2.`<br>`The following conditions are applied to find scans in the mass spec data.`<br>`Finding MS2 peak at m/z 523.2358 a 10.0 PPM tolerance and a minimum percent intensity relative to base peak of 5.0%.`<br>`Finding MS2 peak at m/z 212.1645 a 10.0 PPM tolerance and a minimum percent intensity relative to base peak of 5.0%.` |
| **MassQL Query Link**<br>https://proteomics2.ucsd.edu/ProteoSAFe/status.jsp?task=7489ac36a35b4dbc8e096a75b254e6bf |
| **Additional Data Analysis**<br>n/a |
| **Data Availability**<br>MSV000088046 |

In annotating an extract of *Cupriavidus necator* that produced cupriachelins, an interesting analog was discovered which replaced one of the 3-hydroxy-aspartic acids with glycine. Lacking the second 3-hydroxy-aspartic acid, these analogs are presumed to have a great difference in metal binding activity and were of interest to our group. Lipid tail length variation is common among lipopeptides, and has been reported before with the original isolation of the cupriachelins[16], so we set out to use MassQL to identify these tail length analogs of our glycine analog. We searched the MS/MS data for spectra bearing a canonical cupriachelin fragment (*m/z* 523) and lipid tail fragments containing the glycine (**Figure 1.11.1 - 1**). As expected, searching with the C10 lipid tail length fragment (*m/z* 212) found the glycine analog of interest.

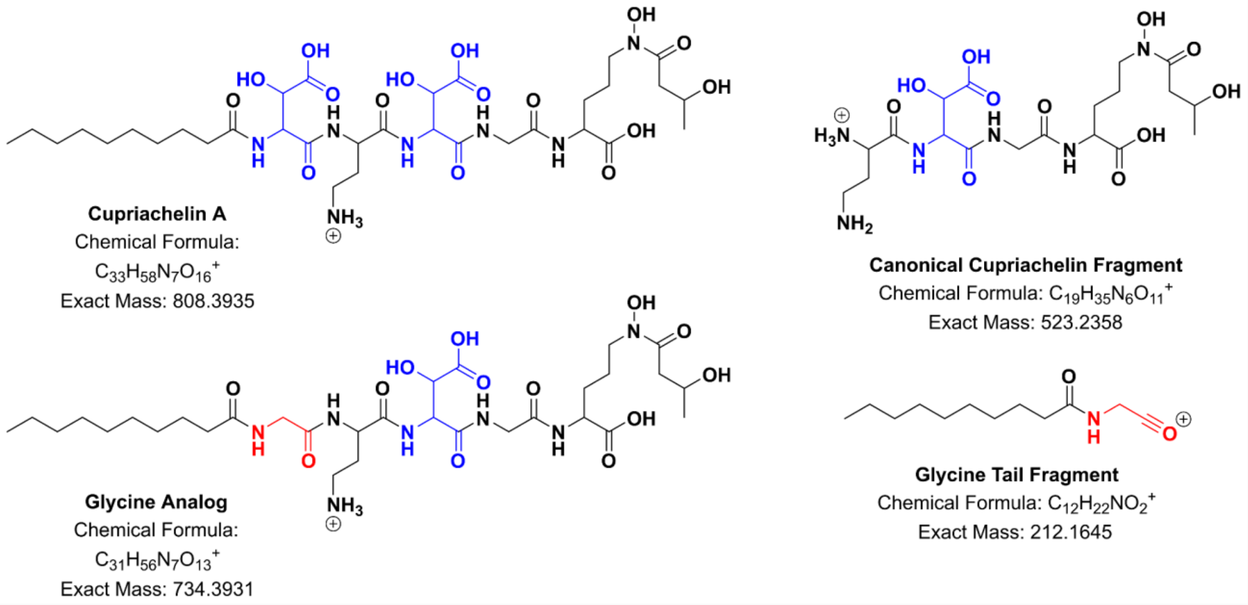


**Figure 1.11.1 - 1.** The *m/z* 523 ion is shared across all tail length variants of the cupriachelins, our "canonical" ion. A MassQL search with this fragment and the fragment ion bearing the lipid tail attached to our glycine residue of interest identified this analog parent mass at *m/z* 734.

##### 1.11.2 Finding Lipid Tail Length Analogs with MassQL: Short Tail Variant

**Author/s: Paul D. Boudreau and Mohammed M. A. Ahmed**

| |
|---|
| **MassQL Query**<br>`QUERY scaninfo(MS2DATA) WHERE`<br>`MS2PROD=523.2358:TOLERANCEPPM=10:INTENSITYPERCENT=5 AND`<br>`MS2PROD=184.1332:TOLERANCEPPM=10:INTENSITYPERCENT=5` |
| **MassQL Translation**<br>`Returning the scan information on MS2.`<br>`The following conditions are applied to find scans in the mass spec data.`<br>`Finding MS2 peak at m/z 523.2358 a 10.0 PPM tolerance and a minimum percent intensity relative to base peak of 5.0%.`<br>`Finding MS2 peak at m/z 184.1332 a 10.0 PPM tolerance and a minimum percent intensity relative to base peak of 5.0%.` |
| **MassQL Query Link**<br>https://proteomics2.ucsd.edu/ProteoSAFe/status.jsp?task=3848e7093eb542e88d60fa6960093380 |
| **Additional Data Analysis**<br>n/a |
| **Data Availability**<br>MSV000088046 |

This subsequent search, changing the tail fragment by -28.0313, a loss of two carbons to the tail, also identified a glycine analog with a different tail length (**SI Figure 1.11.2 - 1**).

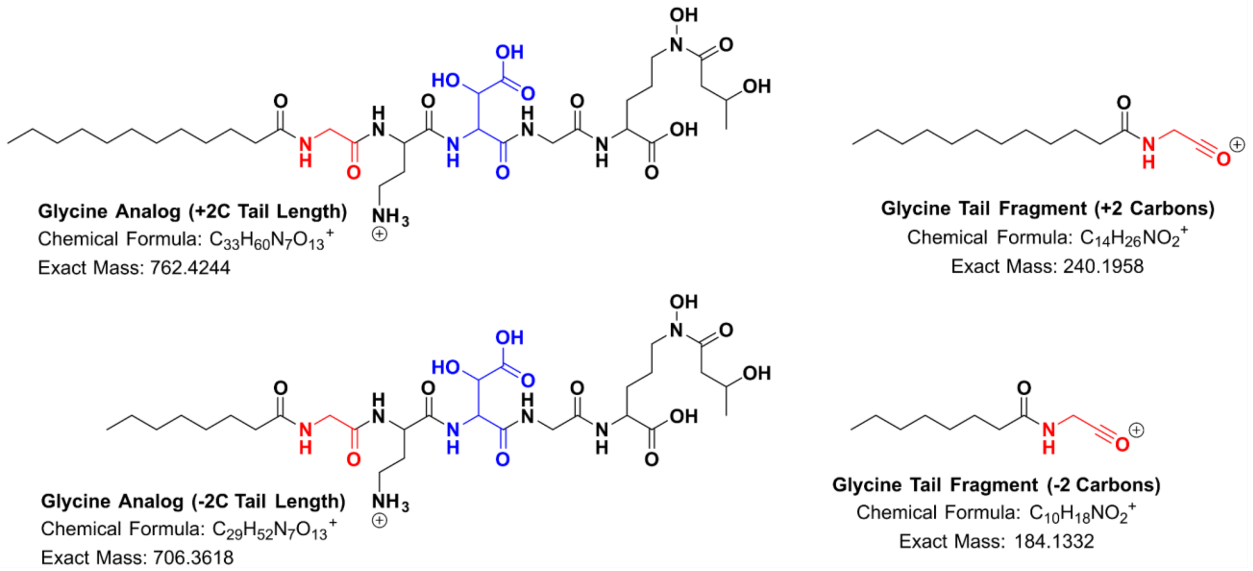


**SI Figure 1.11.2 - 1.** The same search for our core *m/z* 523 ion where we modified the mass of the tail fragment ion easily identified new analogs of varying tail lengths (*m/z* 762 and *m/z* 706).

##### 1.11.3 Finding Lipid Tail Length Analogs with MassQL: Longer Tail Variant

**Author/s: Paul D. Boudreau and Mohammed M. A. Ahmed**

| |
|---|
| **MassQL Query**<br>`QUERY scaninfo(MS2DATA) WHERE`<br>`MS2PROD=523.2358:TOLERANCEPPM=10:INTENSITYPERCENT=5 AND`<br>`MS2PROD=240.1958:TOLERANCEPPM=10:INTENSITYPERCENT=5` |
| **MassQL Translation**<br>`Returning the scan information on MS2.`<br>`The following conditions are applied to find scans in the mass spec data.`<br>`Finding MS2 peak at m/z 523.2358 a 10.0 PPM tolerance and a minimum percent intensity relative to base peak of 5.0%.`<br>`Finding MS2 peak at m/z 240.1958 a 10.0 PPM tolerance and a minimum percent intensity relative to base peak of 5.0%.` |
| **MassQL Query Link**<br>https://proteomics2.ucsd.edu/ProteoSAFe/status.jsp?task=e514b71ce1024d01842fe00d55380568 |
| **Additional Data Analysis**<br>n/a |
| **Data Availability**<br>MSV000088046 |

This subsequent search, changing the tail fragment by +28.0313, a gain of two carbons to the tail, also identified a glycine analog with a different tail length (**SI Figure 1.11.2 - 1**). Together, these three searches show how MassQL is a fast and simple tool for identifying lipid tail length analogs in lipopetides, which can sometimes be confounded by isobaric constitutional isomers modified elsewhere in the lipopeptide. Indeed, other groups could also use MassQL to search for the wide variety of known lipid tail modifications such as hydroxylation, unsaturation, or halogenation.

#### 1.12 Natural Product Peptidogenomics

**Author/s: Max Crüsemann**

##### Introduction

One of the most important goals in natural product discovery and the basis for any state-of-the-art biosynthetic study is the direct connection of a metabolite of interest to its biosynthetic gene cluster (BGC). The concept of natural product peptidogenomics is based on the linkage of a series of detected neutral amino acid mass losses (sequence tag) in a natural product MS/MS spectrum with parts of either the sequence of a ribosomally encoded and posttranslationally modified precursor peptide (RiPP) or with a series of predicted specificities of adenylation domains in a nonribosomal peptide synthetase (NRPS) megaenzyme, which both may be encoded in the producer's genome[17]. Although automated peptidogenomic workflows have subsequently been developed[18], these are rather difficult to handle and modify, may not include modified, nonproteinogenic amino acids and often do not yield valuable results. MassQL offers the flexible and fast detection of mass losses from MS/MS data already stored in GNPS and the design of individual queries, which can be easily modified and fine-tuned by the natural product genome miner. To validate this novel, powerful tool for natural product peptidogenomics, suitable queries were designed and validated with known examples.

##### Results: Validation with known Peptides

As a proof of concept, a specific query for tyrosine mass losses was designed to detect biarylitide A (*m/z* 522.20) (**SI Figure 1.12 - 1**), the first member of a novel class of ribosomally synthesized tripeptides (precursor sequence: MRYYH)[19] from parts of the recently published GNPS dataset of the actinobacterial genus *Planomonospora* (MSV000085376)[20].

| **MassQL Query** |
|---|
| `QUERY scaninfo(MS2DATA) WHERE MS2PROD=X:INTENSITYPERCENT=40:TOLERANCEMZ=0.002 AND MS2PROD=X-163.063:INTENSITYPERCENT=40:TOLERANCEMZ=0.002` |
| **MassQL Translation** |
| `Returning the scan information on MS2.`<br>`The following conditions are applied to find scans in the mass spec data.`<br>`Finding MS2 peak at m/z X a minimum percent intensity relative to base peak of 40.0% and a 0.002 m/z tolerance.`<br>`Finding MS2 peak at m/z X-163.063 a minimum percent intensity relative to base peak of 40.0% and a 0.002 m/z tolerance.` |
| **MassQL Query Link** |
| https://proteomics2.ucsd.edu/ProteoSAFe/status.jsp?task=66814f5f6dd245dd909b293044f09feb |
| **Additional Data Analysis** |

| |
|---|
| n/a |
| **Data Availability**<br>MSV000085376 |

Another query for two amino acid mass mass losses, one of them for a non-proteinogenic amino acid, was designed. Here, the nonribosomal cyclic depsipeptide FR900359 (FR) (Figure 1.12.2 - 1) was used as an example. FR contains the building blocks alanine and methylalanine, which are biosynthesized by an NRPS module containing an alanine-activating adenylation (A) domain and another module with an alanine-specific A domain and a methyltransferase encoded in the FR BGC (MiBIG BGC0001451). Querying a recently published dataset that compares FR producers and culture conditions (MSV000086958)[21] resulted in the detection of FR (*m/z* 1002.54) and all known FR analogs in the dataset.

**MassQL Query**

```
QUERY scaninfo(MS2DATA) WHERE
MS2PROD=X:INTENSITYPERCENT=40:TOLERANCEMZ=0.002 AND MS2PROD=X-
85.052:INTENSITYPERCENT=40:TOLERANCEMZ=0.002
|||
QUERY scaninfo(MS2DATA) WHERE
MS2PROD=X:INTENSITYPERCENT=40:TOLERANCEMZ=0.002 AND MS2PROD=X-
71.037:INTENSITYPERCENT=40:TOLERANCEMZ=0.002
```

**MassQL Translation**

N/A

**MassQL Query Link**

https://proteomics2.ucsd.edu/ProteoSAFe/status.jsp?task=5c7dc85ad5a14cb5827affd88903b913

**Additional Data Analysis**

n/a

**Data Availability**

MSV000086958

**Discussion**

MassQL enables individual, targeted searches for mass losses in MS/MS spectra, and is thus a promising tool for performing natural product peptidogenomic experiments, i.e. matching NRPS/RiPP BGCs with MS/MS data. Particularly advantageous in comparison to the automated pipelines is the possibility to search individually for BGC-specific mass losses, derived from detailed manual bioinformatic predictions, i.e. modified, nonproteinogenic amino acids, but also for other building blocks such as acyl groups or (modified) sugars. Furthermore, the possibility to fine-tune the queries depending on the quality, resolution and amount of the available MS/MS data and the number of hits makes this tool particularly powerful. Thus, MassQL is expected to accelerate and improve many genome-based natural product discovery projects.

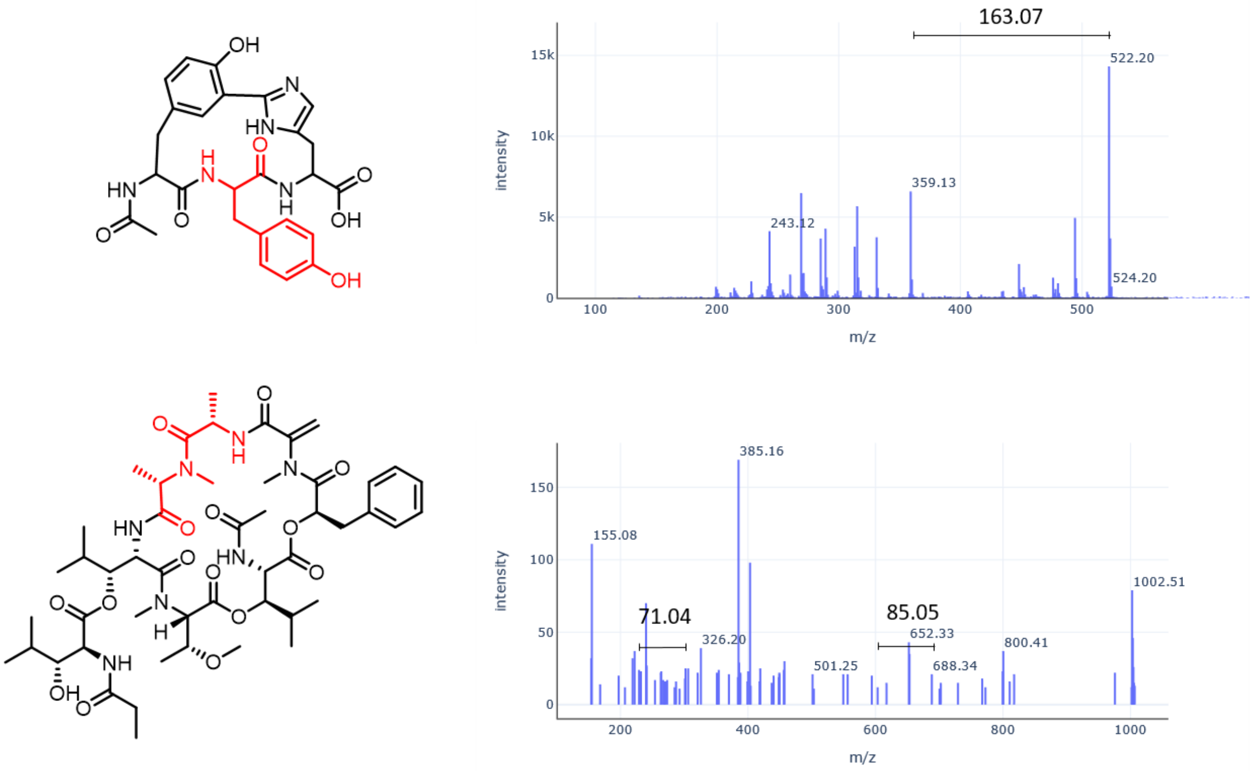


**SI Figure 1.12 - 1.** Detection of tyrosine mass loss from biarylitide A (upper panel), detection of mass losses corresponding to alanine and methylalanine from FR900359 (lower panel). MS2 spectra extracted from GNPS LCMS Dashboard[20].

#### 1.13 Exploring *in vitro* Liver Metabolism of Xenobiotics using MassQL

**Author/s: William J. Crandall and Ken H. Liu**

| **MassQL Query**<br>QUERY scaninfo(MS2DATA) WHERE MS2NL=225.1367:TOLERANCEPPM=10 |
|---|
| **MassQL Translation**<br>Returning the scan information on MS2.<br>The following conditions are applied to find scans in the mass spec data.<br>Finding MS2 neutral loss peak at m/z 225.1367 a 10.0 PPM tolerance. |
| **MassQL Query Link**<br>https://proteomics2.ucsd.edu/ProteoSAFe/status.jsp?task=b652d8cb059a40cba2fe9ea7feeb98ae |
| **Additional Data Analysis**<br>Network analysis:<br>https://gnps.ucsd.edu/ProteoSAFe/status.jsp?task=11da3ab0593748aeb3921147d659cb49 |
| **Data Availability**<br>N/A |

**Background**

Pooled human liver microsomal s9 fractions are used for the *in vitro* high throughput analysis of metabolic products of xenobiotics. A typical workflow includes the curation and analysis of data using feature extraction software and custom written scripts to select peaks corresponding to known metabolic mass shifts. Further characterization of these potential metabolites is done via the criteria of expected RT shift and increase in peak abundance over time of incubation with the enzymes. Potentially identified metabolites could greatly benefit from structural MS2 information and the assessment of structural similarity to the parent xenobiotic compound.

**Methods**

Pooled human liver S9 fractions (20 mg/ml protein) were aliquoted into 0.5 ml microcentrifuge tubes, stored at − 80 °C, and then thawed at room temperature prior to use. Standards were prepared in DMSO and further diluted 1:1000 in water for a final testing concentration of 50 µM. The NADPH regenerating system was reconstituted with addition of 3.5 ml of water to make a final volume of 5 ml. Cofactors were combined to form a 4X cofactor stock as follows, prior to addition into the reaction mixture: 10 mM UDPGA, 2 mM GSH, 2 mg/ml PAPS, 0.1 mM acetyl-CoA, and NADPH regenerating system (1 mM NADP, 5 mM glucose-6-phosphate, 1 unit glucose-6-phosphate dehydrogenase). Reactions were carried out at 30 °C on 96-well plates. S9 fraction was diluted 10-fold in water immediately before mixing with 0.2 M Tris-Cl, pH

7.5/2 mM MgCl2 and 0.3 mM of the xenobiotic solution in a 1:1:1 ratio (15 µL each). and incubated at 30 °C for 5 min. To start the reaction, 15 µL of 4X cofactor stock was added, and incubation was carried out at 30 °C for the indicated times. To terminate the reaction, we added a three-fold volume of acetonitrile, covered the plate with parafilm, vortexed, and froze at − 20 °C to precipitate insoluble materials such as proteins. After thawing and centrifugation of the incubation plate, the supernatants were transferred into polypropylene autosampler vials, which were stored at − 20 °C until instrumental analysis.

**Discussion**

We have found use in MassQL language for the semi-targeted networking of *in vitro* liver metabolic products in an efficient and high throughput manner. As a case example, we present the metabolic transformation of mitragynine using human liver microsomes. A neutral loss at *m/z* 225.1367 was looked for as this is typically the most abundant product ion peak. This neutral loss is conserved for multiple phase I metabolites. Key metabolites such as 9-O-demethyl mitragynine or 16-carboxy mitragynine (*m/z* 385.2120), and 7-hydroxymitragynine (*m/z* 415.2274) are clearly seen in the query results and in the network analysis (**Figure 1.13 - 1**). This MassQL workflow allows for the quick view of the data and relevant, high confidence metabolites. Addition of MS2 data curation into our current workflow increases the confidence of metabolite detection and association with the parent xenobiotic. In conclusion, this tool is complementary to our current workflow and will be useful for subsequent analysis due to its ease of use. The application of MassQL is especially useful for retrospective curation of MS2 data, as we have run over 600 unique xenobiotics through our *in vitro* system.

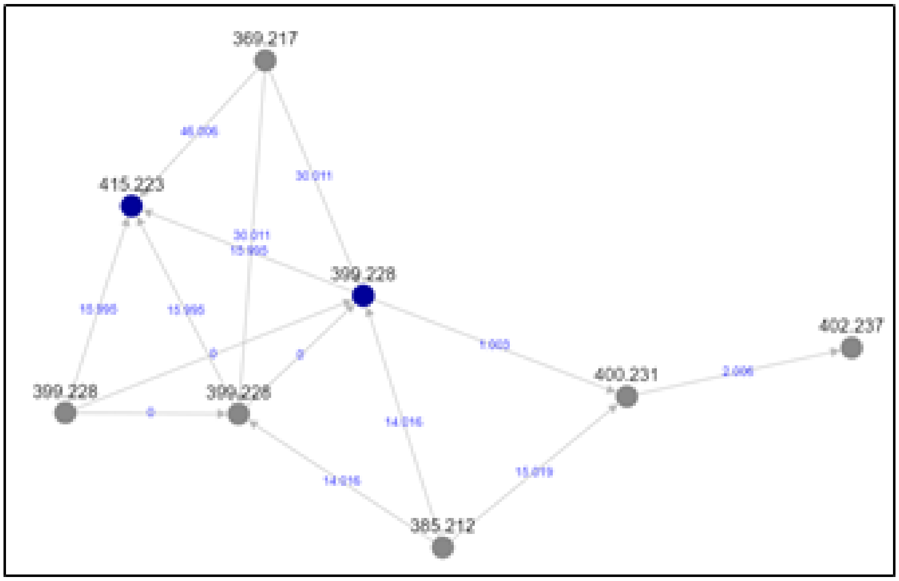


**SI Figure 1.13 - 1.** Molecular Network of neutral loss *m/z* 225.1367 from MassQL results. Mitragynine (*m/z* 399.228) and 7-hydroxymitragynine (*m/z* 415.223) are matching to GNPS databases. Additionally, (*m/z* 385.212) is found in the network, which is a major metabolite of mitragynine.

#### 1.14 MassQL Query for Xenobiotic Conjugation of Malonyl Glucose Conjugates

**Author/s: Chris Brown, Deepa D. Acharya, Tao Xu, Ken Clevenger, Quanbo Xiong, Jeffrey R. Gilbert**

| **MassQL Query** |
|---|
| `QUERY scaninfo(MS2DATA) WHERE MS2NL=248.0535:TOLERANCEMZ=0.1 AND MS2PROD=331.0056:TOLERANCEMZ=0.1` |
| **MassQL Translation** |
| Returning the scan information on MS2.<br>The following conditions are required to return MS2 scan.<br>Finding MS2 scans where neutral loss of 248.0535 Da with a tolerance of 0.1 Da is observed.<br>Finding MS2 peak at 331.0056 with a 0.1 Da tolerance. |

The identification of major metabolites related to an active ingredient is a requirement for registration of agrochemicals around the world. This requirement often necessitates trace level identification of metabolites from complex environmental matrices. Identification of MS/MS product ions that are characteristic of the active ingredient represents one way to begin to distinguish these metabolites. Using  MassQL as an initial filter might aid in organizing product ion spectra prior to structure elucidation.

Plant cell cultures (wheat, soybean, or blackgrass cell lines) were dosed with the active ingredient of the Arylex herbicide. Metabolites generated from that experiment were extracted, chromatographically separated and analyzed on a Thermo Fusion Lumos in positive mode using standard data dependent acquisition methods. Resulting data files were converted to mzML format with ProteoWizard msconvert tool and uploaded to Ometa Labs for analysis using the MassQL workflow.

The conjugation of an active ingredient or its metabolites with glucose or malonyl glucose is common in plant systems. These conjugates represent an analytical challenge because they are labile species when studied using gas phase fragmentation techniques, and therefore special attention is often needed to elucidate their structures. In this example, the neutral loss of 248 Da represents the loss of a malonyl glucose conjugate, while the fragment ion mass of *m/z* 331 is characteristic of the herbicide being used in this study. MassQL can quickly filter spectra that allow the researcher to study these complex species and focus their structure elucidation efforts.

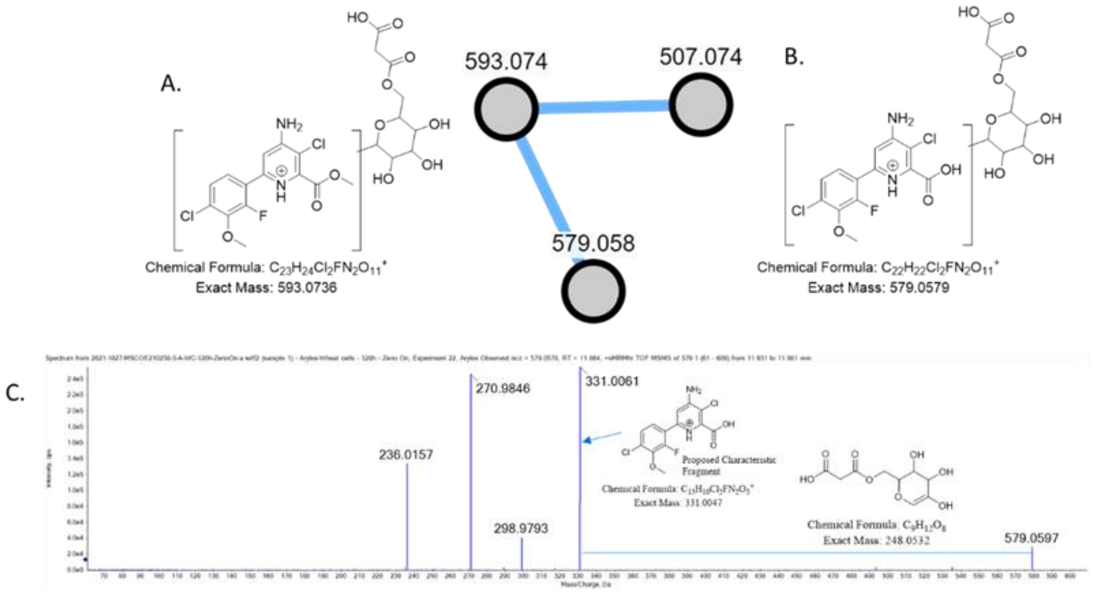


**SI Figure 1.13.1**- Markush structures for two proposed malonyl glucose species observed when running a MassQL query and then molecular networking the resulting spectra are shown in A. A MS2 spectrum showing a malonyl glucose neutral loss of 248 Da and a proposed structure for the characteristic fragment ion are shown in part C.

#### 1.15 Sulfatome analysis of urine samples

**Author/s: Correia, Mario S.P., Globisch, Daniel**

| |
|---|
| **MassQL Query**<br><br>`QUERY scaninfo(MS2DATA) WHERE MS2PROD=X:INTENSITYPERCENT=10 AND MS2PROD=X-79.9568:INTENSITYPERCENT=10` |
| **MassQL Translation**<br><br>Returning the scan information on MS2.<br>The following conditions are applied to find scans in the mass spec data.<br>Finding MS2 peak at m/z X a minimum percent intensity relative to base peak of 10.0%.<br>Finding MS2 peak at m/z X-79.9568 a minimum percent intensity relative to base peak of 10.0%. |
| **MassQL Query Link**<br><br>https://gnps.ucsd.edu/ProteoSAFe/status.jsp?task=dd4db3a3a43e4afa9e064faa7b802b71 |

**Background**

Identification of sulfated metabolites in human samples provides a useful readout of the co-metabolism between the microbiota and their human host. Many examples of this metabolic interaction have been reported including the conversion of xenobiotics from dietary metabolites. In general, macromolecules are initially converted by the gut bacteria and can then further be metabolized to facilitate excretion by humans through urine samples. The MassQL query provides the opportunity to selectively search for sulfated compounds with the characteristic loss of 79.9568 Da in public datasets. This MassQL query is a powerful tool to advance the recently developed sulfatome analysis to identify yet unknown correlations of this co-metabolism readout.

All sulfates were previously identified in three studies and the masses of the previously identified sulfated metabolites were compared with the output data from the MassQL query[22–24]. The data utilized for this query was selected from either Correia et al.[23] or Ballet et al.[24]. The general fragmentation for sulfates has the following three characteristic fragments (**SI Fig. 1.15 - 1**).

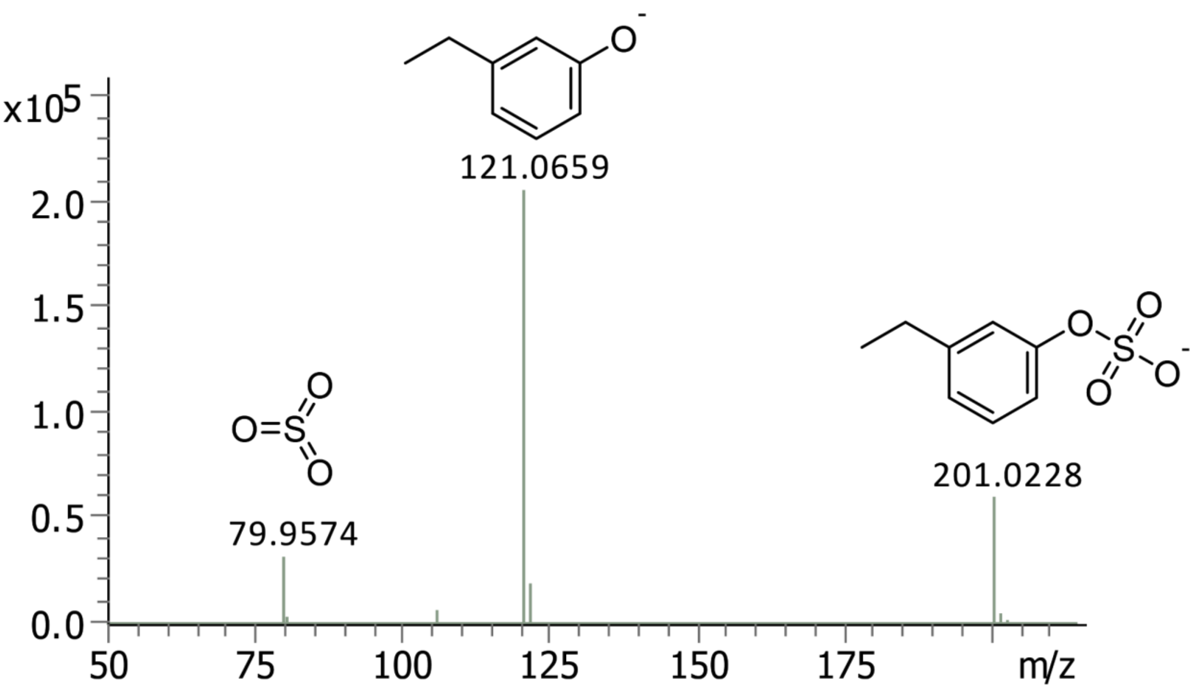


**SI Figure 1.15 - 1** - Fragmentation spectrum for 4-ethylphenylsulfate. Representative annotation of the three main fragmentation peaks that was generally used for the identification of sulfated metabolites ($[M\text{-}H]^-$, $[M\text{-}SO_3\text{-}H]^-$ and $[SO_3]$)

Several sulfated metabolites identified with the MassQL query were present in our previously published data demonstrating the potential of MassQL to identify this compound class. Among those, two example compounds derived from a polyphenolic rich diet are enterodiol glucuronide sulfate or enterolactone sulfate. Sulfate conjugates of *p*-cresol sulfate and dihydroxyindole were identified as well in this query. These compounds are known microbiota-derived metabolites and were recently described to be altered with age.

Most of our data used during identification in previous publications was acquired in negative mode mass spectrometric analysis. The majority of data reported in most databases is in positive mass spectrometric mode analysis. The insights gained from this query based on acquisition in negative mode analysis is of additional value to the scientific community.

#### 1.16 Data exploration for glycosylated macrolides produced in Actinomycetes

**Author/s: Eftychia E. Kontou and Tilmann Weber**

**MassQL Query**

```
QUERY scaninfo(MS2DATA) WHERE
MS2PROD=440.214:TOLERANCEMZ=0.1 AND
MS2PROD=131.089:TOLERANCEMZ=0.1 AND
MS2NL=439.200:TOLERANCEMZ=0.1
```

**MassQL Translation**

```
Returning the summed scan information on MS2.
The following conditions are applied to find scans in the mass spec data.
Finding MS2 peak at m/z 440.214 and a 0.1 PPM tolerance.
Finding MS2 peak at m/z 131.089 and a 0.1 PPM tolerance.
Finding a neutral loss from precursor in the MS2 spectrum at mass 439.200
and a 0.1 PPM tolerance.
```

**MassQL Query Link**

https://gnps.ucsd.edu/ProteoSAFe/status.jsp?task=a22beb16ec4c4bb7a33f83cdaec5c851

**Additional Data Analysis**

**GNPS Feature-Based Molecular Networking**

https://gnps.ucsd.edu/ProteoSAFe/status.jsp?task=cd13be916f0d4c87b664c5aa6cd45a37

**Data Availability**

MSV000087858

Actinobacteria are prolific natural product / specialized metabolite producers. Being responsible for approximately 45% of all known bioactive natural products[25], they still play a key role in antibiotic discovery. There is a desperate need for new classes of antibiotics to tackle the continuous rise of antimicrobial resistant infections[26], and this urgency yielded new approaches and technologies in the field, such as the combination of different omics technologies.

One example is the integration of genomics and metabolomics[27]. First, genetic investigation of an organism takes place through high-quality whole genome sequencing, paired with a genome mining tool, such as antiSMASH[28], for the prediction of gene clusters responsible for the biosynthesis of different classes of compounds. Those predictions are analyzed either in an automated or manual way, and a candidate-strain containing an interesting biosynthetic gene cluster (BGC) in its genome is picked for further chemical investigation. Complementary to these genome mining approaches, metabolomics analysis takes place, which aims to detect possible compounds derived from the BGC of interest. One recent example of such strategy is the discovery of two novel macrolides, epemicins A and B, from *Kutzneria* sp. CA-103260[29,30].

Macrolides are an important class of antibiotics derived from type I polyketide biosynthesis. They consist of a large lactone ring and often of one or more sugar moieties that often are responsible for their bioactive properties. antiSMASH predictions indicated that the genome of strain *Kutzneria* sp. CA-103260 contains a BGC that is related to the aculeximycin BGC[31]. The name of this compound is derived from one of its sugar moieties, the trisaccharide aculexitriose (exact mass, 439.2053), which could possibly be a diagnostic fragment for the discovery of related compounds. Here, instead of an untargeted metabolomics approach, we have tested MassQL for the investigation of aculeximycin-related compounds from crude extracts of strain *Kutzneria* sp. CA-103260.

MassQL returned 44 spectra hits from 6 different mzml files. 11 of those spectra (25%) belonged to three different adduct ions of epemicin B ($[M+2H]^{2+}$, $[M+H]^{+}$, $[M+H+H_2O]^{+}$) detected in many different samples. The other 33 spectra represent approximately 10 unique features that generated no hits from the GNPS MS/MS library search. Molecular networking did not indicate any relevance between epemicin B and the other precursors. Epimicin A was not detected in those six samples or was only detected in very low intensities. These data prove that MassQL can significantly increase the throughput of a combined genomics and metabolomics approach in the part of the chemical investigation of predicted structures.

#### 1.17 Identification of cyclic peptide analytes in plant metabolomes

**Author/s: Roland D. Kersten**

Plants produce a large diversity of cyclic peptides via the ribosomal pathway with applications as sustainable pesticides in agriculture and with therapeutic potential for the treatment of cancer, viral infections and hypertension[32–35]. These ribosomally synthesized and posttranslationally modified peptides (RiPPs) from plants are generally macrocyclized via head-to-tail-linkage or side-chain-crosslinking[36,37]. While head-to-tail-cyclic peptides can be partially or fully sequenced by MS/MS analysis[38], known side-chain-macrocyclic RiPPs with 1-2 macrocyclic bonds and 4-8 amino acids yield few b-y-ion fragments for peptide sequencing but often at least two amino acid iminium ions in the low-m/z region of MS/MS spectra (**Figure 1.17 - 1**). As known plant RiPPs have less posttranslational modifications of amino acids compared to bacterial and fungal RiPPs[39], most detected iminium ions in plant RiPP MS/MS spectra correspond to proteinogenic amino acid masses (**Figure 1.17 - 2**, **Table 1.17 - 1**)[37]. In addition, several plant peptides such as americine or selanine A (**Figure 1.17 - 2**) have mono- or dimethylated N-termini, which can result in amino acid iminium ions with one or two methyl mass shifts, respectively (**Table 1.17 - 1**). Given the presence of these fragments in the MS/MS spectra of plant RiPPs, these natural products could therefore be concentrated in plant MS metabolomic data by searching for MS/MS spectra, which have iminium ion fragments masses of proteinogenic (i.e. unmodified) amino acid iminium ions. In a recent study, we searched 300 plant metabolomes for MS/MS spectra, which have *one* iminium ion of a proteinogenic amino acid, in order to characterize candidate side-chain-macrocyclic plant RiPPs. While multiple RiPP classes could be discovered through this search, the pool of candidate peptide molecular clusters was very large through the inclusion of non-peptidic analytes resulting in a slow identification process[37].

Herein, we applied MassQL with the cardinality command to search the same plant metabolomes for MS/MS spectra with *multiple* amino acid fragments and subsequent molecular networking for improved targeted identification of peptide-specific molecular clusters.

**MassQL Query (for analytes with *multiple* amino acid iminium ion)**

```
QUERY scaninfo(MS2DATA) WHERE MS2PROD=(58.06513 OR 60.04439 OR 70.06513 OR
72.08078 OR 74.06004 OR 84.04439 OR 84.08078 OR 86.09643 OR 87.05529 OR
88.0393 OR 88.07569 OR 100.11208 OR 101.07094 OR 101.10732 OR 102.05495 OR
102.09134 OR 104.05285 OR 110.07127 OR 114.12773 OR 115.08659 OR 115.12297
OR 116.0706 OR 118.0685 OR 120.08078 OR 124.08692 OR 129.10224 OR 129.11347
OR 129.13862 OR 130.08625 OR 132.08415 OR 134.09643 OR 136.07569 OR
138.10257 OR 143.12912 OR 148.11208 OR 150.09134 OR 157.14477 OR 159.09167
OR 164.10699 OR 173.10732 OR
187.12297):CARDINALITY=range(min=2,max=5):TOLERANCEPPM=10:INTENSITYPERCENT=
5
```

**MassQL Translation**

| |
|---|
| `Returning the scan information on MS2.`<br>`The following conditions are applied to find scans in the mass spec data.`<br>`Finding MS2 peak at m/z 58.06513 or 60.04439 or 70.06513 or 72.08078 or 74.06004 or 84.04439 or 84.08078 or 86.09643 or 87.05529 or 88.0393 or 88.07569 or 100.11208 or 101.07094 or 101.10732 or 102.05495 or 102.09134 or 104.05285 or 110.07127 or 114.12773 or 115.08659 or 115.12297 or 116.0706 or 118.0685 or 120.08078 or 124.08692 or 129.10224 or 129.11347 or 129.13862 or 130.08625 or 132.08415 or 134.09643 or 136.07569 or 138.10257 or 143.12912 or 148.11208 or 150.09134 or 157.14477 or 159.09167 or 164.10699 or 173.10732 or 187.12297 with a cardinality minimum of 2.0 and maximum of 5.0 and a 10.0 PPM tolerance and a minimum percent intensity relative to base peak of 5.0%.` |
| **MassQL Query Link**<br>https://gnps.ucsd.edu/ProteoSAFe/status.jsp?task=7f4e581da6f144239e48d7216dc4a994 |
| **Additional Data Analysis**<br>**GNPS Molecular Networking**<br>https://gnps.ucsd.edu/ProteoSAFe/status.jsp?task=5044e88d892c46ca97245524ff54f25a |
| **Data Availability**<br>MSV000087872, MSV000088114 |

LCMS datasets of the Matthaei botanical garden collected on a QExactive Orbitrap mass spectrometer were searched for MS/MS spectra with 2-5 amino acid iminium ions from **Table 1.17 - 1** (excluding masses below 58 Da) with the MassQL-cardinality command specified above. The resulting MS/MS spectra were then analyzed by molecular networking as follows:

A molecular network was created (**Figure 1.17 - 3**) using the online workflow (https://ccms-ucsd.github.io/GNPSDocumentation/) on the GNPS website (http://gnps.ucsd.edu). The data was filtered by removing all MS/MS fragment ions within +/- 17 Da of the precursor m/z. MS/MS spectra were window filtered by choosing only the top 6 fragment ions in the +/- 50Da window throughout the spectrum. The precursor ion mass tolerance was set to 0.05 Da and a MS/MS fragment ion tolerance of 0.25 Da. A network was then created where edges were filtered to have a cosine score above 0.7 and more than 5 matched peaks. Further, edges between two nodes were kept in the network if and only if each of the nodes appeared in each other's respective top 10 most similar nodes. Finally, the maximum size of a molecular family was set to 500, and the lowest scoring edges were removed from molecular families until the molecular family size was below this threshold. The spectra in the network were then searched against GNPS' spectral libraries. The library spectra were filtered in the same manner as the input data. All matches kept between network spectra and library spectra were required to have a score above 0.7 and at least 6 matched peaks..

The final molecular network of the MassQL-cardinality search for analytes with 2-5 amino acid iminium ions (**Figure 1.17 - 3**) was queried for (a) the presence of known plant RiPPs from **Figure 1.17 - 2** and (b) the relative occurrence of cyclic plant peptides predicted based on clustering with reference peptides in the network compared to the total node number. The relative occurrence of known plant RiPPs in the MassQL-cardinality network was 18% (i.e. 323 nodes in molecular clusters with annotated plant peptides among 1798 total nodes). In order to highlight the inclusion of target cyclic plant peptides with the MassQL-cardinality search, the Matthaei botanical datasets were analyzed with the following MassQL-MS2Prod command to search for MS/MS spectra which only included one amino acid iminium ion mass from **Table 1.17 - 1**. The resulting MassQL-searched spectra were also analyzed via molecular networking as specified above for the MassQL-cardinality-searched spectra. The resulting molecular network comprised 10029 nodes, which included 400 nodes (4%) associated with cyclic peptide references. The MassQL-cardinality-search network included almost all reference cyclic peptides except Cyclo-[VPPIFY] (**Figure 1.17 - 2**) which were present in the MassQL-MS2Prod network, which shows that a search criterium for spectra with 2-5 amino acids results in almost the same number of target analytes but alongside less undefined hit spectra. Given the high concentration of cyclic peptide nodes in the MassQL-cardinality-network, its undefined clusters represent good starting points for discovery of plant peptides with new macrocyclization chemistry, which highlights the utility of MassQL coupled to molecular networking in plant RiPP discovery.

| |
|---|
| **MassQL Query (for analytes with *one* amino acid iminium ion)**<br>`QUERY scaninfo(MS2DATA) WHERE MS2PROD=(58.06513 OR 60.04439 OR 70.06513 OR 72.08078 OR 74.06004 OR 84.04439 OR 84.08078 OR 86.09643 OR 87.05529 OR 88.0393 OR 88.07569 OR 100.11208 OR 101.07094 OR 101.10732 OR 102.05495 OR 102.09134 OR 104.05285 OR 110.07127 OR 114.12773 OR 115.08659 OR 115.12297 OR 116.0706 OR 118.0685 OR 120.08078 OR 124.08692 OR 129.10224 OR 129.11347 OR 129.13862 OR 130.08625 OR 132.08415 OR 134.09643 OR 136.07569 OR 138.10257 OR 143.12912 OR 148.11208 OR 150.09134 OR 157.14477 OR 159.09167 OR 164.10699 OR 173.10732 OR 187.12297):TOLERANCEPPM=10:INTENSITYPERCENT=5` |
| **MassQL Translation**<br>Returning the scan information on MS2.<br>The following conditions are applied to find scans in the mass spec data.<br>Finding MS2 peak at m/z 58.06513 or 60.04439 or 70.06513 or 72.08078 or 74.06004 or 84.04439 or 84.08078 or 86.09643 or 87.05529 or 88.0393 or 88.07569 or 100.11208 or 101.07094 or 101.10732 or 102.05495 or 102.09134 or 104.05285 or 110.07127 or 114.12773 or 115.08659 or 115.12297 or 116.0706 or 118.0685 or 120.08078 or 124.08692 or 129.10224 or 129.11347 or 129.13862 or 130.08625 or 132.08415 or 134.09643 or 136.07569 or 138.10257 or 143.12912 or 148.11208 or 150.09134 or 157.14477 or 159.09167 or 164.10699 or 173.10732 or 187.12297 a 10.0 PPM tolerance and a minimum percent intensity relative to base peak of 5.0%. |
| **MassQL Query Link**<br>https://gnps.ucsd.edu/ProteoSAFe/status.jsp?task=dcf546f952af411ca985ccf2b81db2f3 |
| **Additional Data Analysis**<br>**GNPS Molecular Networking**<br>https://gnps.ucsd.edu/ProteoSAFe/status.jsp?task=54a420ebd1ab463b96ad0d2d8bb4487e |

**Data Availability**
MSV000087872, MSV000088114

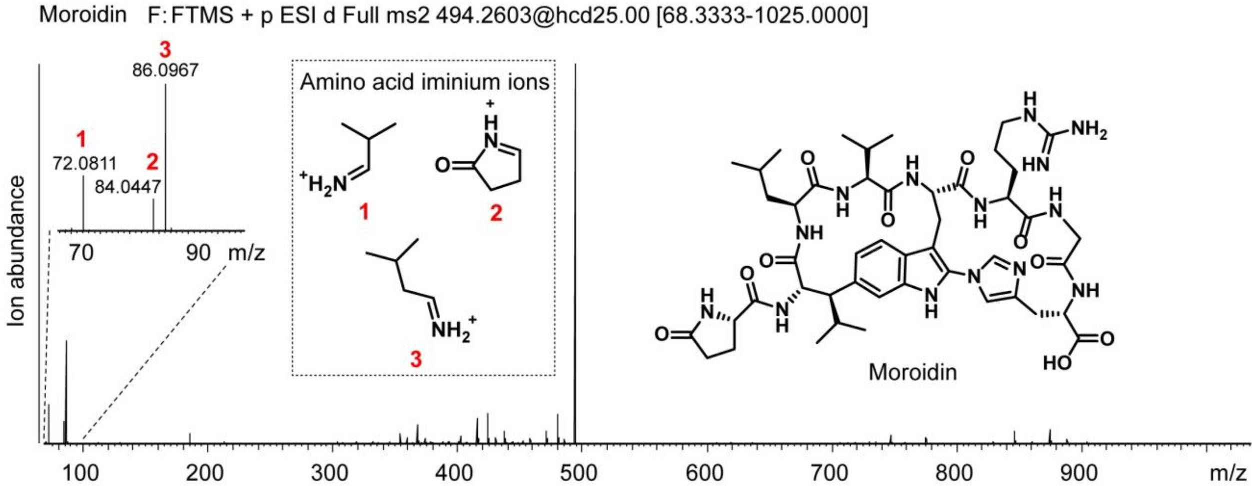


**Figure 1.17 - 1 | Representative MS/MS spectrum of side-chain-macrocyclic RiPP moroidin and detection of three amino acid iminium ions.**

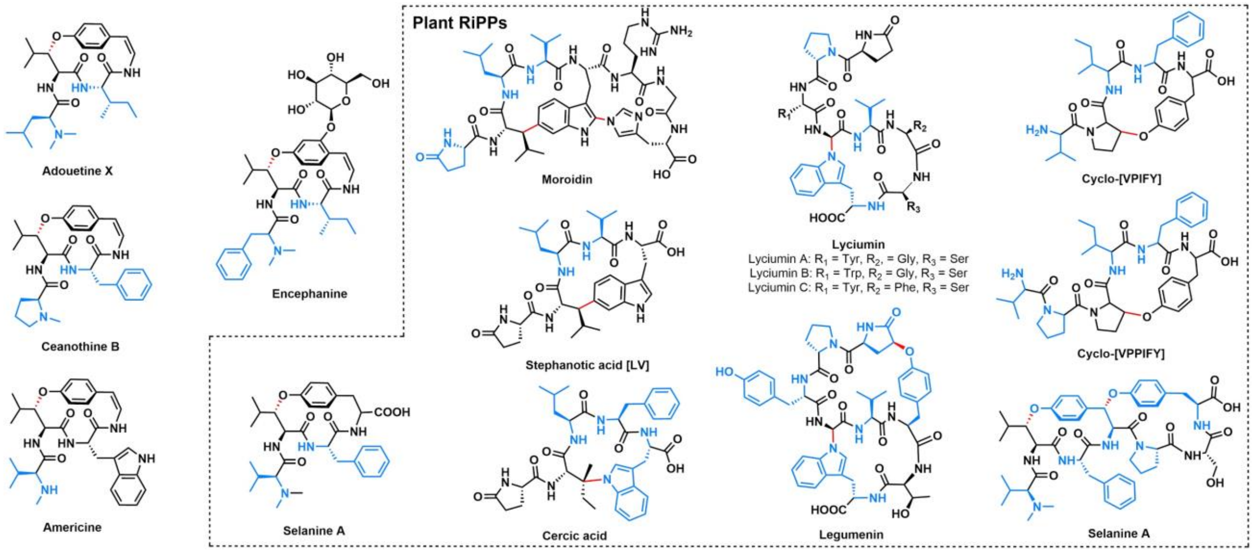


**Figure 1.17 - 2 | Representative structures of side-chain-macrocyclic plant peptides with highlighted residues detectable as iminium ions by tandem mass spectrometry.** (**A**) Peptide structures within the dashed lined box have been defined as plant RiPPs. Side-chain-macrocyclic bonds in all peptides are highlighted in red color. Amino acids which have been experimentally detected as corresponding iminium ion masses after high-energy collision induced dissociation in MS/MS spectra (fragmentation energy 25 eV) on a QExactive Orbitrap are colored in blue. Therefore, all shown peptides have 2-5 different amino acids which have been detected as iminium ions in specified MS/MS spectra.

**Table 1.17 - 1 | Iminium ion masses of proteinogenic amino acids, which are common building blocks of plant cyclic RiPPs.**

| Amino acid | m(calc, iminium ion) | m(calc, monomethylated iminium ion) | m(calc, dimethylated iminium ion) |
|---|---|---|---|
| Alanine | 44.04948 | 58.06513 | 72.08078 |
| Aspartate | 88.0393 | 102.05495 | 116.0706 |
| Glutamate | 102.05495 | 116.0706 | 130.08625 |
| Phenylalanine | 120.08078 | 134.09643 | 148.11208 |
| Glycine | 30.03383 | 44.04948 | 58.06513 |
| Histidine | 110.07127 | 124.08692 | 138.10257 |
| Leucine | 86.09643 | 100.11208 | 114.12773 |
| Isoleucine | 86.09643 | 100.11208 | 114.12773 |
| Lysine | 101.10732 | 115.12297 | 129.13862 |
| Methionine | 104.05285 | 118.0685 | 132.08415 |
| Asparagine | 87.05529 | 101.07094 | 115.08659 |
| Proline | 70.06513 | 84.08078 | n/a |
| Glutamine | 101.07094 | 115.08659 | 129.10224 |
| Arginine | 129.11347 | 143.12912 | 157.14477 |
| Serine | 60.04439 | 74.06004 | 88.07569 |
| Threonine | 74.06004 | 88.07569 | 102.09134 |
| Valine | 72.08078 | 86.09643 | 100.11208 |
| Tryptophan | 159.09167 | 173.10732 | 187.12297 |
| Tyrosine | 136.07569 | 150.09134 | 164.10699 |
| Pyroglutamate | 84.04439 | n/a | n/a |

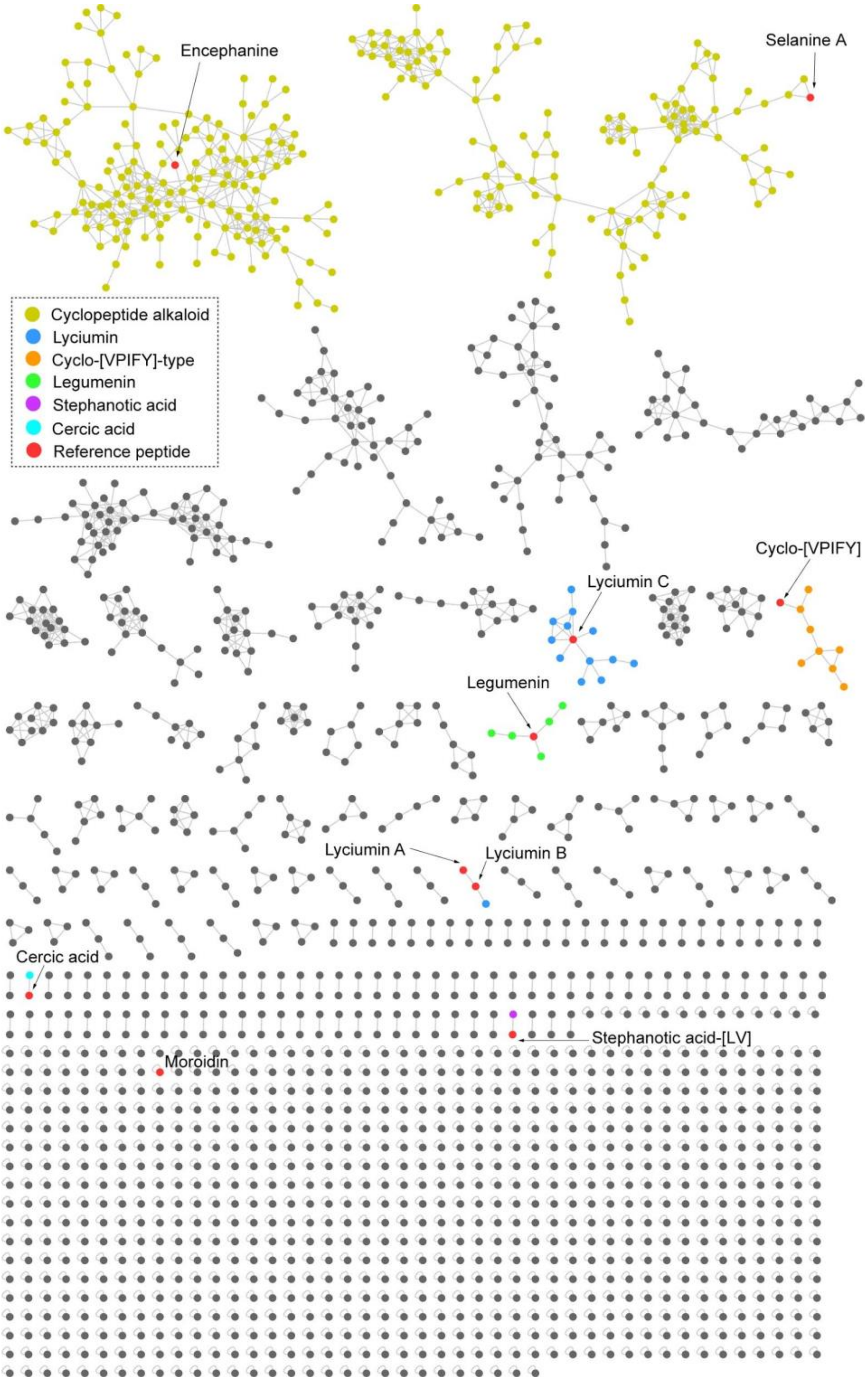


**Figure 1.17 - 3 | Molecular network of MassQL-processed LCMS-datasets with cardinality command of 2-5 different amino acid iminium ions in a given tandem MS spectrum.** Reference peptides from **Figure 1.17 - 1** are highlighted in red, predicted plant RiPPs based on reference peptide clustering are highlighted in color as described in node color legend and candidate peptides in undefined clusters are highlighted in grey.

### 2 MassQL Query using MS/MS Spectral Information and Multiple Queries

#### 2.1 Tracking The Biosynthesis of Isotopically Labeled Phenylpropanoids

**Author/s: Tomáš Pluskal**

| |
|---|
| **MassQL Query**<br>`QUERY scaninfo(MS1DATA) WHERE`<br>`MS1MZ=X:INTENSITYPERCENT=5 AND`<br>`MS1MZ=X+4.025:TOLERANCEMZ=0.001:INTENSITYPERCENT=5`<br>`|||`<br>`QUERY scaninfo(MS1DATA) WHERE`<br>`MS1MZ=X:INTENSITYPERCENT=5 AND`<br>`MS1MZ=X+5.0314:TOLERANCEMZ=0.001:INTENSITYPERCENT=5`<br>`|||`<br>`QUERY scaninfo(MS1DATA) WHERE`<br>`MS1MZ=X:INTENSITYPERCENT=5 AND`<br>`MS1MZ=X+6.0377:TOLERANCEMZ=0.001:INTENSITYPERCENT=5` |
| **MassQL Query Link**<br>https://proteomics2.ucsd.edu/ProteoSAFe/status.jsp?task=ec43a38302434bb7ab238b3273588a1f |
| **Additional Data Analysis**<br>n/a |
| **Data Availability**<br>MSV000087906 |

The phenylpropanoid pathway is a highly conserved biosynthetic pathway for polyketide biosynthesis in plants from cinnamic acid, which is produced by deamination of phenylalanine. We studied the biosynthesis of psychoactive polyketides called kavalactones in kava (*Piper methysticum*)[21]. In one experiment, we performed transient heterologous expression of the kavalactone biosynthetic enzymes SPS1, SPS2, and KOMT1 in tobacco (*Nicotiana benthamiana*) and simultaneously injected an isotopically labeled cinnamic acid-$d_6$ (Sigma cat. # 513962) into the plant. The goal of this experiment was to observe what downstream products are produced from the labeled cinnamic acid by the kavalactone biosynthetic enzymes as well as by native *N. benthamiana* enzymes.

In plants, cinnamic acid-$d_6$ is natively converted by hydroxylations and methylations to a variety of polyketide precursors, as shown here:

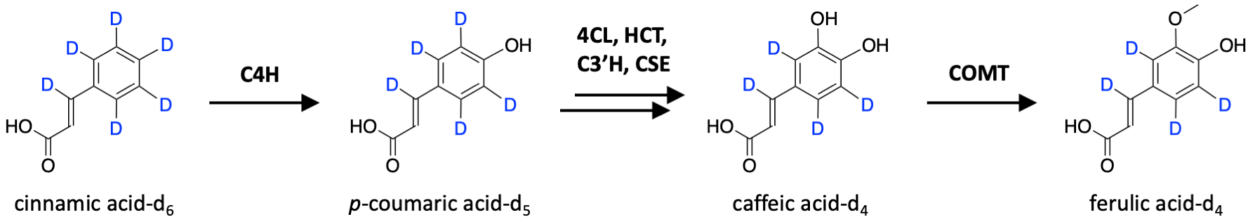


These compounds contain a different number of deuterium atoms, therefore their mass increase over the original (unlabeled) compound is also different:

| **Labeled compound** | **Mass increase (Da)** |
|---|---|
| cinnamic acid-$d_6$ | 6.0376 |
| *p*-coumaric acid-$d_5$ | 5.0313 |
| caffeic acid-$d_4$ | 4.0250 |
| ferulic acid-$d_4$ | 4.0250 |

By designing a MassQL query to find these mass differences in MS1 spectra, we were able to spot a number of downstream labeled products derived from the single labeled precursor in an untargeted manner.

One such product was chlorogenic acid-$d_4$, an ester of caffeic acid-$d_4$ and (−)-quinic acid. Chlorogenic acid is a common intermediate in lignin biosynthesis in plants and its biosynthesis does not require the kavalactone biosynthetic enzymes. However, this proved that the incorporation of the isotopically labeled precursor was successful.

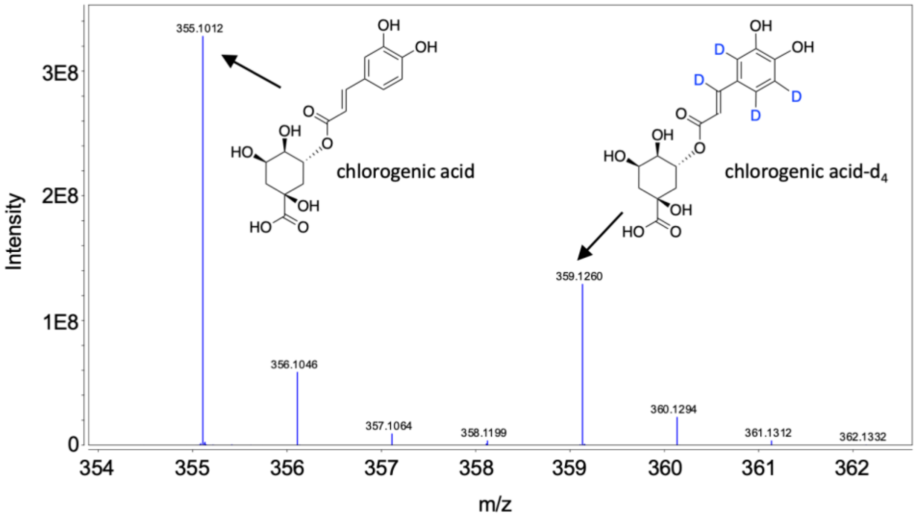


Link to the corresponding MS1 scan.

One of the main labeled products of the kavalactone biosynthetic enzymes that we observed was the labeled kavalactone yangonin-$d_5$, derived from *p*-coumaric acid-$d_5$.

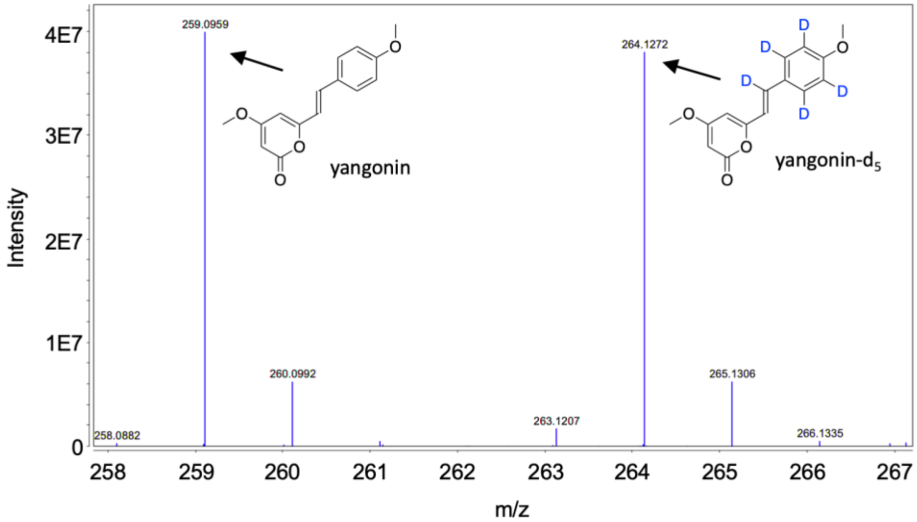


Link to the corresponding MS1 scan.

##### 2.2.1 Discovery of Novel Piperamides in *Piperaceae* plants using MassQL

**Author/s: Tito Damiani**

**MassQL Query**

```
QUERY scaninfo(MS2DATA) WHERE
MS2PROD=201.054:TOLERANCEPPM=5:INTENSITYPERCENT=1 AND
MS2PROD=135.044:TOLERANCEPPM=5:INTENSITYPERCENT=1
```

**MassQL Translation**

```
Returning the scan information on MS2.
The following conditions are applied to find scans in the mass spec data.
Finding MS2 peak at m/z 201.054 a 5.0 PPM tolerance and a minimum percent
intensity relative to base peak of 1.0%.
Finding MS2 peak at m/z 135.044 a 5.0 PPM tolerance and a minimum percent
intensity relative to base peak of 1.0%.
```

**MassQL Query Link**

https://proteomics2.ucsd.edu/ProteoSAFe/status.jsp?task=6cbeb7090dc74412a1461489bb037cc4

**Additional Data Analysis**
**MassQL Sandbox Visualization**

https://msql.ucsd.edu/?query=QUERY%20scaninfo(MS2DATA)%20WHERE%0AMS2PROD%3D201.054%3ATOLERANCEPPM%3D5%3AINTENSITYPERCENT%3D1%20AND%20MS2PROD%3D135.044%3ATOLERANCEPPM%3D5%3AINTENSITYPERCENT%3D1&

**Data Availability**

MSV000078716; MSV000078917; MSV000081451; MSV000081502; MSV000083055; MSV000083571; MSV000083574; MSV000084278; MSV000084946; MSV000085070.

The Piperaceae plant family is well-known as a remarkable source of specialized metabolites with pharmacological properties, with > 300 different amide alkaloids isolated to date[40]. Among them, piperamides represent a particularly attractive category which has shown a wide spectrum of pharmacological effects, from drug-bioavailability enhancing to leishmanicidal activity[41,42]. The chemical structures of few of these compounds are shown below:

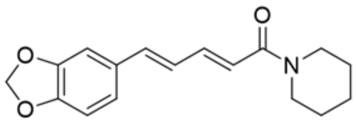

Piperine
$C_{17}H_{19}NO_3$

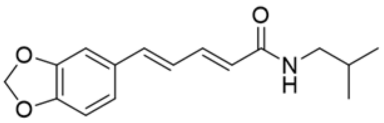

Piperlonguminine
$C_{16}H_{19}NO_3$

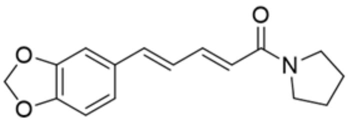

Trichostachine
$C_{16}H_{17}NO_3$

In MS/MS, piperamides produce two prominent and characteristic peaks (i.e., *m/z* = 201.054 and *m/z* = 135.044). As an example, MS/MS spectrum of piperine (Spectrum Link), as well as its molecular network (Molecular Networking Link), is reported below in **SI Fig. 2.2.1 - 1**:

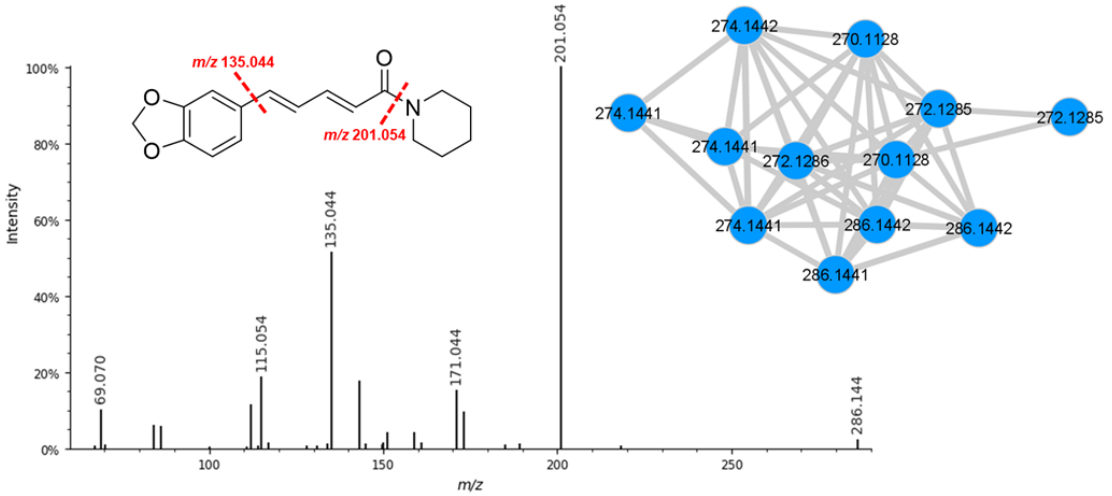


**SI Figure 2.2.1 -1** - MS/MS spectrum of piperine and example of molecular network of piperamide compounds.

A sample set of 27 *Piper* plant species was analyzed on an Orbitrap ID-X Tribrid mass spectrometer (Thermo Fisher Scientific, Waltham, MA) using a DDA mode. Data is publicly accessible at MSV000087894. The goal of the analysis was to screen for the presence of piperamides and map their inter-species distribution. We first used MassQL to search MS2 containing both the characteristic fragment ions.

The MassQL query returned 982 scans that fulfilled the criteria in 25 plant samples. As expected, several noisy spectra and spectra clearly not belonging to piperamides (e.g., Spectrum Link) were found. Nevertheless, we were able to easily and effectively refine the MassQL query by searching for MS2 scans where one of the two diagnostic fragments is the most intense in the spectrum.

##### 2.2.2 Discovery of Novel Piperamides in *Piperaceae* plants using MassQL: Diagnostic Ions Refined

**Author/s: Tito Damiani**

**MassQL Query**

```
QUERY scaninfo(MS2DATA) WHERE
MS2PROD=201.054:TOLERANCEPPM=5:INTENSITYPERCENT=100 AND
MS2PROD=135.044:TOLERANCEPPM=5
|||
QUERY scaninfo(MS2DATA) WHERE
MS2PROD=201.054:TOLERANCEPPM=5 AND
MS2PROD=135.044:TOLERANCEPPM=5:INTENSITYPERCENT=100
```

**MassQL Translation**

```
Returning the scan information on MS2.
The following conditions are applied to find scans in the mass spec data.
Finding MS2 peak at m/z 201.054 a 5.0 PPM tolerance and a minimum percent
intensity relative to base peak of 100.0%.
Finding MS2 peak at m/z 135.044 a 5.0 PPM tolerance.

Returning the scan information on MS2.
The following conditions are applied to find scans in the mass spec data.
Finding MS2 peak at m/z 201.054 a 5.0 PPM tolerance.
Finding MS2 peak at m/z 135.044 a 5.0 PPM tolerance and a minimum percent
intensity relative to base peak of 100.0%.
```

**MassQL Query Link**

https://proteomics2.ucsd.edu/ProteoSAFe/status.jsp?task=25a8465d49e3497e85a546c11849c4c9

**Data Availability**

MSV000078716; MSV000078917; MSV000081451; MSV000081502; MSV000083055; MSV000083571; MSV000083574; MSV000084278; MSV000084946; MSV000085070.

The new query string dramatically reduced the number of hits, from 982 to 247 MS2 scans meeting the query criteria. Manual inspection of the results revealed MS2 spectra clearly related to piperamides which were, however, not part of the molecular network due to a low cosine similarity (cosine threshold = 0.6). For instance, a MS2 spectra (precursor *m/z* = 543.2487 - Spectrum Link) was found to contain a product ion with *m/z* = 272.1284, which corresponds to the $[M+H]^+$ adduct of trichostachine. The substantial spectral similarity, along with available background knowledge, led us to putatively assign *m/z* = 543.2487 to a trichostachine dimer. Although piperamides dimers have already been reported in literature[40], to the best of our knowledge, trichostachine dimers have never been characterized thus far. MS2 spectra of trichostachine and the putative dimer are shown below in a mirror view (Mirror Plot Link):

It must be noted that such metabolite was not connected to a node because of the low similarity score (i.e., 0.57) and, therefore, it would have likely been missed by applying only molecular networking for data investigation. In conclusion, the present use case demonstrates how MassQL constitutes an additional means for MS data mining to be used alongside the existing tools.

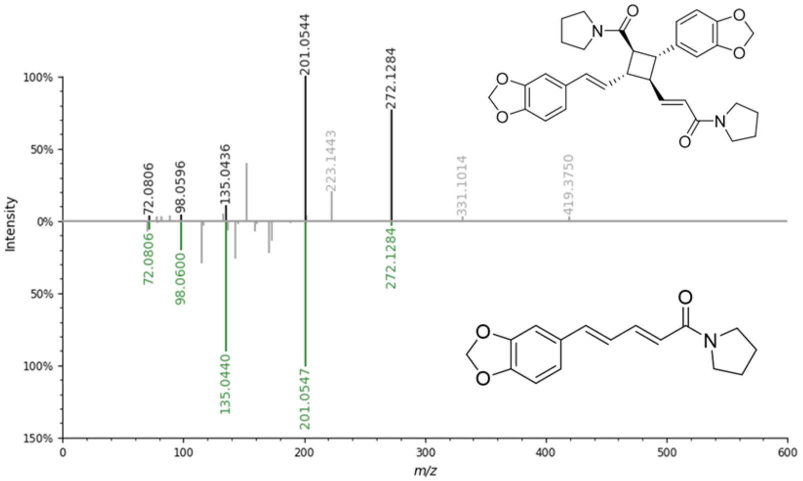


**SI Figure 2.2.2 - 1** - MS/MS spectra of trichostachine (bottom) and putative trichostachine dimer (top) shown as a mirror plot.

#### 2.3 Searching for Glycoalkaloids from *Solanum* species using MassQL

**Author/s: Ricardo Moreira Borges**

| **MassQL Query** |
| --- |
| `QUERY scaninfo(MS2DATA) WHERE MS2PROD=396.32:TOLERANCEMZ=0.1 AND MS2PROD=253.19:TOLERANCEMZ=0.1 \|\|\| QUERY scaninfo(MS2DATA) WHERE MS2PROD=273.22:TOLERANCEMZ=0.1 AND MS2PROD=255.21:TOLERANCEMZ=0.1` |
| **MassQL Query Link**<br>https://proteomics2.ucsd.edu/ProteoSAFe/status.jsp?task=abde4e2dd65a4ea496827434ad407dd7 |
| **Additional Data Analysis**<br>**GNPS Molecular Networking**<br>https://proteomics2.ucsd.edu/ProteoSAFe/status.jsp?task=a5c3f79864e843fb833abc5371e067c5 |
| **Data Availability**<br><br>MSV000087711 |

Glycoalkaloids are a subclass of saponins that contains a nitrogen atom in its aglycone moiety naturally occurring in various plant species of *Solanum* (Solanaceae)[43,44]. Their importance is exemplified by its potential toxicity to humans in high levels and their production in several popular food sources, such as potato, tomato, aubergines, etc. Hence, the need for faster and comprehensive approaches for screening glycoalkaloids in food samples is desirable and highly sought after. Compared to other more well-known classes of natural products, glycoalkaloids are less investigated; consequently, far fewer representatives of this class are usually found in databases for compound identification.

Not surprisingly, this MassQL based approach has a parallel with features that characterize a Mass2Motif from the MS2LDA method. In this case, the key fragments used as queries (at *m/z* 396.32 and *m/z* 253.19 and also *m/z* 273.22 and *m/z* 255.21) which represents some of the diagnostic fragments for a steroid-like structure is also found when applying a similar dataset in MS2LDA.

Throughout this application, we were able to filter scans successfully annotated as glycoalkaloids, and others annotated as the aglycone (**SI Fig. 2.3 - 1, 2.3 - 2**). The sequential loss of sugar residues were not explored as queries since they can vary for different compounds, but they can be visualized through manual validation.

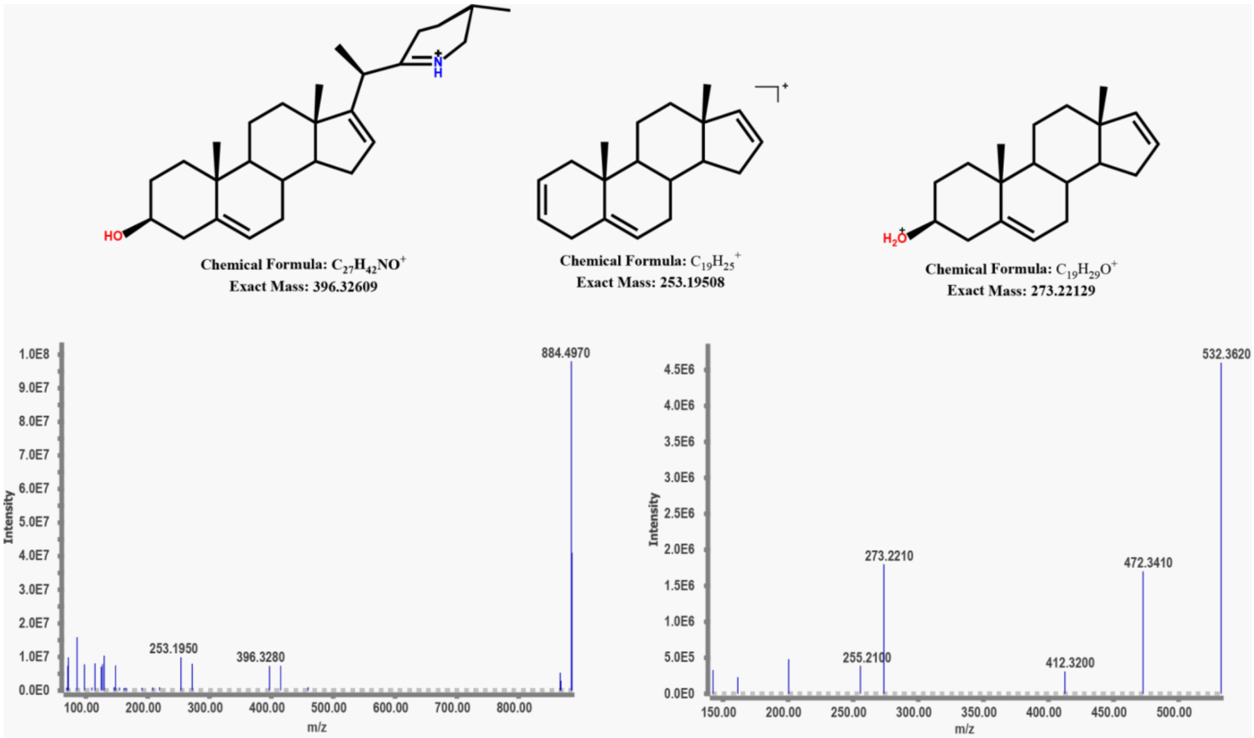


**SI Figure 2.3 - 1** - MS/MS spectra of different glycoalkaloids sharing similar steroid-like aglycone.

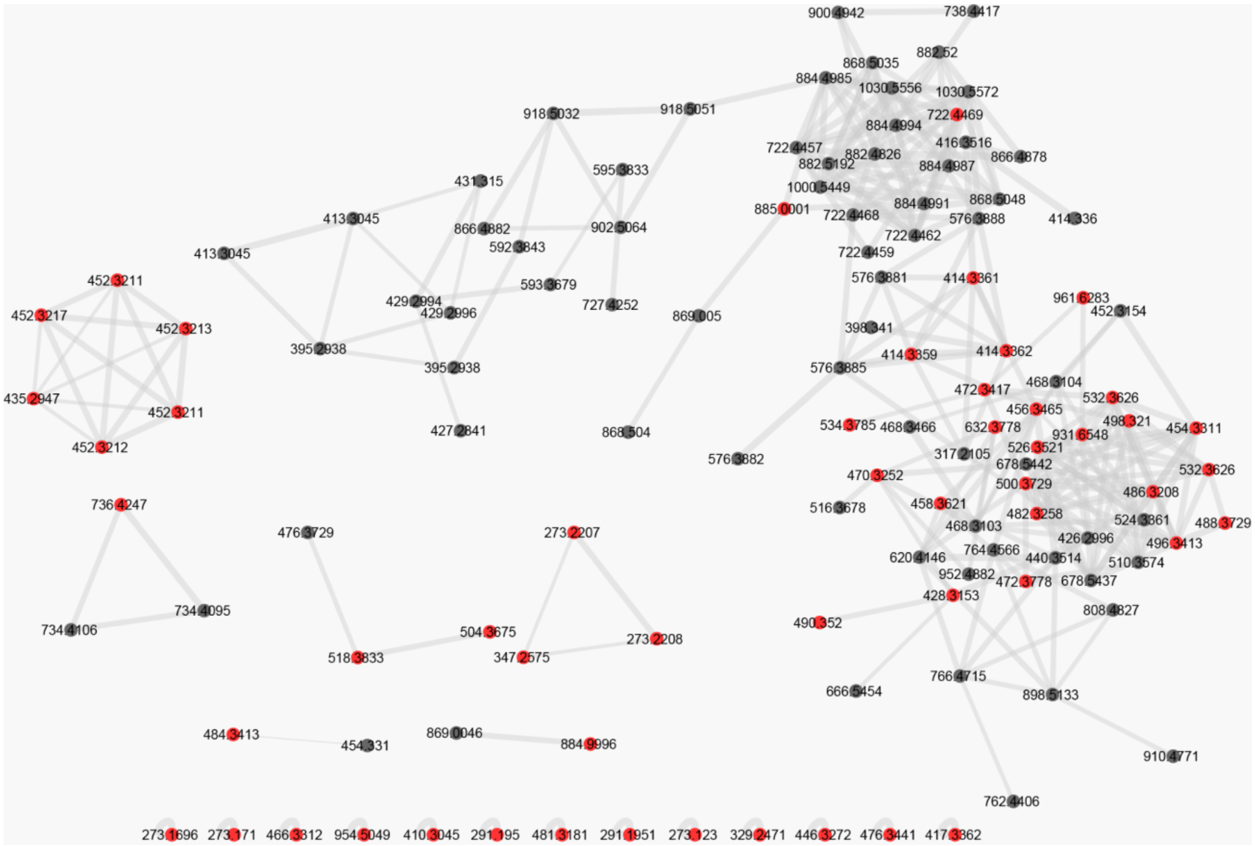

**SI Figure 2.3 - 2** : Molecular networks with nodes identified by the query as glycoalkaloids highlighted in red.

#### 2.4 Search for Sources of Compounds of Interest for SARS-CoV-2

**Author/s: Ricardo Moreira Borges and Fernanda Oliveira Chagas**

| **MassQL Query** |
| --- |
| `QUERY scaninfo(MS2DATA) WHERE MS1MZ=443.16:TOLERANCEMZ=0.1 AND MS2PREC=443.16:TOLERANCEMZ=0.1 AND MS2PROD=249.06 ||| QUERY scaninfo(MS2DATA) WHERE MS1MZ=437.14:TOLERANCEMZ=0.1 AND MS2PREC=437.14:TOLERANCEMZ=0.1 AND MS2PROD=270.12 ||| QUERY scaninfo(MS2DATA) WHERE MS1MZ=441.19:TOLERANCEMZ=0.1 AND MS2PREC=441.19:TOLERANCEMZ=0.1 AND MS2PROD=312.11 ||| QUERY scaninfo(MS2DATA) WHERE MS1MZ=356.10:TOLERANCEMZ=0.1 AND MS2PREC=356.10:TOLERANCEMZ=0.1 AND MS2PROD=338.09 ||| QUERY scaninfo(MS2DATA) WHERE MS1MZ=461.15:TOLERANCEMZ=0.1 AND MS2PREC=461.15:TOLERANCEMZ=0.1 AND MS2PROD=443.14 ||| QUERY scaninfo(MS2DATA) WHERE MS1MZ=301.07:TOLERANCEMZ=0.1 AND MS2PREC=301.07:TOLERANCEMZ=0.1 AND MS2PROD=283.06` |
| **MassQL Query Link**<br>https://proteomics2.ucsd.edu/ProteoSAFe/status.jsp?task=b3233f7bc1ba4ab295a387f7953e2063 |
| **Data Availability**<br>MSV000078716; MSV000078917; MSV000081451; MSV000081502; MSV000083055; MSV000083571; MSV000083574; MSV000084278; MSV000084946; MSV000085070. |

*In silico* tools have been used to filter chemical structure databases attempting to point out promising compounds that might interact with specific enzymes, but the link to the organism that produces these promising compounds is often missing. In this context, a MS/MS query approach can be resumed within a pipeline to enable the search for compounds of interest among data from repository such as MassIVE/GNPS which, because it has an organized metadata, yield useful information regarding organism, sample preparation, and leading research group. MassQL is a promising tool to link both *in silico* derived compounds of interest to search in data repositories.

Here, we briefly line up the pipeline using the search for potential anti-SARS-CoV-2 drug candidates as an example. The following steps were used: (1) The reference[45] suggested six compounds as hit; (2) the MS/MS spectra of those six compounds were predicted using CFM-ID[46] 3.0 (https://cfmid.wishartlab.com/predict) for ESI ionization and positive mode ($[M+H]^+$); (3) diagnostic fragments were selected manually; (4) specific queries for each of the six hits were developed using the expected precursor *m/z* (in MS1) and the diagnostic *m/z* (in MS2). The queries were submitted to selected data from MassIVE: MSV000078716; MSV000078917; MSV000081451; MSV000081502; MSV000083055; MSV000083571; MSV000083574; MSV000084278; MSV000084946; MSV000085070.

The extracted scans can be manually inspected and the metadata from the MassIVE indicates the organisms where those queried spectra were detected.

#### 2.5 Homoserine Lactone

**Author/s:** **Ben Bowen and Trent Northen**

**MassQL Query**

```
QUERY scaninfo(MS2DATA) WHERE
MS2PROD=74.061:INTENSITYPERCENT=5:TOLERANCEMZ=0.005 AND
MS2PROD=84.045:INTENSITYPERCENT=5:TOLERANCEMZ=0.005 AND
MS2PROD=102.055:INTENSITYPERCENT=50:TOLERANCEMZ=0.005 AND
MS2PREC=300:TOLERANCEMZ=100
```

**MassQL Translation**

```
Returning the scan information on MS2.
The following conditions are applied to find scans in the mass spec data.
Finding MS2 peak at m/z 74.061 a minimum percent intensity relative to base
peak of 5.0% and a 0.005 m/z tolerance.
Finding MS2 peak at m/z 84.045 a minimum percent intensity relative to base
peak of 5.0% and a 0.005 m/z tolerance.
Finding MS2 peak at m/z 102.055 a minimum percent intensity relative to base
peak of 50.0% and a 0.005 m/z tolerance.
Finding MS2 spectra with a precursor m/z 300.0 a 100.0 m/z tolerance.
```

**MassQL Query Link**

https://msql.ucsd.edu//?query=QUERY+scaninfo%28MS2DATA%29+WHERE+%0AMS2PROD%3D74.061%3AINTENSITYPERCENT%3D5%3ATOLERANCEMZ%3D0.005+AND+%0AMS2PROD%3D84.045%3AINTENSITYPERCENT%3D5%3ATOLERANCEMZ%3D0.005+AND+%0AMS2PROD%3D102.055%3AINTENSITYPERCENT%3D50%3ATOLERANCEMZ%3D0.005+AND%0AMS2PREC%3D300%3ATOLERANCEMZ%3D100%0A&filename=ALL_GNPS.json&x_axis=&y_axis=&facet_column=&scan=&x_value=500&y_value=1&ms1_usi=&ms2_usi=

**Additional Data Analysis**

n/a

**Data Availability**

Public GNPS Libraries - https://gnps-external.ucsd.edu/gnpslibrary

**Introduction.** This query identifies N-acyl L-homoserine lactones (AHL). These molecules are an important class of signaling molecules involved in bacterial quorum sensing. This query identified AHLs with a range of acyl chain lengths and allows for both isothiocyanate modification and carbonyl modifications making it a nice demonstration of MassQL capabilities[47]. The below queries and results have demonstrated the common ions of these compounds are sufficient to identify them and validated this approach on a variety of standards. Here, the relative abundance of lactone ring fragments that enabled identification of AHL standards were found to be 102.055, 84.045, 74.061, and 56.050 *m/z*.

**Discussion.** We found that the query above produced excellent results against the GNPS public spectral libraries. As can be seen in the following table, all of the hits returned are AHL structures. The difference between this query and the parameters identified by Patel *et al* is the omission of the *m/z*=56 fragment ion from the query. The three ions used in this query are present in all 31 AHL structures in the NIST20 positive mode MS/MS spectral library and were sufficient in identifying AHL compounds.

| scan | precmz | Compound_Name | Adduct | library |
|---|---|---|---|---|
| CCMSLIB00000221300 | 220.082 | ReSpect:PT106400 O-Succinyl-L- | [M+H] | RESPECT |
| CCMSLIB00000223125 | 220.082 | Massbank:PR100299 O-Succinyl-L | [M+H]+ | MASSBANK |
| CCMSLIB00005720378 | 220.082 | O-SUCCINYL-L-HOMOSERINE | [M+H]+ | PSU-MSMLS |
| CCMSLIB00005723676 | 355.168 | ITC-12 | M+H | GNPS-LIBRARY |
| CCMSLIB00005723689 | 228.123 | N-(3-oxoheptanoyl)-L-homoserin | M+H | GNPS-LIBRARY |
| CCMSLIB00005723690 | 256.154 | 3-oxo-C9-HSL | M+H | GNPS-LIBRARY |
| CCMSLIB00005723691 | 284.185 | 3-oxo-C11-HSL | M+H | GNPS-LIBRARY |
| CCMSLIB00005723692 | 312.217 | 3-oxo-C13-HSL | M+H | GNPS-LIBRARY |
| CCMSLIB00005723693 | 326.233 | N-(3-Oxotetradecanoyl)-L-homos | M+H | GNPS-LIBRARY |
| CCMSLIB00005723698 | 214.107 | N-(3-Oxohexanoyl)-L-homoserine | M+H | GNPS-LIBRARY |
| CCMSLIB00005723699 | 242.138 | N-(3-Oxooctanoyl)-L-homoserine | M+H | GNPS-LIBRARY |
| CCMSLIB00005737983 | 200.128 | Massbank:RP020202 C6-homoserin | M+H | MASSBANK |

| | | | | |
|---|---|---|---|---|
| CCMSLIB00005738414 | 242.139 | Massbank:RP020902 3-oxo-C8-hom | M+H | MASSBANK |
| CCMSLIB00005738685 | 300.217 | Massbank:RP021702 3-hydroxy-C1 | M+H | MASSBANK |
| CCMSLIB00005759668 | 220.082 | Massbank:PT106400 O-Succinylho | M+H | MASSBANK |
| CCMSLIB00005883959 | 220.082 | O-SUCCINYL-L-HOMOSERINE - 40.0 | M+H | GNPS-LIBRARY |
| CCMSLIB00005883960 | 220.082 | O-SUCCINYL-L-HOMOSERINE - 50.0 | M+H | GNPS-LIBRARY |
| CCMSLIB00005883961 | 220.082 | O-SUCCINYL-L-HOMOSERINE - 60.0 | M+H | GNPS-LIBRARY |

#### 2.6 Screening Chemical Diversity of Peptide Natural Products

**Author/s:** Scott A. Jarmusch

Siderophores are iron-chelating secondary metabolites produced via nonribosomal peptide-synthetase (NRPS). Due to their oligomeric assemblage and peptidic nature, predicting fragmentation patterns of these metabolites is more straightforward than other secondary metabolites. Molecular networking has been used several times for the observation and continued discovery of siderophores, like desferrioxamines. However, without anchor points containing annotations, discovering further chemical space is difficult. In a previous study, we utilized the combination of molecular networking and MS2LDA to evaluate desferrioxamine chemical space[48]. Similar to manually annotating MS/MS fingerprints in MS2LDA, I queried only a single file from the public dataset, using neutral loss queries on desferrioxamine building blocks.

| |
|---|
| **MassQL Query**<br>`See Table 1 for additional queries:`<br>`QUERY scaninfo(MS2DATA) WHERE MS2NL=160.1206:TOLERANCEMZ=0.005 AND MS2NL=82` |
| **MassQL Translation**<br>`Returning the scan information on MS2.`<br>`The following conditions are applied to find scans in the mass spec data.`<br>`Finding MS2 neutral loss product of 160.1206 Da with a 0.005 m/z tolerance also with a MS2 neutral loss product of 82 Da.` |
| **MassQL Query Link**<br>See Table |
| **Additional Data Analysis**<br>MSMS Library Searches available in Table 1. |
| **Data Availability**<br>MSV000085618 |

The results from these various queries can be seen in Table 1 and they highlight the power of being able to tailor each workflow to different neutral loss parameters. In order to yield narrower results, each query also contains the requirement to find a second neutral loss based on desferrioxamine structure and biosynthesis. From a single file in the public dataset, numerous desferrioxamine derivatives are pulled from the data and additional library searches were run on each search. Pulling from the *N*-hydroxycadaverine and succinate search, library matches return hits on desferrioxamine B and its derivatives, which is expected. Derivatives like desferrioxamine D1 which contain these same moieties internally instead of at the C-terminus are returned in the search. In addition to locating knowns and new derivatives of known metabolites, this tool may also allow for discovery of unknowns with similar moieties that do not link via molecular networking. We envision that most peptide natural products can be queried in a similar fashion to facilitate fast identification of metabolites with specific moieties without requirement to manually search raw data.

| **MS2NL input** | **# of features detected** | **MassQL Query Link** | **MSMS Library Search** | **# of annotations** |
|---|---|---|---|---|
| 160.1206 AND 82 | 25 | https://proteomics2.ucsd.edu/ProteoSAFe/status.jsp?task=e38afea27ec9483a887bab266aa2e1e2 | *https://gnps.ucsd.edu/ProteoSAFe/status.jsp?task=a13be8c9391e414aba6bd693a7f5517a* | 16 |
| 118.1101 AND 82 | 31 | https://proteomics2.ucsd.edu/ProteoSAFe/status.jsp?task=88f92361d5ce419a913497c35d09ba19 | *https://gnps.ucsd.edu/ProteoSAFe/status.jsp?task=d80f4e61dd9742cba2e08f598b70d3f1* | 24 |
| 104.0944 AND 82 | 9 | https://proteomics2.ucsd.edu/ProteoSAFe/status.jsp?task=deed922a50f44174bd22b74c7315ef0a | *https://gnps.ucsd.edu/ProteoSAFe/status.jsp?task=764d18af0cbf4e7189e4d780808d512c* | 5 |
| 90.0788 AND 82 | 6 | https://proteomics2.ucsd.edu/ProteoSAFe/status.jsp?task=5a95f9f31165484fad0ec710669f42f5 | *https://gnps.ucsd.edu/ProteoSAFe/status.jsp?task=b7e4203df5ae49dd90cf3571aec86eeb* | 4 |
| 102.1157 AND 82 | 5 | https://proteomics2.ucsd.edu/ProteoSAFe/status.jsp?task=5b2a86dd4e944b38a35ead08428a4935 | https://gnps.ucsd.edu/ProteoSAFe/status.jsp?task=568ebbd72eb54fe899d95009878fdb93 | 4 |
| 88.0995 AND 82 | 2 | https://proteomics2.ucsd.edu/ProteoSAFe/status.jsp?task=39cfd59bf6c14d9dbed9c287dbc0b0ab | https://gnps.ucsd.edu/ProteoSAFe/status.jsp?task=c66d96dc3b9747eb946def9219f0c73b | 2 |

#### 2.7 MassQL to Assess Mass2Motif Substructure Patterns and Annotate Mass2Motif Substructure Patterns: Acylcarnitines as a Case Study

**Author/s: Justin J.J. van der Hooft**

**Motivation.** This example of the acylcarnitine demonstrates how with MassQL we could assess the specificity of a previously defined acylcarnitine filter in modern LC-MS/MS profiles. It also shows how MassQL is able to successfully group a biochemically relevant compound class that is difficult to capture using molecular networking alone.

**Introduction.** Mass2Motif substructure patterns discovered by MS2LDA[6] can consist of a number of mass fragments and neutral losses that together describe the presence of one or several related substructures. The decomposition of complex metabolite mixtures into substructures and connected metabolites has proven to be useful[6,49–52]. However, a remaining challenge is the structural annotation of the resulting fragmentation patterns, in particular those including neutral losses, in the absence of easy ways of querying large amounts of mass spectra with such spectral features. The development of MassQL makes it possible to interact with MS/MS data in an easy manner. Furthermore, in the future more complex queries can be imagined that will allow integration with LC-MSn data as well, or facilitate more specific scaffold hunting than is currently possible. In particular, the facilitated querying of both mass fragments and neutral losses represents a major step forward in assessing how specific Mass2Motif patterns are to describe a particular compound class.

In previous work, a manually constructed acylcarnitine mass fragment and neutral loss filter was established based on urine LC-MS/MS data[53]. This chemical pattern resulted in the discovery of many previously uncharacterized acylcarnitine species. Acylcarnitines can play important roles as biomarkers for diseases such as mitochondrial disorders or autism[54]. In later work, various acylcarnitine-related Mass2Motifs were annotated in urine samples[49]. Furthermore, the technological advances also enabled mass spectrometers to fragment many more metabolite features during a single LC-MS/MS run, thus increasing the coverage of the urine metabolome and raising the question whether the original acylcarnitine screening filter would remain sufficiently specific. It is therefore interesting to assess the specificity of the original acylcarnitine mass fragment and neutral loss filter using MassQL.

**Results.** Here, we took 11 urine LC-MS/MS files acquired in positive ionization mode available from Van der Hooft *et al.*, 2016[55] and queried them with an increasing number of mass fragments and neutral losses to assess the acylcarnitine specificity of selected mass spectra. To aid this process, a subset of ~10,000 GNPS library spectra was also queried using the same MassQL queries (see below). Starting with just using the mass fragment of *m/z* 85.0284 (with a tolerance that reflects the measurement on an Orbitrap instrument), on average 1161 mass spectra returned from the MassQL query. Whilst we expected many acylcarnitine species, this number was really quite high. Browsing through the list, it quickly became obvious that many non-acylcarnitine related metabolite features were included. Adding the neutral loss of 59.0735 Da to the MassQL query (making it effectively the same as the originally proposed acylcarnitine filter) reduced this number to 268 mass spectra on average per LC-MS/MS urine profile. Prompted by these results, we decided to run the same MassQL queries against the GNPS library to assess the specificity of the filter there. This query led to 869 and 65 returned mass spectra, respectively. The combined mass fragment and neutral loss filter contained 20 false positives and 45 carnitine-related spectra. Therefore, inspired by the acylcarnitine-related Mass2Motifs previously found, another mass fragment was added to the MassQL query: 60.0825 *m/z*. This query returned 42 mass spectra from the library with only 1 false positive. In the urine profiles, on average 245 mass spectra were found using the extended filter, and based on the library results, it is expected that the majority are acylcarnitine-related. It is important to note that in the GNPS library multiple spectra of the same metabolite are present, and that in the LC-MS/MS profiles, multiple mass spectra can be obtained of the same metabolite feature.

To further investigate the MassQL results, molecular networking experiments were done. Mostly regular differences in parent masses are expected due to different lengths and modifications of the acyl chains, i.e., hydroxylations and methylations, etc. (see **SI Fig. 2.7 - 1-6**). Indeed, upon adding additional features to the screening filter, the regular pattern emerges in the figures. It also became clear that most acylcarnitines do not cluster together in molecular networking, despite putting relatively low threshold values for the edges: the zero mass difference is mostly populated in the most strict filter. Inspection of the networks reveals that the same or very close (isomeric) analogs do cluster in one molecular family, but the MS/MS spectra do differ too much to match most variable acylcarnitine species. Hence, MassQL and Mass2Motifs offer an attractive and powerful alternative to group biochemically relevant metabolites.

**Discussion.** We show how MassQL has the power to effectively select and map acylcarnitines from a large dataset as an alternative to annotation of molecular families using molecular networking in combination with library matching that provides seed node annotations. In the future, a connection to MotifDB[50] could be established where MassQL-inspired substructure patterns could become annotated Mass2Motifs in a MotifDB MotifSet. In addition, querying the presence of mass differences will allow the user to further specify their queries. This example underlines the complementarity of the unsupervised MS2LDA and the supervised MassQL and how they can be used in tandem to enhance mining of mass spectrometry data.

**MassQL Queries**

| **MassQL Query**<br>`QUERY scannum(MS2DATA) WHERE MS2PROD=85.0284:TOLERANCEMZ=0.005` |
|---|
| **MassQL Translation**<br>`Finding MS2 spectra scan number.`<br>`The following conditions are applied to find scans in the mass spec data.`<br>`Finding MS2 peak at m/z 85.0284 with a 0.005 m/z tolerance.` |
| **MassQL Query Link**<br>https://proteomics2.ucsd.edu/ProteoSAFe/status.jsp?task=a2c881737c244ab597cfe387b11ffea3 |

| **MassQL Query**<br>`QUERY scannum(MS2DATA) WHERE MS2PROD=85.0284:TOLERANCEMZ=0.005 AND MS2NL=59.0735:TOLERANCEMZ=0.005` |
|---|
| **MassQL Translation**<br>`Finding MS2 spectra scan number.`<br>`The following conditions are applied to find scans in the mass spec data.`<br>`Finding MS2 peak at m/z 85.0284 with a 0.005 m/z tolerance.`<br>`Finding MS2 neutral loss peak at m/z 59.0735 with a 0.005 m/z tolerance.` |
| **MassQL Query Link**<br>https://proteomics2.ucsd.edu/ProteoSAFe/status.jsp?task=0b20723dde2648058faf10c07ebaa6c9 |

| **MassQL Query**<br>`QUERY scannum(MS2DATA) WHERE MS2PROD=85.0284:TOLERANCEMZ=0.005 AND MS2PROD=60.0825:TOLERANCEMZ=0.005 AND MS2NL=59.0735:TOLERANCEMZ=0.005` |
|---|
| **MassQL Translation** |

```
Finding MS2 spectra scan number.
The following conditions are applied to find scans in the mass spec data.
Finding MS2 peak at m/z 85.0284 with a 0.005 m/z tolerance.
Finding MS2 peak at m/z 60.0825 with a 0.005 m/z tolerance.
Finding MS2 neutral loss peak at m/z 59.0735 with a 0.005 m/z tolerance.
```

**MassQL Query Link**
https://proteomics2.ucsd.edu/ProteoSAFe/status.jsp?task=bc72cf62c2654d10a766d88d79dd5749

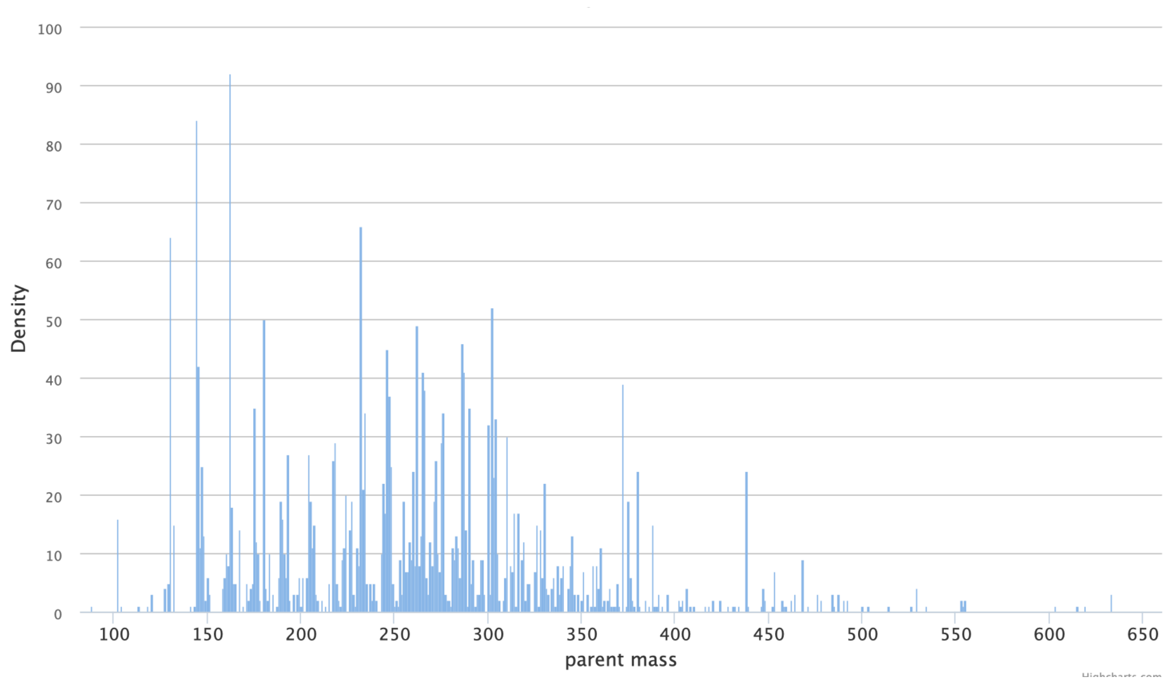


**SI Figure 2.7 - 1**. Density plot of parent masses in molecular network of selected spectra in urine data based on Filter 1 (85 mass fragment). The consensus spectra with parent masses are resulting from MSClust preprocessing that is part of the classical molecular networking job workflow. MN Link

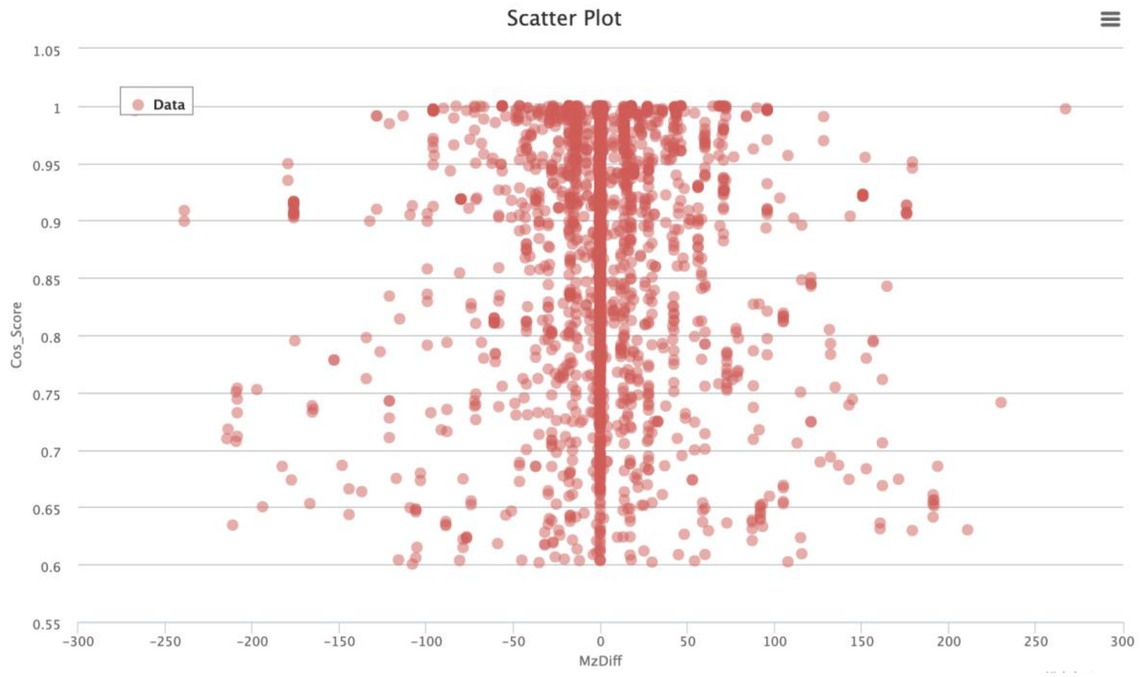


**SI Figure 2.7 - 2**. Scatter plot of cosine scores versus mass differences found in molecular network of selected spectra in urine data based on Filter 1 (85 mass fragment). MN Link

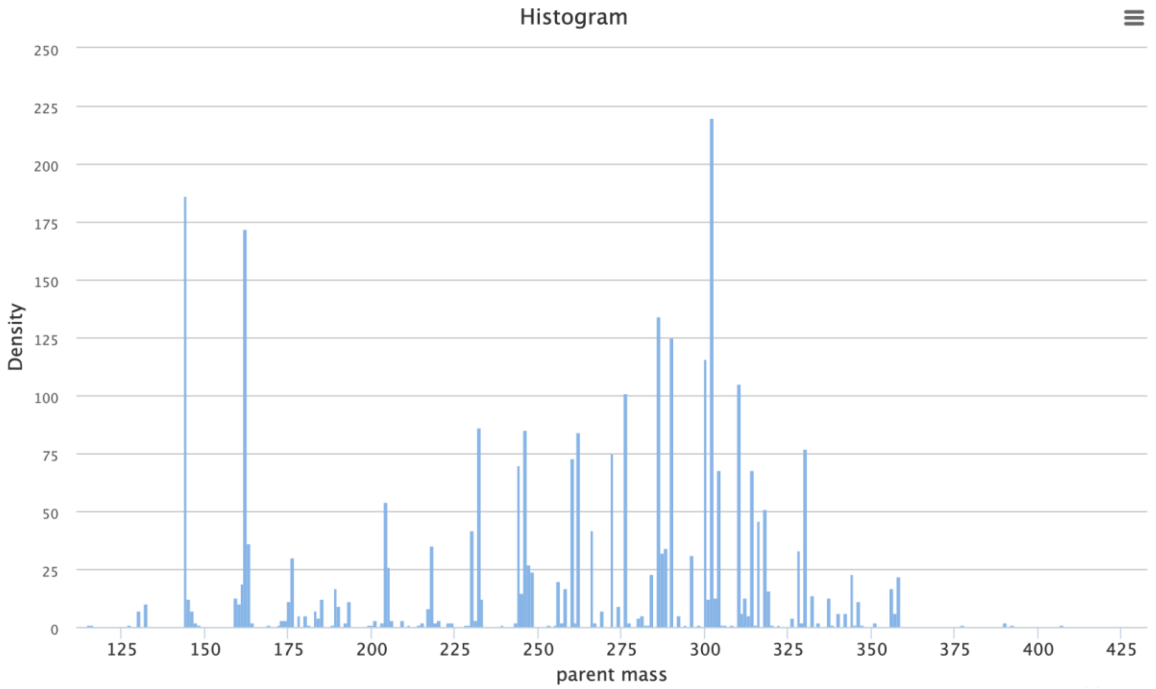


**SI Figure 2.7 - 3**. Density plot of parent masses in molecular network of selected spectra in urine data based on Filter 2  (85 mass fragment and 59 neutral loss). The consensus spectra with parent masses are resulting from MSClust preprocessing that is part of the classical molecular networking job workflow. MN Link

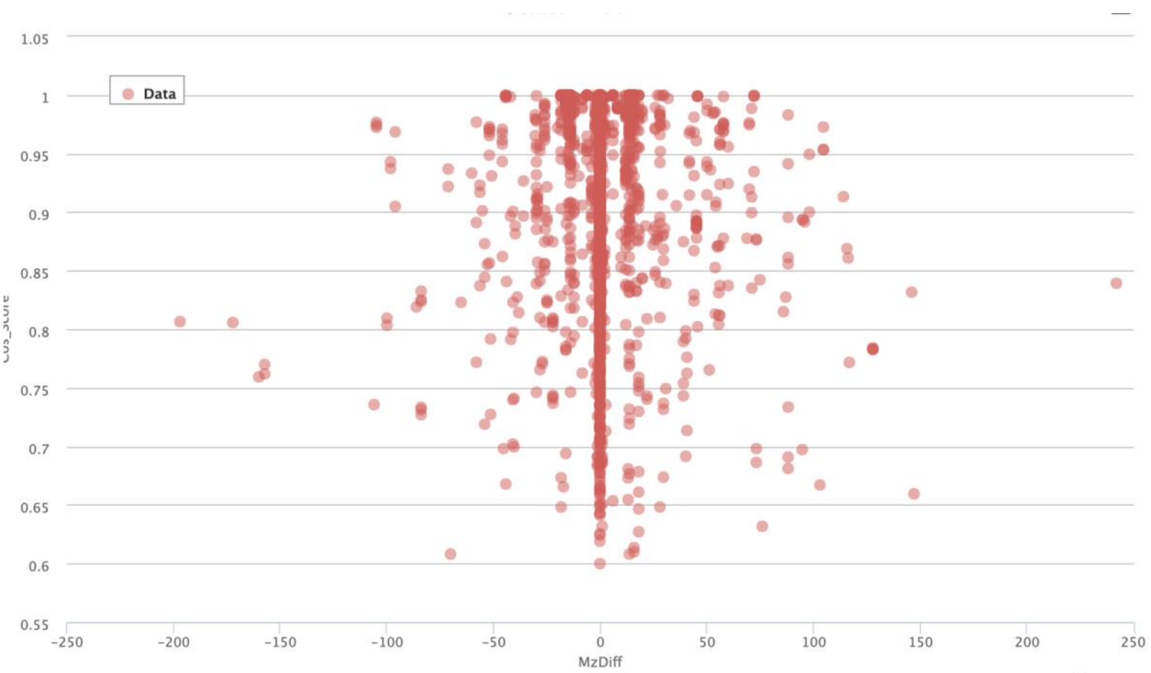


**SI Figure 2.7 - 4**. Scatter plot of cosine scores versus mass differences found in molecular network of selected spectra in urine data based on Filter 2 (85 mass fragment and 59 neutral loss). MN Link

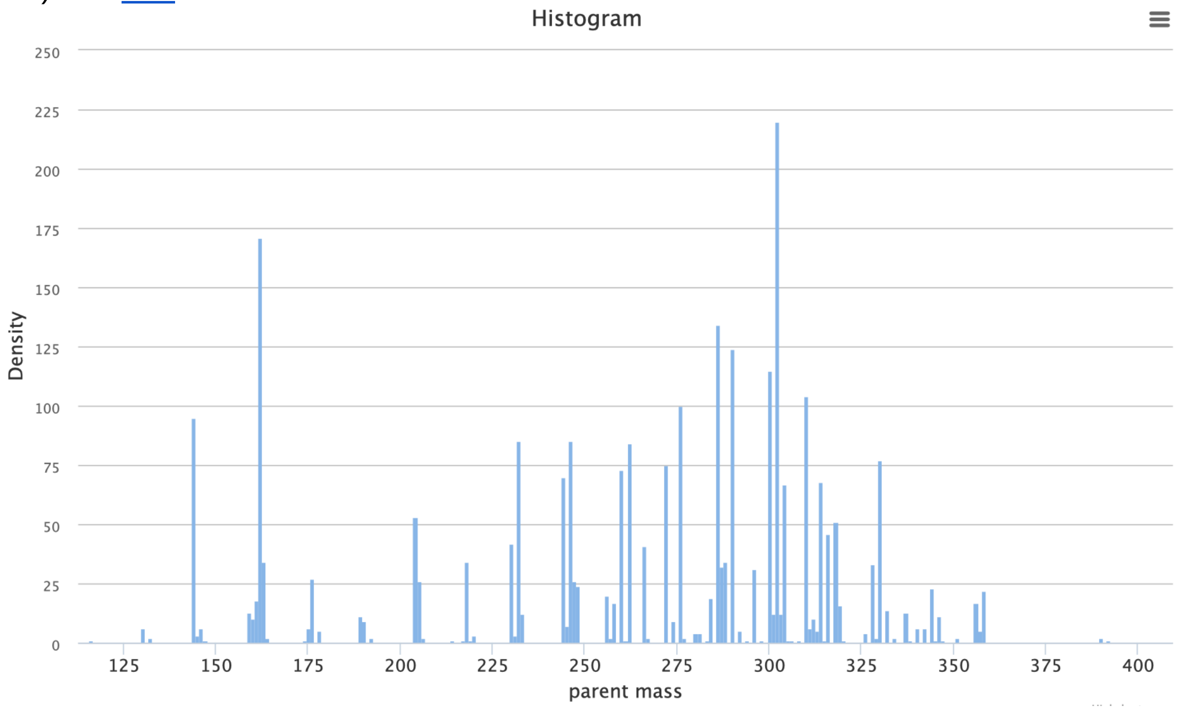


**SI Figure 2.7 - 5.** Density plot of parent masses in molecular network of selected spectra in urine data based on Filter 3 (85 and 60 mass fragments and 59 neutral loss). The consensus spectra with parent masses are resulting from MSClust preprocessing that is part of the classical molecular networking job workflow. MN Link

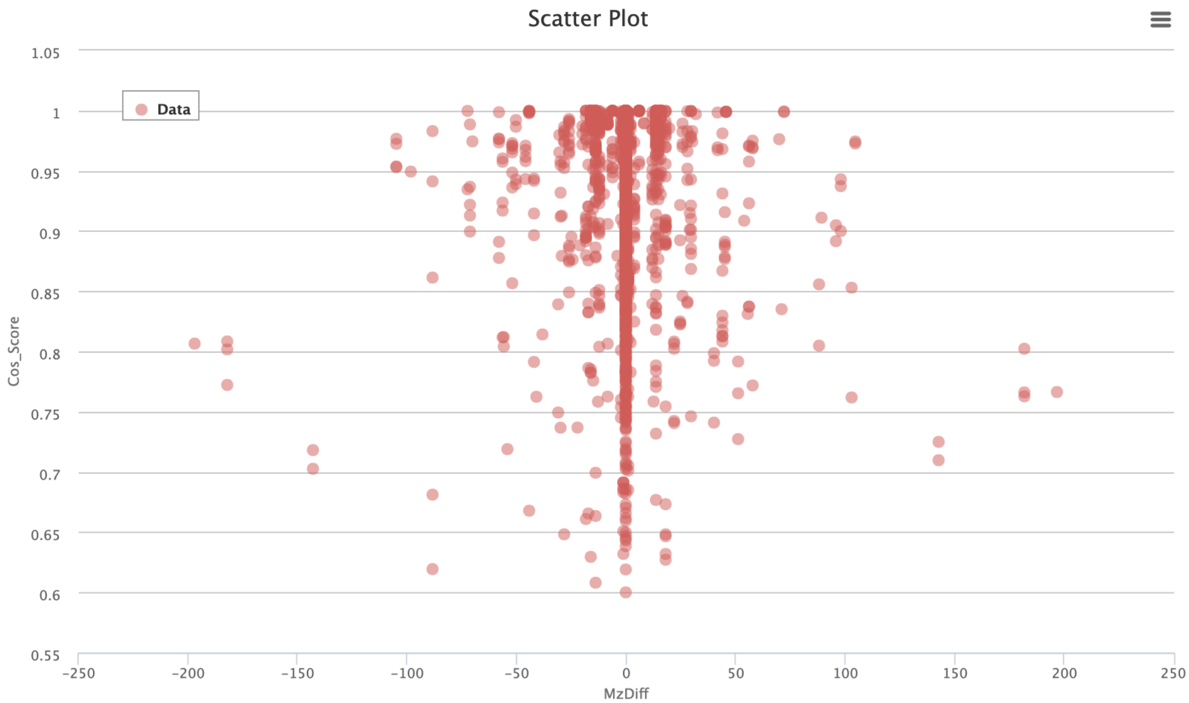


**SI Figure 2.7 - 6**. Scatter plot of cosine scores versus mass differences found in molecular network of selected spectra in urine data based on Filter 3 (85 and 60 mass fragments and 59 neutral loss). MN Link

Experimental data used for this case study is available at MSV000082971.

The GNPS spectral libraries can be found here https://gnps.ucsd.edu/ProteoSAFe/libraries.jsp and were queried through https://msql.ucsd.edu/.

**2.8 - MassQL for the Distinction of C-hexoglycosides, O-hexoglycosides and Pentoglycosides**

**Author/s: Elys Rodriguez and Oliver Fiehn**

**MassQL Query**

**For O-hexoglycosidase**

QUERY scaninfo(MS2DATA) WHERE MS2PROD=162.05 (for galactose or glucose)

QUERY scaninfo(MS2DATA) WHERE MS2PROD=176.04 (for glucuronic acid)

**For C-hexglycosides**

QUERY scaninfo(MS2DATA) WHERE MS2PROD=162.05 (glucose)

QUERY scaninfo(MS2DATA) WHERE MS2PROD=120.04 (glucose fragmentation)

QUERY scaninfo(MS2DATA) WHERE MS2PROD=90.04 (glucose fragmentation)

**For O-pentoglycosides**

QUERY scaninfo(MS2DATA) WHERE MS2PROD=132.06 (arabinose)

QUERY scaninfo(MS2DATA) WHERE MS2PROD=145.06 (rhamnose)

**MassQL Translation**

Returning the scan information on MS2.
The following conditions are applied to find scans in the mass spec data.
Finding MS2 neutral loss peak at m/z 162.05

Returning the scan information on MS2.
The following conditions are applied to find scans in the mass spec data.
Finding MS2 neutral loss peak at m/z 176.04

Returning the scan information on MS2.
The following conditions are applied to find scans in the mass spec data.
Finding MS2 neutral loss peak at m/z 162.05
Finding MS2 neutral loss peak at m/z 120.04
Finding MS2 neutral loss peak at m/z 90.04

Returning the scan information on MS2.
The following conditions are applied to find scans in the mass spec data.
Finding MS2 neutral loss peak at m/z 132.06
Finding MS2 neutral loss peak at m/z 145.06

**MassQL Query Link**

https://proteomics2.ucsd.edu/ProteoSAFe/result.jsp?task=214b22da4bfe4d0899efde9fd8f85c07&view=query_results

https://proteomics2.ucsd.edu/ProteoSAFe/result.jsp?task=78fb96a33979401db1e0cdecf53b389a&view=query_results

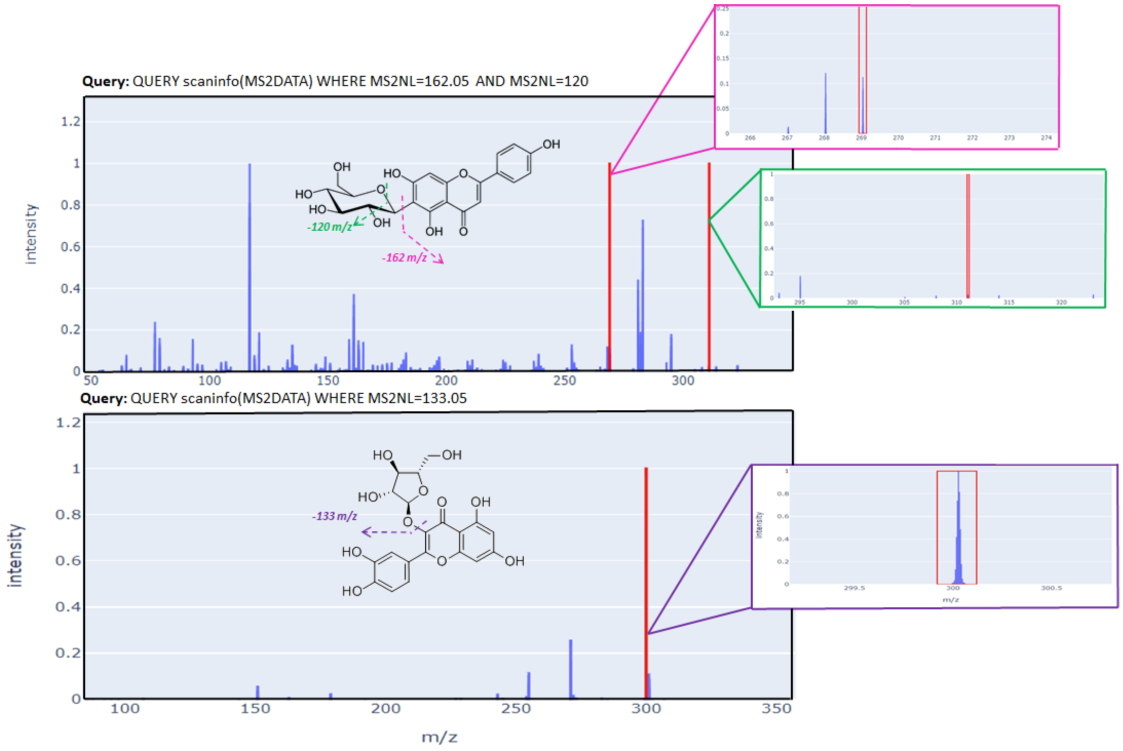


**SI Figure 2.8 - 1.** MS/MS spectra of a) Apigenin-6-C-glucoside and b) Quercetin-3-arabinopyranoside. Pink squared spectra shows -162 Da glucose loss, green squared spectra shows -120 Da glucose fragmentation and purple squared spectra shows -133 Da arabinose loss.

Glycosylation may be one of the most efficient PTM in all kingdoms of life as 1) sugar can be added to the most abundant heteroatoms (i.e., N, O, S) and carbon and 2) products may be spontaneously cleaved upon transportation to low or high pH, thus avoiding further enzymatic input. Furthermore, upon glycosylation, hydrophobic metabolites become more water-soluble which improves their bio-distribution and metabolism. Due to glycosylation's ability to sequester secondary metabolites, it is of major importance to identify glycoconjugates in order to construct an accurate atlas of secondary metabolites in nature. With the use of MassQL, the distinction of O-glucosides, O-arabinoside, and C-glucosides can be observed by simply searching for neutral loss. MassQL may be an excellent tool for untargeted analysis of glycoconjugates.

### 3 MassQL Query using MS/MS Spectral Information and MS Isotope Pattern

#### 3.1 Manual Exploration versus MassQL for Processing Untargeted Metabolomics Data: Data Exploration for Sulfur-containing Perfluorinated Compounds in Human Plasma (NIST Standard Reference Material 1950)

**Author/s: Alan K. Jarmusch and Kirsten E. Overdahl**

| |
|---|
| **MassQL Query**<br>`QUERY scaninfo(MS2DATA) WHERE`<br>`MS1MZ=X:INTENSITYPERCENT=20 AND`<br>`MS1MZ=X+1.996:TOLERANCEPPM=5 AND`<br>`MS2PROD=79.9563:TOLERANCEPPM=10:INTENSITYPERCENT=10 AND`<br>`MS2PROD=98.9552:TOLERANCEPPM=10:INTENSITYPERCENT=10` |
| **MassQL Translation**<br>`Returning the scan information on MS2.`<br>`The following conditions are applied to find scans in the mass spec data.`<br>`Finding MS1 peak at m/z X a minimum percent intensity relative to base peak of 20.0%.`<br>`Finding MS1 peak at m/z X+1.996 a 5.0 PPM tolerance.`<br>`Finding MS2 peak at m/z 79.9563 a 10.0 PPM tolerance and a minimum percent intensity relative to base peak of 10.0%.`<br>`Finding MS2 peak at m/z 98.9552 a 10.0 PPM tolerance and a minimum percent intensity relative to base peak of 10.0%.` |
| **MassQL Query Link**<br>https://proteomics2.ucsd.edu/ProteoSAFe/status.jsp?task=305e851d79a94145b63a773d217c0f58 |
| **Additional Data Analysis**<br>n/a |
| **Data Availability**<br>n/a |

Environmental health implications of per- and polyfluoroalkyl substances (PFAS) are of increasing concern. It is known that PFAS are present in drinking water and commercial products, and that they are suspected to cause reproductive, developmental, liver, and kidney damage. Perfluorooctanesulfonic acid and related perfluorinated sulfonate compounds (PFOS) have been detected in human blood[56]. However, the vast number (>5,000) of possible PFOS structures, coupled with lack of comprehensive structural or regulatory databases, presents challenges to

PFOS identification and toxicity testing. Untargeted mass spectrometry is ideally suited for exploring and putatively identifying sulfur-containing perfluorinated chemicals.

NIST Standard Reference Material 1950 (Metabolites in Frozen Human Plasma) was analyzed using reverse-phase ultra-high performance liquid chromatography (Kinetex® 2.6 µm F5 100Å 100 x 2.1 mm column, Phenomenex; Thermo-Fisher Vanquish) coupled to high resolution mass spectrometry (Thermo-Fisher Orbitrap Tribrid Fusion) with heated electrospray ionization in negative ionization mode. Separation performed with an initial isocratic condition for two minutes, 98% water with 0.1% acetic acid - 2% acetonitrile with 0.1% acetic acid (B), followed by a ten minute linear gradient to 100% B. Data-dependent MS/MS acquisition was performed via AcquireX (deep scan mode) from 5 replicate injections.

To represent a manual spectral interpretation process, one SRM 1950 replicate injection file was interpreted in FreeStyle (Thermo Scientific). Extracted ion chromatograms were generated using known PFOS precursor ion masses. The mass error, $^{34}S$ isotope pattern, and MS/MS spectra were evaluated manually. Manual interpretation yielded two reported perfluorinated sulfonates (**SI Table 3.1 - 1**): perfluorohexanesulfonic acid (PFHxS) and perfluorooctanesulfonic acid (PFOS). Characteristic product ion of *m/z* 79.9563 ($SO_3^-$) associated with sulfates was observed. The observation of *m/z* 98.9552 ($FSO_3^-$) supported the presence of at least one fluorine atom. The potential sulfonated product ion of *m/z* 96.9596 ($HSO_4^-$) was not detected in the perfluorinated sulfonate spectra, attributed to the absence of hydrogens in the perfluorinated compounds. The amount of time spent for this manual interpretation, plotting, and comparison with literature information was estimated at 2 hours – **SI Figure 3.1 - 1** and **SI Figure 3.2 - 2**.

All replicate injection files (n=5) of NIST SRM 1950 were then queried with MassQL, utilizing the anticipated $^{34}S$ isotope pattern and the presence of *m/z* 79.9563 ($SO_3^-$) and *m/z* 98.9552 ($FSO_3^-$) product ions. A MassQL query utilizing only the $^{34}S$ isotope pattern yielded too many hits to be of use. Using our workflow described here, the query took 53 minutes to perform (link above). The resulting query table was interpreted, and the MS/MS spectra were quickly reviewed using the Universal Spectrum Resolver Interface[57]. The MS/MS spectra for putative PFOS and PFHxS were recapitulated using the MassQL query (**SI Figure 3.1 - 3** and **SI Figure 3.1 - 4**); the relative abundance of product ions varied as the scan for PFOS was selected from a subsequent file (ID_n_03) opposed to the manual example. The presence of *m/z* 79.9563 ($SO_3^-$) and *m/z* 98.9552 ($FSO_3^-$) supports the putative annotation containing at least one fluorine atom and a sulfate group. Additional hits were observed in the query table; however, exhaustive interpretation was not performed due to time. In summary, the MassQL provided an easy and efficient method to query multiple data files and provide a tabular result for exploration of the data.

**SI Table 3.1 - 1.** Putatively annotated perfluorinated sulfonates in NIST SRM 1950

| **Name** | **Molecular Formula** | **Theoretical Monoisotopic Mass** | **Theoretical *m/z* $[M-H]^-$** |
|---|---|---|---|
| Perfluorohexanesulfonic acid (PFHxS) | $C_6HF_{13}O_3S$ | 399.9439 | 398.9366 |

| | | | |
|---|---|---|---|
| Perfluorooctanesulfonic acid (PFOS) | $C_8HF_{17}O_3S$ | 499.9375 | 498.9302 |

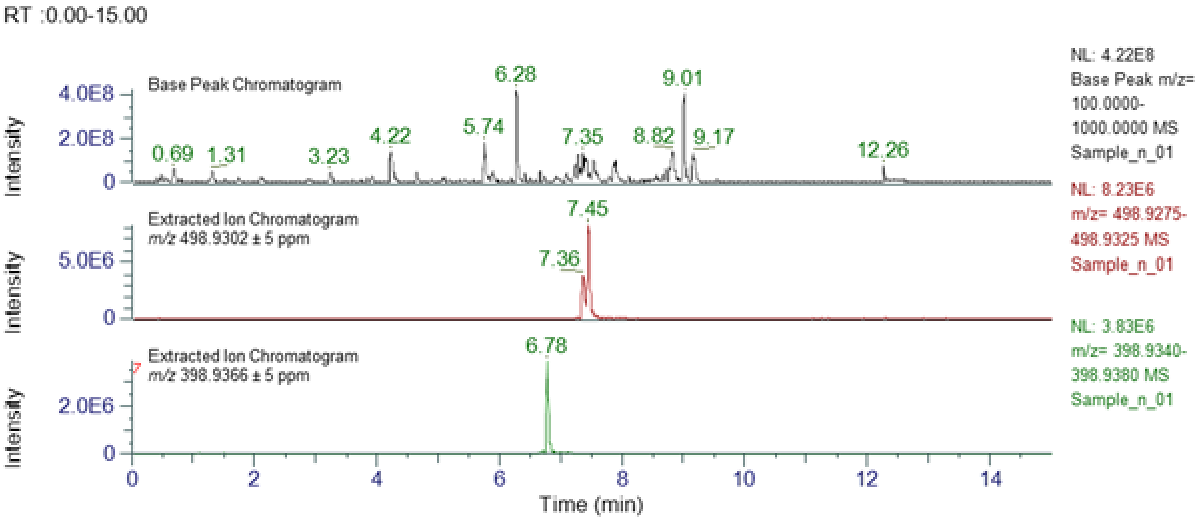


**SI Figure 3.1 - 1.** Base peak chromatogram of NIST SRM 1950 and extracted ion chromatograms for putatively annotated PFOS (*m/z* 498.9302 ± 5 ppm) and PFHxS (*m/z* 398.9366 ± 5 ppm).

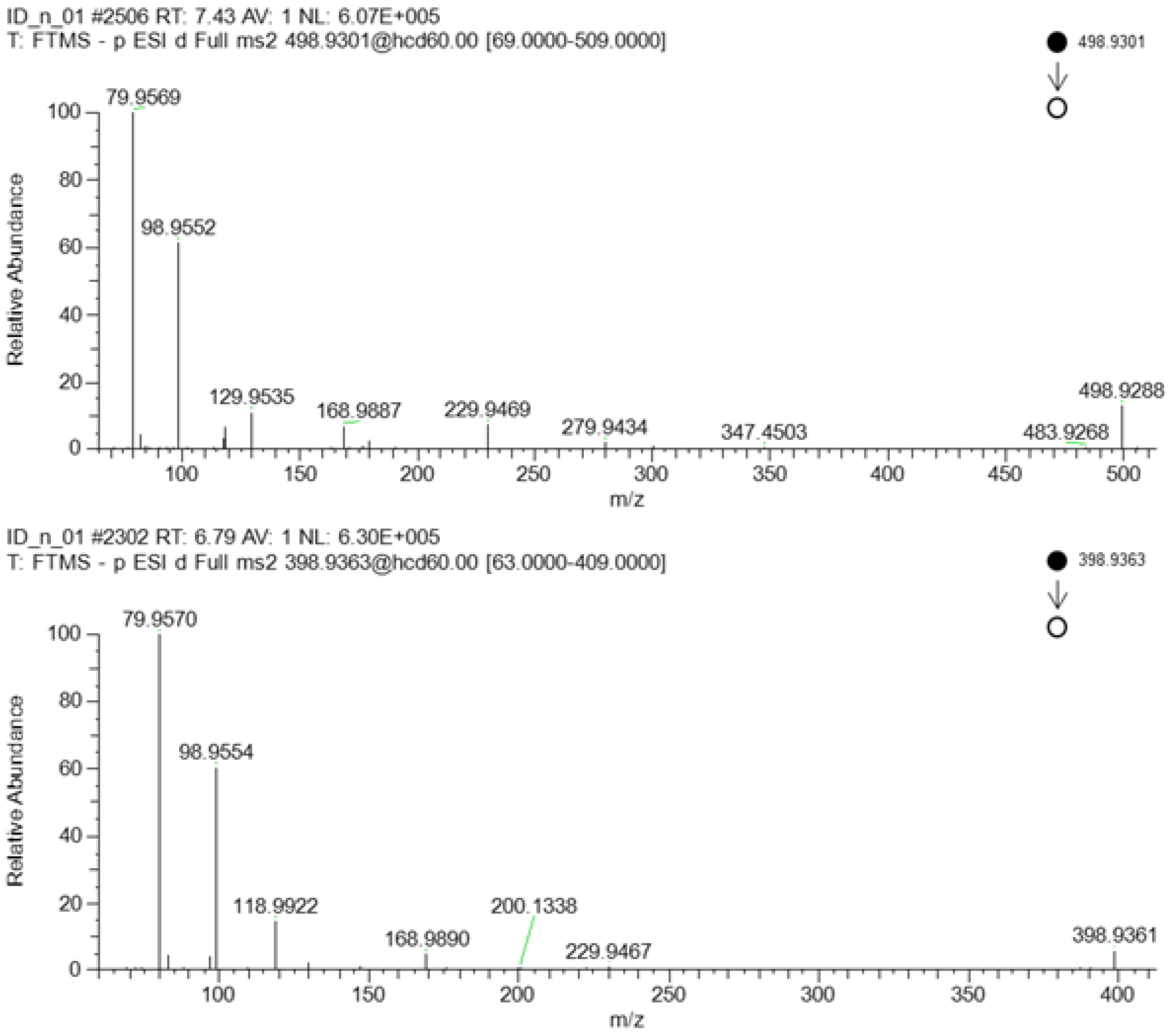


**SI Figure 3.1 - 2.** MS/MS spectra for putatively annotated Perfluorooctanesulfonic acid (*m/z* 498.9302 ± 5 ppm) and Perfluorohexanesulfonic acid (*m/z* 398.9366 ± 5 ppm) from NIST SRM 1950.

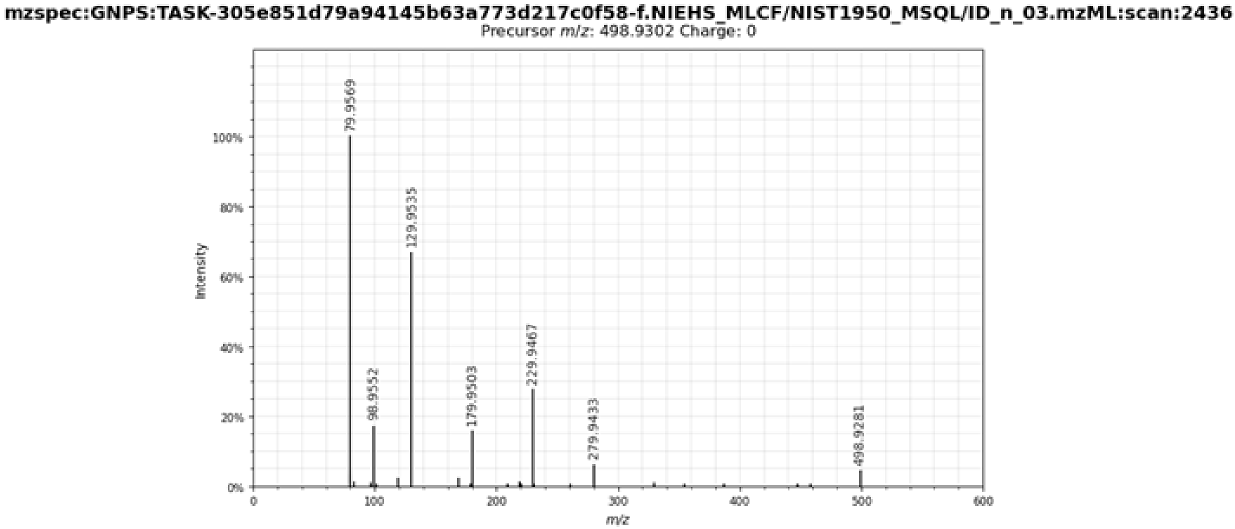


**SI Figure 3.1 - 3.** MS/MS spectra for putatively annotated Perfluorooctanesulfonic acid from NIST SRM 1950 obtained using MassQL (mzspec:GNPS:TASK-305e851d79a94145b63a773d217c0f58-f.NIEHS_MLCF/NIST1950_MSQL/ID_n_03.mzML:scan:2436)

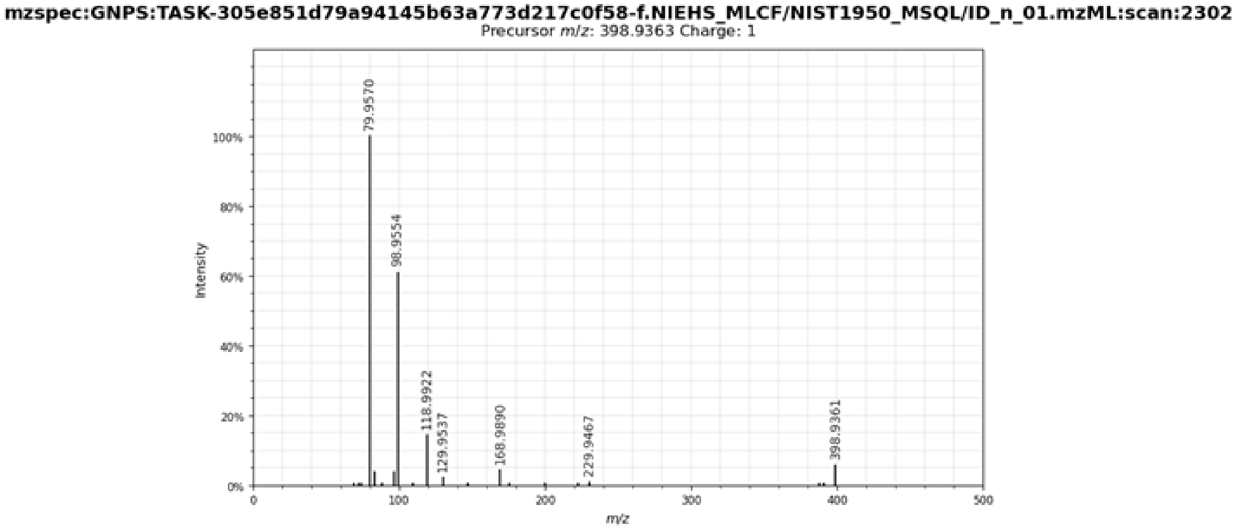


**SI Figure 3.1 - 4.** MS/MS spectra for putatively annotated Perfluorohexanesulfonic acid from NIST SRM 1950 obtained using MassQL (mzspec:GNPS:TASK-305e851d79a94145b63a773d217c0f58-f.NIEHS_MLCF/NIST1950_MSQL/ID_n_01.mzML:scan:2302)

##### 3.2.1 Exploring Drug Metabolism in Human Plasma via MassQL

**Author/s: Alan K. Jarmusch and Kirsten E. Overdahl**

| **MassQL Query**<br>`QUERY scaninfo(MS2DATA) WHERE`<br>`MS2PROD=167.0857:TOLERANCEPPM=5:INTENSITYPERCENT=10` |
|---|
| **MassQL Translation**<br>`Returning the scan information on MS2.`<br>`The following conditions are applied to find scans in the mass spec data.`<br>`Finding MS2 peak at m/z 167.0857 a 5.0 PPM tolerance and a minimum percent intensity relative to base peak of 10.0%.` |
| **MassQL Query Link**<br>https://proteomics2.ucsd.edu/ProteoSAFe/status.jsp?task=afceda948f2c49b1ac8973193c0e9d95 |
| **Data Availability**<br>MSV000085944 |

The metabolism of drugs is complex and variable in humans. Much work has gone into developing computational tools to predict drug transformations and into developing methods by which to highlight drugs and drug metabolites in mass spectrometry data. While untargeted mass spectrometry data can be explored via many methods and tools, MassQL provides the means to selectively, yet flexibility, search for drugs and drug metabolites in complex data.

We queried a single subject sample from publicly-available human drug metabolism data (MSV000085944). In the study, human volunteers were given diphenhydramine (single oral dose, 50 mg) and were subject to multiple blood draws over a 24 hr period. The sample of subject 07 (taken after 2hrs) was examined for diphenhydramine metabolites. Manual interpretation of the data revealed an *m/z* matching the theoretical *m/z* for diphenhydramine ($[M+H]^+$) at 3.41 min. Manual inspection of the associated MS/MS product ion scan indicated characteristic product ions of *m/z* 167.0857 (diphenyl cation). Manual exploration (calculation of mass error in the monoisotopic *m/z* value, product ion scan evaluation, and plotting) was repeated for the known diphenhydramine metabolites including desmethyldiphenhydramine, diphenhydramine N-oxide, and diphenhydramine N-glucuronide. Time spent on manual interpretation was estimated at 1 hour.

The following query was performed via MassQL which reported MS2 scans which contained the characteristic diphenyl cation of diphenhydramine and its metabolites. From hundreds of MS/MS product ion scans in the file, 9 scans meet the query criteria (**SI Table 3.2.1 - 1**). The query was completed in 6 minutes and 13 seconds. The product ion scan of diphenhydramine (scan 1316) was included in the results, and matched that obtained via manual interpretation. Further, additional metabolites were observed including desmethyldiphenhydramine (scan 1293), diphenhydramine N-oxide (scan 1369), and diphenhydramine N-glucuronide (1237).

**SI Table 3.2.1 - 1.** MassQL query results for finding MS2 spectra containing the diphenyl cation product ion

| **Scan** | **Retention Time (min)** | **Measured Precursor *m/z*** | **Abundance** | **File** |
|---|---|---|---|---|
| 1237 | 3.2 | 432.2023 | 165552.9 | MSV000085944/peak/blood/bld_plt1_07_120_1.mzXML |
| 1288 | 3.34 | 167.0857 | 788076.9 | MSV000085944/peak/blood/bld_plt1_07_120_1.mzXML |
| 1293 | 3.36 | 242.154 | 72227.56 | MSV000085944/peak/blood/bld_plt1_07_120_1.mzXML |
| 1316 | 3.42 | 256.1697 | 779002 | MSV000085944/peak/blood/bld_plt1_07_120_1.mzXML |
| 1335 | 3.47 | 167.1544 | 179702.8 | MSV000085944/peak/blood/bld_plt1_07_120_1.mzXML |
| 1369 | 3.56 | 272.1646 | 430642.7 | MSV000085944/peak/blood/bld_plt1_07_120_1.mzXML |
| 1442 | 3.76 | 167.0857 | 256290.5 | MSV000085944/peak/blood/bld_plt1_07_120_1.mzXML |
| 1640 | 4.3 | 167.0856 | 3285581 | MSV000085944/peak/blood/bld_plt1_07_120_1.mzXML |
| 1670 | 4.38 | 270.1491 | 28608.77 | MSV000085944/peak/blood/bld_plt1_07_120_1.mzXML |

##### 3.2.2 Exploring Drug Metabolism in Human Plasma via MassQL: Product and Neutral Loss

**Author/s: Alan K. Jarmusch and Kirsten E. Overdahl**

**MassQL Query**

```
QUERY scaninfo(MS2DATA) WHERE
MS2PROD=167.0857:TOLERANCEPPM=5:INTENSITYPERCENT=10 AND
MS2NL=176.0321:TOLERANCEPPM=5
```

**MassQL Translation**

```
Returning the scan information on MS2.
The following conditions are applied to find scans in the mass spec data.
Finding MS2 peak at m/z 167.0857 a 5.0 PPM tolerance and a minimum percent
intensity relative to base peak of 10.0%.
Finding MS2 neutral loss peak at m/z 176.0321 a 5.0 PPM tolerance.
```

**MassQL Query Link**

https://proteomics2.ucsd.edu/ProteoSAFe/status.jsp?task=f0fcd29d096d4d84be84471b5d984204

**Data Availability**

MSV000085944

The following more specific query was performed to highlight any ions which contained the characteristic diphenyl cation (167.0857) and the neutral loss of 176.0321 associated with glucuronidation, a phase II metabolic process. The query was completed in 6 minutes and 5 seconds. One product ion scan (1237), **SI Figure 3.2.2 - 1**, was observed to meet the criteria of the query, containing the neutral loss between *m/z* 432.2023 and 256.1696 ([diphenhydramine+H]$^+$) as well as the diphenyl cation (*m/z* 167.0850).

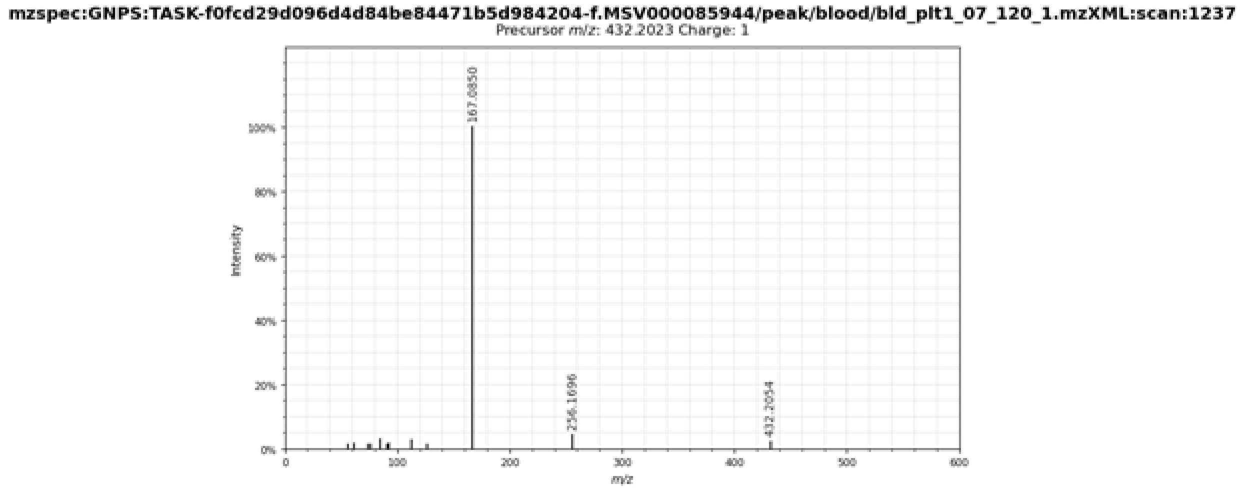


**SI Figure 3.2.2 - 1.** MS/MS spectrum (scan 1237) that resulted from the MassQL query.

##### 3.3.1 Discovering Iron Binding Molecules from Fungi: $^{54}$Fe peak

**Author/s: Allegra Aron**

| **MassQL Query** |
| --- |
| `QUERY scaninfo(MS1DATA) WHERE MS1MZ=X-2:INTENSITYMATCH=Y*0.063:INTENSITYMATCHPERCENT=25 AND MS1MZ=X:INTENSITYMATCH=Y:INTENSITYMATCHREFERENCE:INTENSITYPERCENT=5 AND MS2PREC=X FILTER MS1MZ=X` |
| **MassQL Translation** |
| Returning the scan information on MS1.<br>The following conditions are applied to find scans in the mass spec data.<br>Finding MS1 peak at m/z X-2.0 an expected relative intensity to reference peak of Y*0.063and accepting variability of 25.0% in relative intensity.<br>Finding MS1 peak at m/z X an expected relative intensity to reference peak of Yand this peak is used as the intensity reference for other peaks in the spectrumand a minimum percent intensity relative to base peak of 5.0%.<br>Finding MS2 spectra with a precursor m/z X.<br>Finding MS1 peak at m/z X. |
| **MassQL Query Link** |
| https://proteomics2.ucsd.edu/ProteoSAFe/status.jsp?task=8ac5da634dc74c16a2a8d639d24244d1 |
| **Data Availability** |
| MSV000084030 |

Systematic methods for the discovery of metal-small molecule complexes from biological samples are limited, even as metals play a number of essential roles in biology. Recently, we described a two-step native electrospray ionization mass spectrometry method, in which post-column pH adjustment and metal-infusion are combined with ion identity molecular networking to facilitate identification of metal-binding compounds in complex samples based on defined mass (*m/z*) offsets of ion species with the same chromatographic profiles[58]. Combining the native metabolomics experimental workflow with the requirements for a metal-specific isotope pattern in the MS1[59] can add an additional layer of confidence to metal-binding molecules identified using the native metabolomics approach. MassQL can be used as an alternate analysis strategy to the ion identity molecular networking (IIMN) workflow utilized in native metabolomics. Additionally, we envisioned carrying out a repository search using the search query optimized below in order to identify all potential iron-binding molecules in the GNPS-MassIVE repository.

In order to test this approach, the MassQL was assessed on extracts from the wine fungus *Eutypa lata* that were treated with a post-liquid chromatography iron addition, as described in Aron *et al*.[58] The initial query for iron-binding molecules contained a requirement for iron isotope pattern in MS1. This first (and simplest) query searches for a peak at *x* with intensity equal to 1 and a peak at *x-2* with intensity of 0.063; this is based on the stable isotope ratio of $^{54}$Fe to $^{56}$Fe. The simplest version of this search yielded multiple orders of magnitude more hits than the native metabolomics approach, and manual inspection of hits suggested that many were false positives.

##### 3.3.2 Discovering Iron Binding Molecules from Fungi: $^{54}$Fe peak, $^{13}$C peak, and apo peak

**Author/s: Allegra Aron**

| **MassQL Query** |
|---|
| QUERY scaninfo(MS1DATA) WHERE MS1MZ=X-1.993:INTENSITYMATCH=Y*0.063:INTENSITYMATCHPERCENT=25:TOLERANCEPPM=10 AND MS1MZ=X:INTENSITYMATCH=Y:INTENSITYMATCHREFERENCE:INTENSITYPERCENT=5 AND MS1MZ=X+1:INTENSITYMATCH=Y*0.5:INTENSITYMATCHPERCENT=60 AND MS1MZ=X-52.91:TOLERANCEPPM=10 AND MS2PREC=X FILTER MS1MZ=X |
| **MassQL Translation**<br>Returning the scan information on MS1.<br>The following conditions are applied to find scans in the mass spec data.<br>Finding MS1 peak at m/z X-1.993 an expected relative intensity to reference peak of Y*0.063and accepting variability of 25.0% in relative intensity and a 10.0 PPM tolerance.<br>Finding MS1 peak at m/z X an expected relative intensity to reference peak of Yand this peak is used as the intensity reference for other peaks in the spectrum and a minimum percent intensity relative to base peak of 5.0%.<br>Finding MS1 peak at m/z X+1.0 an expected relative intensity to reference peak of Y*0.5and accepting variability of 60.0% in relative intensity.<br>Finding MS1 peak at m/z X-52.91 a 10.0 PPM tolerance.<br>Finding MS2 spectra with a precursor m/z X.<br>Finding MS1 peak at m/z X. |
| **MassQL Query Link**<br>https://proteomics2.ucsd.edu/ProteoSAFe/status.jsp?task=b217f5a481f24fd7852e695a54d8a797 |
| **Additional Data Analysis**<br>n/a |
| **Data Availability**<br>MSV000084030 |

In order to remove the false positives described in Use Case 3.3.1, the next query included the requirement for a $^{13}$C isotope peak at *x+1* and an apo (unbound) peak at *x-52.91*. This apo peak corresponds to an *m/z* delta of [M+Fe-2H]+ in MS1; the presence of this unbound peak in the same scan as the bound compound adds confidence in the hit, as both peaks are commonly observed in the same scan (especially when a post-LC iron infusion is utilized). Finally, *m/z* tolerances were set to 10 ppm to take advantage of the Q-Exactive Orbitrap used for these experiments. This more stringent query yielded approximately ninety unique compounds with *m/z* >400. Six of these compounds were also found using the native metabolomics workflow, including coprogen B and hydroxymethyl coprogen B. Interestingly, two compounds that were identified using native metabolomics were not identified using the MassQL workflow - iron-bound *m/z* 838.3049 and *m/z* 910.3612. To understand why these compounds were identified using native metabolomics but not MassQL, we investigated the raw data using the MassQL visualizer (https://msql.ucsd.edu/) and found that the *x-2* isotope peak in both compounds fell outside of the specified *m/z* tolerance and the intensity percent tolerance (**SI Figure 3.3.2 - 2**). Given this, these two putative iron-binding compounds from the native metabolomics workflow are less reliable hits than the other six compounds that were identified using both native metabolomics and MassQL, though the low *m/z* accuracy could be due to the fact that this isotope peak is low abundance and so these hits should not be completely disregarded. The other six compounds exhibit both apo and bound *m/z* in the same scan and also satisfy the predicted iron isotope pattern both in intensity and in *m/z* tolerance.

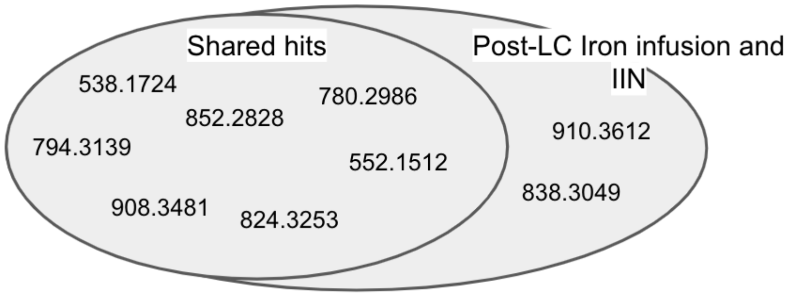


**SI Figure 3.3.2 - 1.** Comparison of *m/z* found using MassQL versus the native metabolomics (post-LC iron infusion and ion identity networking) workflow.

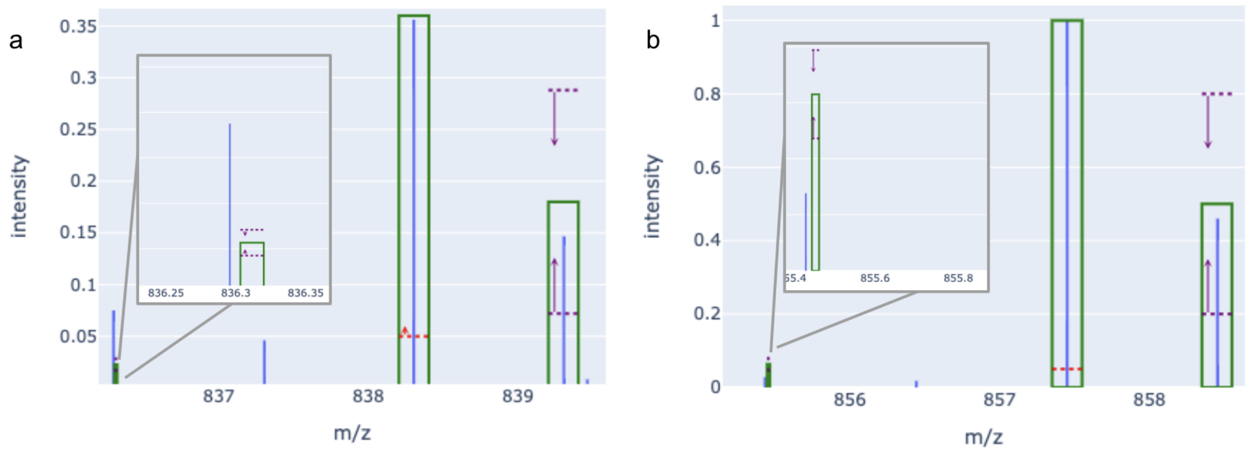


**SI Figure 3.3.2 - 2.** Raw data visualized using the MassQL visualizer (https://msql.ucsd.edu/) for (a) *m/z* 838.3049 and (b) *m/z* 910.3612. The inset zoom highlights the *x-2* isotope peak in both compounds, which falls outside of the specified *m/z* tolerance and the specified intensity percent tolerance for both compounds.

#### 3.4 Discovering Putative Iron Binding Molecules in Repository Search

**Author/s: Allegra Aron**

| **MassQL Query** |
| --- |
| `QUERY scaninfo(MS1DATA) WHERE MS1MZ=X-1.993:INTENSITYMATCH=Y*0.063:INTENSITYMATCHPERCENT=25:TOLERANCEPPM=10 AND MS1MZ=X:INTENSITYMATCH=Y:INTENSITYMATCHREFERENCE:INTENSITYPERCENT=5 AND MS1MZ=X+1:INTENSITYMATCH=Y*0.5:INTENSITYMATCHPERCENT=60 AND MS1MZ=X-52.91:TOLERANCEPPM=10 AND MS2PREC=X-52.91` |
| **MassQL Translation** |
| Returning the scan information on MS1.<br>The following conditions are applied to find scans in the mass spec data.<br>Finding MS1 peak at m/z X-1.993 an expected relative intensity to reference peak of Y*0.063and accepting variability of 25.0% in relative intensity and a 10.0 PPM tolerance.<br>Finding MS1 peak at m/z X an expected relative intensity to reference peak of Yand this peak is used as the intensity reference for other peaks in the spectrum and a minimum percent intensity relative to base peak of 5.0%.<br>Finding MS1 peak at m/z X+1.0 an expected relative intensity to reference peak of Y*0.5and accepting variability of 60.0% in relative intensity.<br>Finding MS1 peak at m/z X-52.91 a 10.0 PPM tolerance.<br>Finding MS2 spectra with a precursor m/z X-52.91. |
| **MassQL Query Link** |
| https://proteomics3.ucsd.edu/ProteoSAFe/status.jsp?task=05974eace08047108091afa5f07899d6 |
| **Data Availability** |
| Full GNPS-MassIVE repository |

Given the fact that MassQL can identify iron-bound compounds in *Eutypa lata* extracts treated with post-LC iron infusion, we were eager to test the MassQL language to find putative iron-binding compounds in the MassIVE-GNPS repository. A repository search utilizing MassQL would rely on the fact that some iron-binding compounds will still remain bound to iron with both *x* and *x-2* at detectable levels. This strategy is complementary to the native metabolomics strategy - while it does not require any experimental set-up or pre-processing using feature finding and can be applied to a repository search, the requirement for iron-binding means that many weak-binding or low abundance compounds will not be observed. Even with these caveats, we still found 946,743 spectra matching the query above.

##### 3.5.1 Polybrominated analogs and putative biosynthetic precursors of the "eagle killer toxin", aetokthonotoxin

**Author/s: Raphael Reher, Timo Niedermeyer**

Recently, the pentabrominated biindole alkaloid aetokthonotoxin (AETX) was identified as cyanobacterial neurotoxin that acts as the causal agent of fatal vacuolar myelinopathy, responsible for the mass mortality of bald eagles and other animals in the southeastern US[60]. The study shows the complex indirect and direct anthropogenic effects on the natural world, linking invasive plants as substrate for the growth of a previously unidentified cyanobacterium, exposure to high bromide concentrations, and cyanobacterial production of a neurotoxin leading to fatal neuropathy in animals that prey on the plants and also bioaccumulates to kill predators such as bald eagles. Here, we used iterative MassQL queries to discover new di-, tri-, and tetra-, and pentabrominated analogs of AETX as well as putative biosynthetic precursors from extracts derived from the cyanobacterium *Aetokthonos hydrillicola*, sampled from lakes where vacuolar myelinopathy outbreaks were reported as well as from laboratory cultures.

Indeed, applying iterative MassQL queries for the characteristic MS1 isotopic patterns of various bromination levels, we succeeded in the detection of numerous putatively new di-, tri-, tetra-, and pentabrominated AETX analogs (**SI Figure 3.5.1 - 1**). All structures are proposals based on MS1/MS2 data only and have to be confirmed by isolation and rigorous subsequent structure elucidation.

The single queries for the various bromination levels could also have been piped into one big query, but for the ease of downstream data analysis, we recommend having the single queries as specific as possible to keep the numbers of detected features and false positives low. This recommendation is especially true for repository scale queries. Furthermore, in the case of polybrominated natural products, there are other effects that artificially increase the number of detected features, e.g. for pentabrominated natural products there are already six *m/z* features (X-4, X-2, X, X+2, X+4, X+6) for a single natural product on the MS1 level. These six features then get multiplied by each adduct (e.g. $[M+H]^+$, $[M-H]^-$ , $[M+Na]^+$ etc.) and again multiplied by repeated detection at several retention time points, if not excluded during the data acquisition.

As with every new tool, MassQL also bears the danger of over- and/or misinterpretation. To detect the very low abundant tribrominated AETX analogs, we lowered the minimum percent intensity relative to the base peak to 5.0%. In that way, we were able to detect the putative AETX-$Br_3$ analog *m/z* 489.8199. However, the lower the intensity, the harder it gets to distinguish the detected isotopic pattern from noise. On a related note, mono- and tribrominated isotopic patterns are a subset of pentabrominated isotopic patterns. After manual inspection of the feature *m/z* 661.6385, it seems more likely to us that this feature is actually a hydroxylated pentabrominated analog of AETX, rather than atribrominated AETX precursor. However, *m/z* 661.6385 was not detected with the query for pentabrominated natural products, because the X-4 peak was not assigned to the corresponding isotopic peaks for X-2, X, X+2, X+4, and the X+6 is missing completely due to the general very low abundance of the compound (**SI Figure 3.5.1 - 2**). For future optimization, we aim to use the MassQL "EXCLUDE" operator to design more selective queries that can distinguish between mono-, tri-, and penta as well as di-, and tetrabrominated compounds from each other.

Concluding, MassQL queries can greatly assist researchers to answer specific mass spec-related questions, while still requiring careful manual inspection of the results.

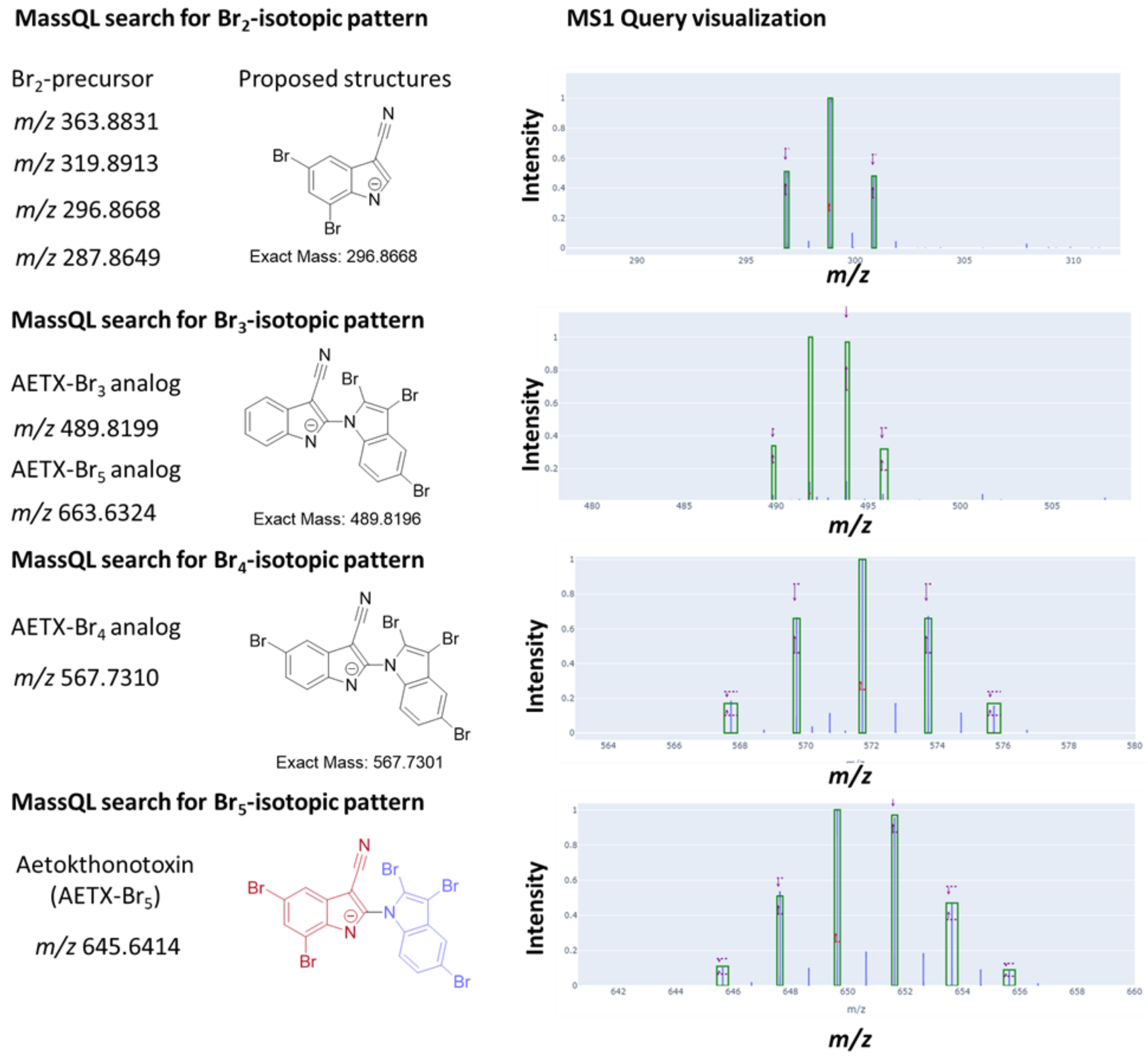


**SI Figure 3.5.1 - 1**: MassQL queries for di-, tri-, tetra-, and pentabrominated natural products were used to detect several putatively new analogs and precursors of the cyanobacterial “eagle killer toxin”, aetokthonotoxin.

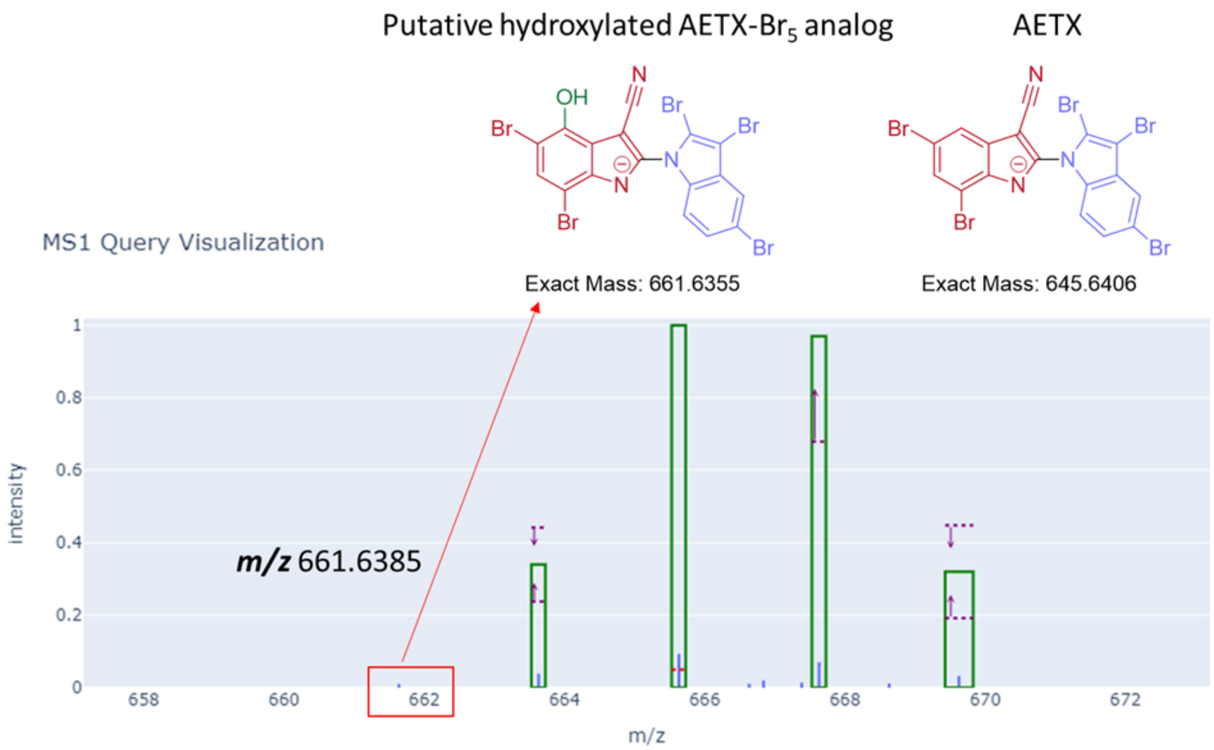


**SI Figure 3.5.1 - 2**: Low abundant features might lead to wrong assignments. A probable pentabrominated AETX analog m/z 661.6385 was detected with the query for tribrominated compounds, because of the low abundancy or missing X-4 and X+6 peaks, respectively. The position for the hydroxylation is speculative and serves visualization purposes only.

##### 3.5.1.1 Polybrominated analogs and precursor of the “eagle killer” aetokthonotoxin - Dibrominated precursor/analogs

**MassQL Query**

```
QUERY scaninfo(MS2DATA) WHERE
MS1MZ=X:TOLERANCEMZ=0.1:INTENSITYPERCENT=25:INTENSITYMATCH=Y:INTENSITYMATCH
REFERENCE
AND
MS1MZ=X+2:TOLERANCEMZ=0.1:INTENSITYMATCH=Y*0.48:INTENSITYMATCHPERCENT=30
AND MS1MZ=X-
2:TOLERANCEMZ=0.1:INTENSITYMATCH=Y*0.51:INTENSITYMATCHPERCENT=30
AND
MS2PREC=X:TOLERANCEMZ=4 AND X=range(min=200, max=700)
```

**MassQL Translation**

```
Returning the scan information on MS2.
The following conditions are applied to find scans in the mass spec data.
Finding MS1 peak at m/z X with a 0.1 m/z tolerance and a minimum percent
intensity relative to base peak of 25.0% and an expected relative intensity to
reference peak of Y and this peak is used as the intensity reference for other
peaks in the spectrum.
Finding MS1 peak at m/z X+2.0 with a 0.1 m/z tolerance and an expected relative
intensity to reference peak of Y*0.48 and accepting variability of 30.0% in
relative intensity.
Finding MS1 peak at m/z X-2.0 with a 0.1 m/z tolerance and an expected relative
intensity to reference peak of Y*0.51 and accepting variability of 30.0% in
relative intensity.
Finding MS2 spectra with a precursor m/z X with a 4.0 m/z tolerance.
Enabling variable X with range (200.0, 700.0).
```

**MassQL Query Link**

https://gnps.ucsd.edu/ProteoSAFe/status.jsp?task=9dc6314862ff436a96f0d835507c1bf6

**Data Availability**

MSV000088461

##### 3.5.1.2 Polybrominated analogs and precursor of the “eagle killer” aetokthonotoxin - Tribrominated precursor/analogs

**MassQL Query**

```
QUERY scaninfo(MS2DATA) WHERE
MS1MZ=X:TOLERANCEMZ=0.1:INTENSITYPERCENT=5:INTENSITYMATCH=Y:INTENSITYMATCHR
EFERENCE

AND
MS1MZ=X+2:TOLERANCEMZ=0.1:INTENSITYMATCH=Y*0.97:INTENSITYMATCHPERCENT=30

AND MS1MZ=X-
2:TOLERANCEMZ=0.1:INTENSITYMATCH=Y*0.34:INTENSITYMATCHPERCENT=30
```

```
AND
MS1MZ=X+4:TOLERANCEMZ=0.2:INTENSITYMATCH=Y*0.32:INTENSITYMATCHPERCENT=40
AND

MS2PREC=X:TOLERANCEMZ=2 AND X=range(min=300, max=900)
```

**MassQL Translation**

```
Returning the scan information on MS2.
The following conditions are applied to find scans in the mass spec data.
Finding MS1 peak at m/z X with a 0.1 m/z tolerance and a minimum percent
intensity relative to base peak of 5.0% and an expected relative intensity to
reference peak of Y and this peak is used as the intensity reference for other
peaks in the spectrum.
Finding MS1 peak at m/z X+2.0 with a 0.1 m/z tolerance and an expected relative
intensity to reference peak of Y*0.97 and accepting variability of 30.0% in
relative intensity.
Finding MS1 peak at m/z X-2.0 with a 0.1 m/z tolerance and an expected relative
intensity to reference peak of Y*0.34 and accepting variability of 30.0% in
relative intensity.
Finding MS1 peak at m/z X+4.0 with a 0.2 m/z tolerance and an expected relative
intensity to reference peak of Y*0.32 and accepting variability of 40.0% in
relative intensity.
Finding MS2 spectra with a precursor m/z X with a 2.0 m/z tolerance.
Enabling variable X with range (300.0, 900.0).
```

**MassQL Query Link**

https://gnps.ucsd.edu/ProteoSAFe/status.jsp?task=bf46e7d102e241fc925af3c05debdb93

**Data Availability**

MSV000088461

###### 3.5.1.3 Polybrominated analogs and precursor of the “eagle killer” aetokthonotoxin - Tetrabrominated precursor

**MassQL Query**

```
QUERY scaninfo(MS2DATA) WHERE
MS1MZ=X:TOLERANCEMZ=0.1:INTENSITYPERCENT=25:INTENSITYMATCH=Y:INTENSITYMATCH
REFERENCE

AND
MS1MZ=X+2:TOLERANCEMZ=0.1:INTENSITYMATCH=Y*0.66:INTENSITYMATCHPERCENT=30

AND MS1MZ=X-
2:TOLERANCEMZ=0.1:INTENSITYMATCH=Y*0.66:INTENSITYMATCHPERCENT=30

AND
MS1MZ=X+4:TOLERANCEMZ=0.2:INTENSITYMATCH=Y*0.17:INTENSITYMATCHPERCENT=40

AND MS1MZ=X-
4:TOLERANCEMZ=0.2:INTENSITYMATCH=Y*0.17:INTENSITYMATCHPERCENT=40

AND

MS2PREC=X:TOLERANCEMZ=4 AND X=range(min=400, max=1200)
```

**MassQL Translation**

Returning the scan information on MS2.
The following conditions are applied to find scans in the mass spec data.
Finding MS1 peak at m/z X with a 0.1 m/z tolerance and a minimum percent intensity relative to base peak of 25.0% and an expected relative intensity to reference peak of Y and this peak is used as the intensity reference for other peaks in the spectrum.
Finding MS1 peak at m/z X+2.0 with a 0.1 m/z tolerance and an expected relative intensity to reference peak of Y*0.66 and accepting variability of 30.0% in relative intensity.
Finding MS1 peak at m/z X-2.0 with a 0.1 m/z tolerance and an expected relative intensity to reference peak of Y*0.66 and accepting variability of 30.0% in relative intensity.
Finding MS1 peak at m/z X+4.0 with a 0.2 m/z tolerance and an expected relative intensity to reference peak of Y*0.17 and accepting variability of 40.0% in relative intensity.
Finding MS1 peak at m/z X-4.0 with a 0.2 m/z tolerance and an expected relative intensity to reference peak of Y*0.17 and accepting variability of 40.0% in relative intensity.
Finding MS2 spectra with a precursor m/z X with a 4.0 m/z tolerance.
Enabling variable X with range (400.0, 1200.0).

**MassQL Query Link**

https://gnps.ucsd.edu/ProteoSAFe/status.jsp?task=a28f790ccf5f4d5e8975b6985d5e0273

**Data Availability**

MSV000088461

##### 3.5.1.4 Polybrominated analogs and precursor of the "eagle killer" aetokthonotoxin - pentabrominated precursor/analogs

**MassQL Query**

```
QUERY scaninfo(MS2DATA) WHERE
MS1MZ=X:TOLERANCEMZ=0.1:INTENSITYPERCENT=25:INTENSITYMATCH=Y:INTENSITYMATCH
REFERENCE AND

MS1MZ=X+2:TOLERANCEMZ=0.1:INTENSITYMATCH=Y*0.97:INTENSITYMATCHPERCENT=10
AND

MS1MZ=X-2:TOLERANCEMZ=0.1:INTENSITYMATCH=Y*0.51:INTENSITYMATCHPERCENT=20
AND
MS1MZ=X+4:TOLERANCEMZ=0.2:INTENSITYMATCH=Y*0.47:INTENSITYMATCHPERCENT=20
AND

MS1MZ=X-4:TOLERANCEMZ=0.2:INTENSITYMATCH=Y*0.11:INTENSITYMATCHPERCENT=40
AND

MS1MZ=X+6:TOLERANCEMZ=0.2:INTENSITYMATCH=Y*0.09:INTENSITYMATCHPERCENT=40
AND

MS2PREC=X:TOLERANCEMZ=2 AND X=range(min=400, max=1200)
```

**MassQL Translation**

Returning the scan information on MS2.
The following conditions are applied to find scans in the mass spec data.
Finding MS1 peak at m/z X with a 0.1 m/z tolerance and a minimum percent intensity relative to base peak of 25.0% and an expected relative intensity to reference peak of Y and this peak is used as the intensity reference for other peaks in the spectrum.
Finding MS1 peak at m/z X+2.0 with a 0.1 m/z tolerance and an expected relative intensity to reference peak of Y*0.97 and accepting variability of 10.0% in relative intensity.
Finding MS1 peak at m/z X-2.0 with a 0.1 m/z tolerance and an expected relative intensity to reference peak of Y*0.51 and accepting variability of 20.0% in relative intensity.
Finding MS1 peak at m/z X+4.0 with a 0.2 m/z tolerance and an expected relative intensity to reference peak of Y*0.47 and accepting variability of 20.0% in relative intensity.
Finding MS1 peak at m/z X-4.0 with a 0.2 m/z tolerance and an expected relative intensity to reference peak of Y*0.11 and accepting variability of 40.0% in relative intensity.
Finding MS1 peak at m/z X+6.0 with a 0.2 m/z tolerance and an expected relative intensity to reference peak of Y*0.09 and accepting variability of 40.0% in relative intensity.
Finding MS2 spectra with a precursor m/z X with a 2.0 m/z tolerance.
Enabling variable X with range (400.0, 1200.0).

**MassQL Query Link**

https://gnps.ucsd.edu/ProteoSAFe/status.jsp?task=9ec8134461bc4ffebde566e022b2e478

**Data Availability**

MSV000088461

##### 3.5.2 Repository Scale MassQL Search to find Pentabrominated Natural Products

**Author/s: Raphael Reher, Timo Niedermeyer**

After the successful implementation of MassQL queries to find aeokthonotoxin analogs and precursors of various bromination levels in three mzML files, we wondered whether we can detect pentabrominated molecules similar to AETX in all other datasets from GNPS that used the same mass analyzer (Q exactive). We therefore queried all QE-data against the characteristic MS1 pentabromine isotopic pattern as a query.

Analyzing the resulting 425 features, we detected an interesting pentabrominated feature with *m/z* 641 that matches the exact mass of aspidostomide F, a bromotryptophan-derived moiety linked to a pyrroloketopiperazine lactam, from a marine bryozoan (**SI Figure 3.5.2 - 1**). There is some structural similarity to aetokthonotoxin such as the dibrominated indole moiety that is coupled to another tribrominated *N*-heterocycle. However, the bromination pattern of both heterodimeric moieties is different for both natural products.

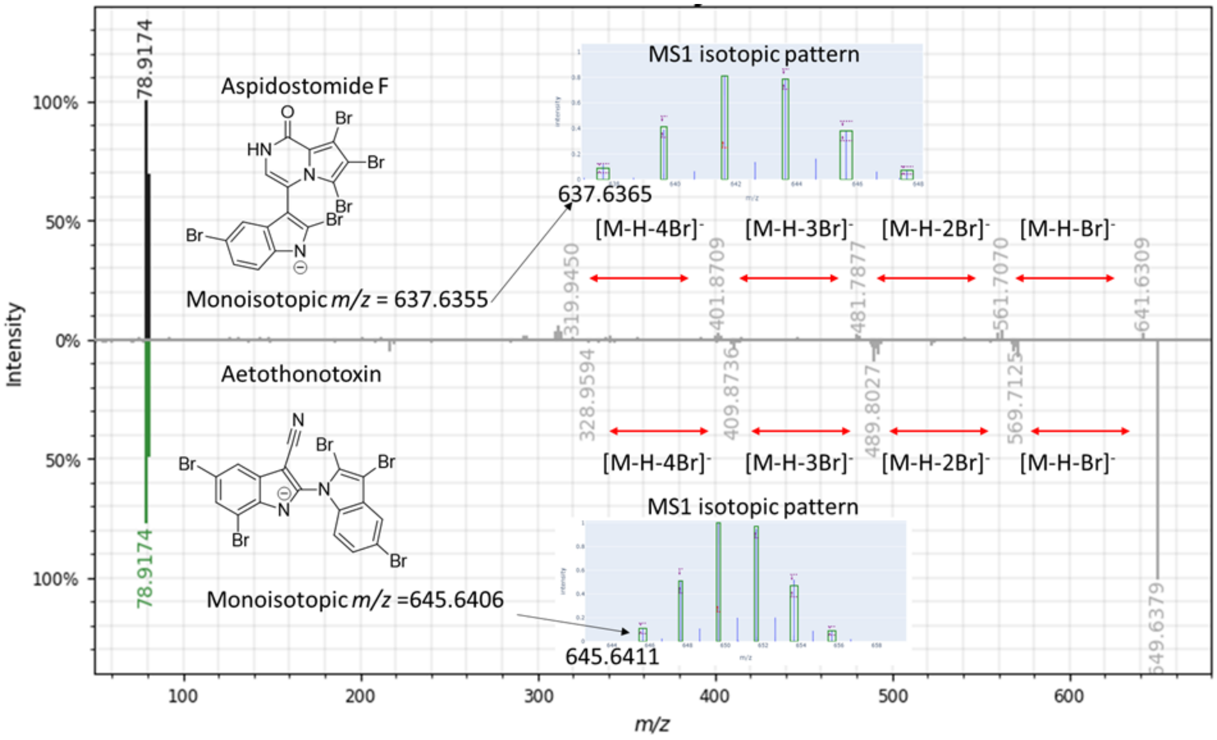


**SI Figure 3.5.2 - 1**: Comparison of MS2 spectra of the cyanobacterial aetokthonotoxin and the pentabrominated aspidostomide F found in a *Streptomyces* dataset.

Strikingly, even though originally reported from the marine bryozoan, in this search we found aspidostomide F[61] in a Canadian *Streptomyces* dataset (MSV000084595). We were curious whether we could find new tetrabrominated analogs, as we did for AETX. Indeed, we found an *m/z* feature corresponding to a compound containing one bromine less than aspidostomide F (characteristic tetrabrominated isotopic MS1 pattern, high cosine similarity (0.95) to aspidostomide F; **SI Figure 3.5.2 - 2**).

Concluding, MassQL is an exciting tool that allows the user to design their queries to answer their often very specific research questions. Here, we demonstrated the use of MassQL to mine small as well as repository scale datasets for polybrominated natural products of ecological/environmental relevance.

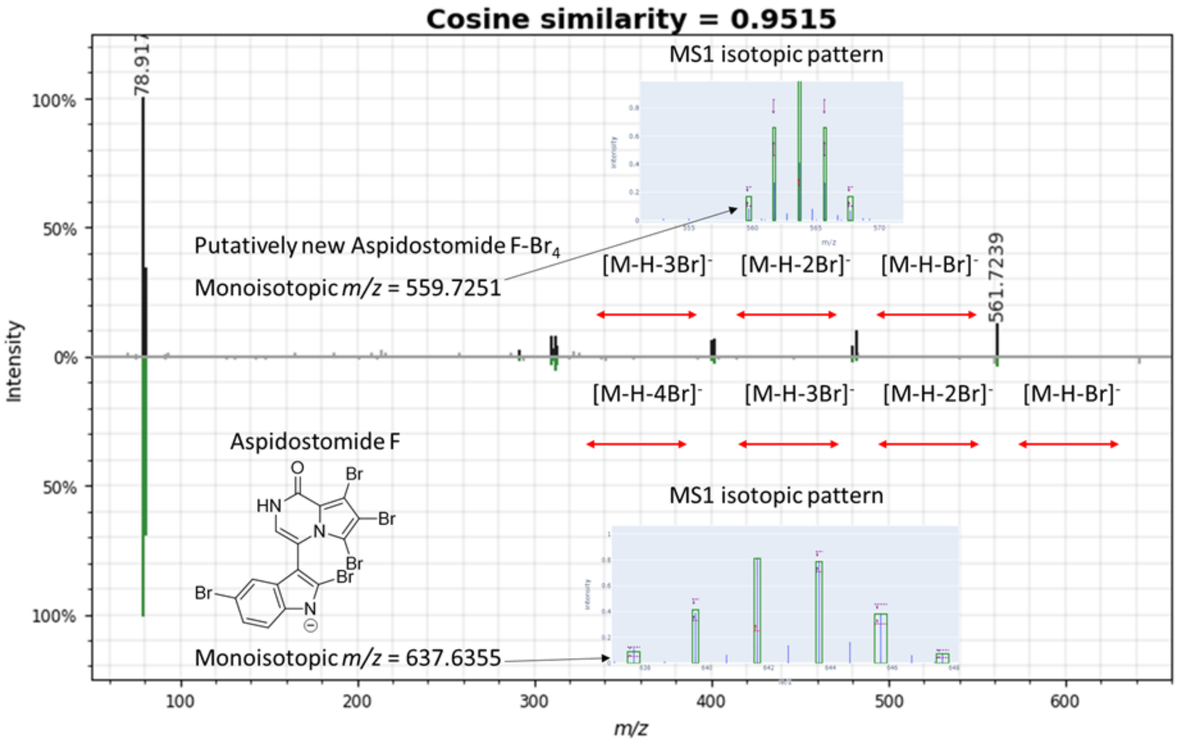


**SI Figure 3.5.2 - 2**: MS2 spectra of aspidobromide A and a putatively new tetrabrominated analog from Streptomyces (MSV000084595).

##### 3.5.2.1 Repository Scale MassQL search for pentabrominated Natural Products

**MassQL Query**

```
QUERY scaninfo(MS2DATA) WHERE

MS1MZ=X:TOLERANCEMZ=0.1:INTENSITYPERCENT=5:INTENSITYMATCH=Y:INTENSITYMATCHR
EFERENCE AND

MS1MZ=X+2:TOLERANCEMZ=0.1:INTENSITYMATCH=Y*0.97:INTENSITYMATCHPERCENT=10
AND

MS1MZ=X-2:TOLERANCEMZ=0.1:INTENSITYMATCH=Y*0.51:INTENSITYMATCHPERCENT=20
AND
MS1MZ=X+4:TOLERANCEMZ=0.2:INTENSITYMATCH=Y*0.47:INTENSITYMATCHPERCENT=20
AND

MS1MZ=X-4:TOLERANCEMZ=0.2:INTENSITYMATCH=Y*0.11:INTENSITYMATCHPERCENT=40
AND

MS1MZ=X+6:TOLERANCEMZ=0.2:INTENSITYMATCH=Y*0.09:INTENSITYMATCHPERCENT=40
AND

MS2PREC=X:TOLERANCEMZ=4 AND X=range(min=500, max=1500)
```

**MassQL Translation**

```
Returning the scan information on MS2.
The following conditions are applied to find scans in the mass spec data.
Finding MS1 peak at m/z X with a 0.1 m/z tolerance and a minimum percent
intensity relative to base peak of 5.0% and an expected relative intensity to
reference peak of Y and this peak is used as the intensity reference for other
peaks in the spectrum.
Finding MS1 peak at m/z X+2.0 with a 0.1 m/z tolerance and an expected relative
intensity to reference peak of Y*0.97 and accepting variability of 10.0% in
relative intensity.
Finding MS1 peak at m/z X-2.0 with a 0.1 m/z tolerance and an expected relative
intensity to reference peak of Y*0.51 and accepting variability of 20.0% in
relative intensity.
Finding MS1 peak at m/z X+4.0 with a 0.2 m/z tolerance and an expected relative
intensity to reference peak of Y*0.47 and accepting variability of 20.0% in
relative intensity.
Finding MS1 peak at m/z X-4.0 with a 0.2 m/z tolerance and an expected relative
intensity to reference peak of Y*0.11 and accepting variability of 40.0% in
relative intensity.
Finding MS1 peak at m/z X+6.0 with a 0.2 m/z tolerance and an expected relative
```

```
intensity to reference peak of Y*0.09 and accepting variability of 40.0% in
relative intensity.
Finding MS2 spectra with a precursor m/z X with a 4.0 m/z tolerance.
Enabling variable X with range (500.0, 1500.0).
```

| |
|---|
| **MassQL Query Link**<br>https://proteomics2.ucsd.edu/ProteoSAFe/status.jsp?task=a932337a3898426d9c3faf33d687ef5a |
| **Additional Data Analysis**<br>n/a |
| **Data Availability**<br>MSV000088461 |

###### 3.5.2.2 MassQL search for tetrabrominated analogs of aspidostomide F in *Streptomyces*

**MassQL Query**

```
QUERY scaninfo(MS2DATA) WHERE
MS1MZ=X:TOLERANCEMZ=0.1:INTENSITYPERCENT=5:INTENSITYMATCH=Y:INTENSITYMATCHR
EFERENCE

AND
MS1MZ=X+2:TOLERANCEMZ=0.1:INTENSITYMATCH=Y*0.66:INTENSITYMATCHPERCENT=30

AND MS1MZ=X-
2:TOLERANCEMZ=0.1:INTENSITYMATCH=Y*0.66:INTENSITYMATCHPERCENT=30

AND
MS1MZ=X+4:TOLERANCEMZ=0.2:INTENSITYMATCH=Y*0.17:INTENSITYMATCHPERCENT=40

AND MS1MZ=X-
4:TOLERANCEMZ=0.2:INTENSITYMATCH=Y*0.17:INTENSITYMATCHPERCENT=40

AND

MS2PREC=X:TOLERANCEMZ=4 AND X=range(min=400, max=900)
```

**MassQL Translation**

Returning the scan information on MS2.

The following conditions are applied to find scans in the mass spec data.

Finding MS1 peak at m/z X with a 0.1 m/z tolerance and a minimum percent intensity relative to base peak of 5.0% and an expected relative intensity to reference peak of Y and this peak is used as the intensity reference for other peaks in the spectrum.

Finding MS1 peak at m/z X+2.0 with a 0.1 m/z tolerance and an expected relative intensity to reference peak of Y*0.66 and accepting variability of 30.0% in relative intensity.

Finding MS1 peak at m/z X-2.0 with a 0.1 m/z tolerance and an expected relative intensity to reference peak of Y*0.66 and accepting variability of 30.0% in relative intensity.

Finding MS1 peak at m/z X+4.0 with a 0.2 m/z tolerance and an expected relative intensity to reference peak of Y*0.17 and accepting variability of 40.0% in relative intensity.

```
Finding MS1 peak at m/z X-4.0 with a 0.2 m/z tolerance and an expected relative
intensity to reference peak of Y*0.17 and accepting variability of 40.0% in
relative intensity.

Finding MS2 spectra with a precursor m/z X with a 4.0 m/z tolerance.
Enabling variable X with range (400.0, 900.0).
```

**MassQL Query Link**

https://gnps.ucsd.edu/ProteoSAFe/status.jsp?task=96b3bdca0ba74856bf2bf54bc98eaa22

**Additional Data Analysis**

n/a

**Data Availability**

MSV000088461

##### 3.6.1 Chlorinated Compounds in Lichen Thalli Extracts: Monochlorinated Compounds

**Author/s: Kyo Bin Kang**

| |
|---|
| **MassQL Query**<br>`QUERY scaninfo(MS1DATA) WHERE`<br>`MS1MZ=X:TOLERANCEMZ=0.1:INTENSITYPERCENT=25:INTENSITYMATCH=Y:INTENSITYMATCHREFERENCE AND`<br>`MS1MZ=X+2:TOLERANCEMZ=0.1:INTENSITYMATCH=Y*0.33:INTENSITYMATCHPERCENT=30 AND`<br>`MS2PREC=X FILTER MS1MZ=X` |
| **MassQL Translation**<br>Returning the scan information on MS1.<br>The following conditions are applied to find scans in the mass spec data.<br>Finding MS1 peak at m/z X a 0.1 m/z tolerance and a minimum percent intensity relative to base peak of 25.0%and an expected relative intensity to reference peak of Y and this peak is used as the intensity reference for other peaks in the spectrum.<br>Finding MS1 peak at m/z X+2.0 a 0.1 m/z tolerance and an expected relative intensity to reference peak of Y*0.33 and accepting variability of 30.0% in relative intensity.<br>Finding MS2 spectra with a precursor m/z X.<br>Finding MS1 peak at m/z X. |
| **MassQL Query Link**<br>https://proteomics2.ucsd.edu/ProteoSAFe/status.jsp?task=fdde552899594d5b9ccfc7050fabf3b3 |
| **Additional Data Analysis**<br>n/a |
| **Data Availability**<br>MSV00087749 |

To natural product chemists, halogenated compounds have been molecules of interest due to their high potential of bioactivity. Here, we applied MassQL to discover chlorinated compounds from a collection of lichen thalli extracts. We identified several monochlorinated, dichlorinated, and trichlorinated compounds from the data as shown below.

##### 3.6.2 Chlorinated Compounds in Lichen Thalli Extracts: Dichlorinated Compounds

**Author/s: Kyo Bin Kang**

| **MassQL Query** |
| --- |
| `QUERY scaninfo(MS1DATA) WHERE MS1MZ=X:TOLERANCEMZ=0.1:INTENSITYPERCENT=25:INTENSITYMATCH=Y:INTENSITYMATCHREFERENCE AND MS1MZ=X+2:TOLERANCEMZ=0.1:INTENSITYMATCH=Y*0.66:INTENSITYMATCHPERCENT=30 AND MS1MZ=X+4:TOLERANCEMZ=0.1:INTENSITYMATCH=Y*0.11:INTENSITYMATCHPERCENT=30 AND MS2PREC=X FILTER MS1MZ=X` |
| **MassQL Translation** |
| Returning the scan information on MS1.<br>The following conditions are applied to find scans in the mass spec data.<br>Finding MS1 peak at m/z X a 0.1 m/z tolerance and a minimum percent intensity relative to base peak of 25.0% and an expected relative intensity to reference peak of Yand this peak is used as the intensity reference for other peaks in the spectrum.<br>Finding MS1 peak at m/z X+2.0 a 0.1 m/z tolerance and an expected relative intensity to reference peak of Y*0.66 and accepting variability of 30.0% in relative intensity.<br>Finding MS1 peak at m/z X+4.0 a 0.1 m/z tolerance and an expected relative intensity to reference peak of Y*0.11 and accepting variability of 30.0% in relative intensity.<br>Finding MS2 spectra with a precursor m/z X.<br>Finding MS1 peak at m/z X. |
| **MassQL Query Link** |
| https://proteomics2.ucsd.edu/ProteoSAFe/status.jsp?task=40eaab2ce6e84febbffb4b97904a5d65 |
| **Data Availability** |
| MSV00087749 |

##### 3.6.3 Chlorinated Compounds in Lichen Thalli Extracts: Trichlorinated Compounds

**Author/s: Kyo Bin Kang**

**MassQL Query**

```
QUERY scaninfo(MS1DATA) WHERE
MS1MZ=X:TOLERANCEMZ=0.01:INTENSITYPERCENT>80:INTENSITYMATCH=Y:INTENSITYMATC
HREFERENCE AND
MS1MZ=X+2:TOLERANCEMZ=0.01:INTENSITYMATCH=Y:INTENSITYMATCHPERCENT=10 AND
MS1MZ=X+4:TOLERANCEMZ=0.02:INTENSITYMATCH=Y*0.33:INTENSITYMATCHPERCENT=20
```

**MassQL Translation**

```
Returning the scan information on MS1.
The following conditions are applied to find scans in the mass spec data.
Finding MS1 peak at m/z X a 0.01 m/z tolerance and a minimum percent
intensity relative to base peak of 80.0% and an expected relative intensity
to reference peak of Y and this peak is used as the intensity reference for
other peaks in the spectrum.
Finding MS1 peak at m/z X+2.0 a 0.01 m/z tolerance and an expected relative
intensity to reference peak of Y and accepting variability of 10.0% in
relative intensity.
Finding MS1 peak at m/z X+4.0 a 0.02 m/z tolerance and an expected relative
intensity to reference peak of Y*0.33 and accepting variability of 20.0% in
relative intensity.
```

**MassQL Query Link**

https://proteomics2.ucsd.edu/ProteoSAFe/status.jsp?task=7b9142fedd44481996fc256c3c8c3dfa

**Additional Data Analysis**

n/a

**Data Availability**

MSV00087749

Mono-, di-, and tri-chlorinated compounds were found from the lichen extract dataset, by using MassQL queries designed based on the MS1 isotopic patterns (M:M+2 = 3:1, M:M+2:M+4 = 9:6:1, and M:M+2:M+4 = 3:3:1, respectively). Although MS/MS spectral matching to reference spectra of chlorinated depsides or depsidones in LDB[62] gave ambiguous results, we hypothesize that these chlorinated lichen metabolites are depside or depsidone derivatives (**SI Figure 3.6.3 - 1**).

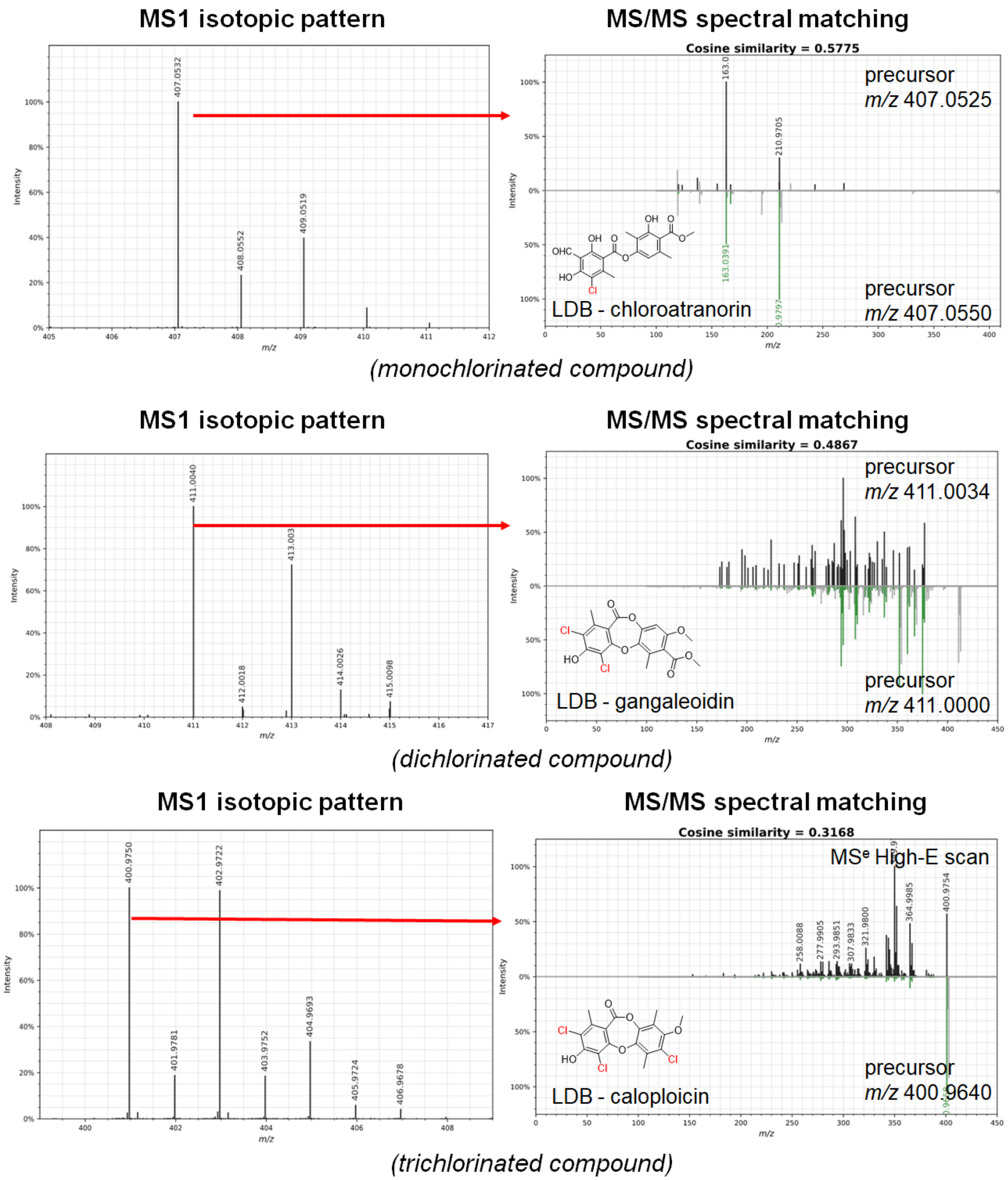


**SI Figure 3.6.3 - 1** - MSQL queries found mono-, di-, and tri-chlorinated depsides and depsidones from lichen specialized metabolite dataset. Chlorinations were confirmed by their MS1 isotopic patterns. Structures of some spectra were annotated by spectral matching against LDB, although they showed relatively low cosine similarities.

##### 3.7.1 Halogenated Compounds: Chloride

**Author/s:** Omri Nahor and Tal Luzzatto Knaan

| |
|---|
| **MassQL Query**<br>`QUERY scaninfo(MS2DATA) WHERE`<br>`MS1MZ=X:INTENSITYMATCH=Y:INTENSITYMATCHREFERENCE:INTENSITYPERCENT=5 AND`<br>`MS1MZ=X+2:INTENSITYMATCH=Y*0.3:INTENSITYMATCHPERCENT=20 AND MS2PREC=X` |
| **MassQL Translation**<br>`Returning the scan information on MS2.`<br>`The following conditions are applied to find scans in the mass spec data.`<br>`Finding MS1 peak at m/z X an expected relative intensity to reference peak of Y and this peak is used as the intensity reference for other peaks in the spectrum and a minimum percent intensity relative to base peak of 5.0%.`<br>`Finding MS1 peak at m/z X+2.0 an expected relative intensity to reference peak of Y*0.3 and accepting variability of 20.0% in relative intensity.`<br>`Finding MS2 spectra with a precursor m/z X.` |
| **MassQL Query Link**<br>https://proteomics2.ucsd.edu/ProteoSAFe/status.jsp?task=6a1b36b121c545989e2716d8268a75ec |
| **Additional Data Analysis**<br>n/a |
| **Data Availability**<br>MSV000078568 |

Halogenated compounds are very common in the marine environment[63] and of special interest as potential active molecules. The naturally abundant isotopic ratio of halogens are distinct and easily identified by MS. The isotopic pattern can vary, depending on the number of halogenated atoms ($^{79}$Br/ $^{81}$Br and $^{35}$Cl / $^{37}$Cl) in the molecule[64].

| #Atoms | Peak Ratio |
|---|---|
| Cl | 1: 0.32 |
| $Cl_2$ | 1: 0.64: 0.102 |
| $Cl_3$ | 1: 0.96: 0.307: 0.033 |
| $Cl_4$ | 0.781: 1: 0.48: 0.102: 0.08 |
| $Cl_5$ | 0.62: 1: 0.64: 0.205: 0.03: 0.002 |

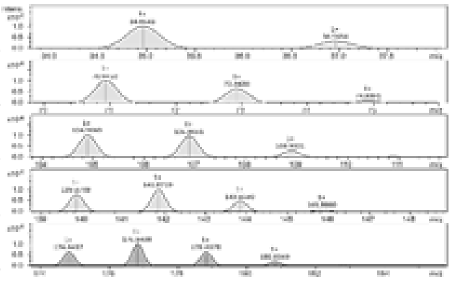

| #Atoms | Peak Ratio |
|---|---|
| Br | 1: 0.973 |
| $Br_2$ | 0.51: 1: 0.486 |
| $Br_3$ | 0.343: 1: 0.973: 0.315 |
| $Br_4$ | 0.176: 0.686: 1: 0.64: 0.15 |
| $Br_5$ | 0.106: 0.514: 1: 0.973: 0.473: 0.092 |

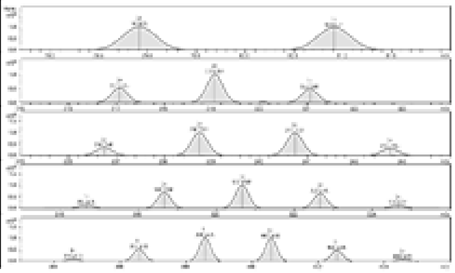

**SI Figure 3.7.1 - 1** - Natural isotopic abundance as reflected by the peak ratios of Chlorine and Bromine

We were interested in mining our marine cyanobacteria dataset for various halogenated molecules. This dataset has previously yielded several novel halogenated compounds that served as a test case for MassQL efficacy. Using MassQL has highlighted additional halogenated candidates for future studies.

Several misannotations are expected as the Br (M/M+2) ratio is similar to $Cl_3$ (M/M+2). Yet, both indicate the presence of halogens.

MassQL was tested in comparison the standard GNPS molecular networking data to validate the halogen signature with  annotated known compounds.

Full Data Network Link

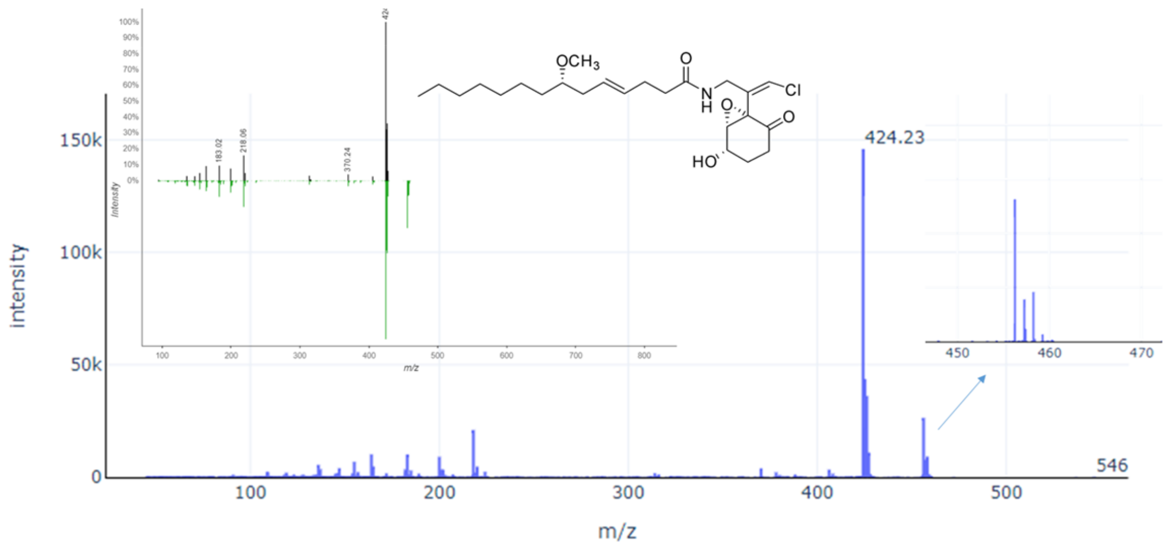


**SI Figure 3.7.1 - 2**: Annotation of Malyngamide C ($C_{24}H_{38}ClNO_5$) by the single chlorine MassQL query. Spectrum with isotopic pattern and mirror plot as annotated by GNPS library

#### 3.8 MassQL Query for Stable Isotope Labeling and Compound Specific Fragment Analysis for Relevant Xenobiotic Metabolites

**Author/s:** Chris Brown, Deepa Acharya, Tao Xu, Ken Clevenger, Quanbo Xiong, Jeff Gilbert

**MassQL Query**

```
QUERY scaninfo(MS2DATA) WHERE
MS1MZ=X:INTENSITYMATCH=Y:INTENSITYMATCHREFERENCE AND
MS1MZ=X+6:INTENSITYMATCH=Y:INTENSITYMATCHPERCENT=30 AND MS2PREC=X AND
MS2PROD=285.0001:TOLERANCEMZ=0.1
```

**MassQL Translation**

```
Returning scans where the following criteria are met.
MS2 acquired on a precursor X that has a signal Y that is m/z 6 shifted at
a relative intensity equal to X.
Finding a MS2 peak at 285.0001 with a 0.1 Da tolerance.
```

The identification of major metabolites related to an active ingredient is a requirement for registration of agrochemicals around the world. These studies often require trace level identification of metabolites from complex environmental matrices. One tool that can aid in these studies is the incorporation of stable-isotope labeled blends of parent material. In one example, plant cells were dosed with a solution containing a blend of unlabeled starting material and a stable labeled (SL) material, where six $^{12}C$ atoms are replaced by six $^{13}C$ atoms. When these were carefully blended, a unique isotope pattern was created that was characteristic of the starting material as it is metabolized. Identification of these isotope patterns has been met with mixed success based on the tools that are provided from commercial vendors.

Active ingredients used for agricultural applications are often dosed at trace levels. Unrelated contaminant species may co-elute with the peaks of interest, causing spurious observation of a peak matching the expected isotope pattern. Careful data analysis is often required to identify peaks that co-elute with the unique label (M and M+6), thus ensuring identification of true metabolites. In addition to the MS1 level label, characteristic fragment ions from the MS/MS spectra may provide additional filters. The combination of a MS1 level SL analysis and characteristic fragment ion analysis will be explored using the MassQL workflow.

Plant cell cultures (wheat, soybean, or blackgrass cell lines) were dosed with the active ingredient of the Arylex herbicide. Metabolites generated from that experiment were extracted, chromatographically separated and analyzed on a Thermo Fusion Lumos in positive mode using standard data dependent acquisition methods. Resulting data files were converted to mzML format with ProteoWizard msconvert tool and uploaded to Ometa Labs for analysis using the MassQL workflow.

From the MassQL filtered MS/MS spectra, we were able to generate a molecular family of species that appear to be related to the metabolism of the parent material dosed into a soybean cell line.

This query is shown to correctly filter species that have the precursor isotope pattern of interest. The substructure that is labeled in the stable isotope approach is also incorporated as part of the fragment ion used for additional filtering (*m/z* 285). This provides further evidence that the spectra incorporated into the molecular network generated from the MassQL filter are likely to originate from the active ingredient that is being studied and can aid in focusing additional structure elucidation efforts.

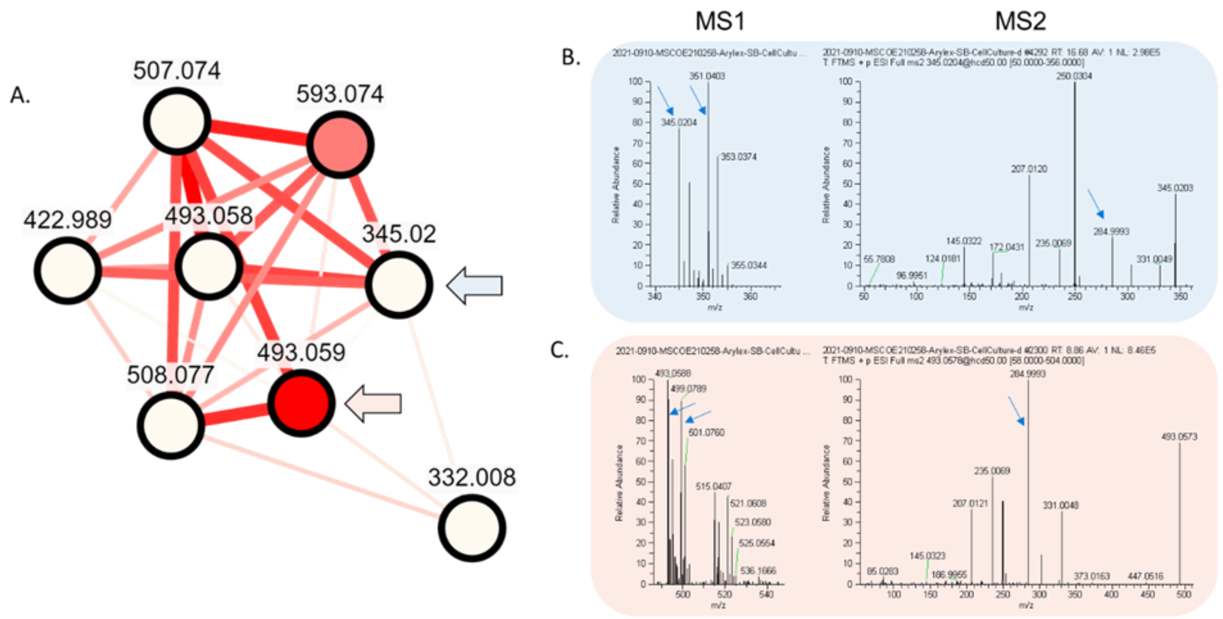


**SI Figure 3.8 - 1** – A molecular network family built from spectra that have been filtered using the MassQL workflow for a MS1 stable isotope pattern and MS2 characteristic fragment ion is shown in A. Nodes are filled using an intensity dependent gradient, and connected using edges that reflect the score between nodes. Nodes expanded upon in panels B and C are marked with arrows showing the stable isotope pattern (M and M+6) and the characteristic fragment ion (m/z 285). MS1 and MS2 spectra of the initial herbicide molecule ([M+H]+ = 345.02) are shown in B. MS1 and MS2 representing a metabolite of interest ([M+H]+ = 493.059) are shown in C.

##### 3.9 Application of MassQL for Mass Defect Filtering - Searching for Acylphloroglucinolated Catechins from *Agrimonia pilosa*

**Author/s:** Hyun Woo Kim

| **MassQL Query** |
| --- |
| `QUERY scansum(MS1DATA) FILTER MS1MZ=515:TOLERANCEMZ=35:MASSDEFECT=massdefect(min=0.1332, max=0.2112)` |
| **MassQL Translation** |
| `Returning the summed scan information on MS1. The following conditions are applied to find scans in the mass spec data. Finding MS1 peak at m/z 515.0 with a 35.0 m/z tolerance and a mass defect minimum of 0.1332 and maximum of 0.2112.` |
| **MassQL Query Link** |
| https://proteomics2.ucsd.edu/ProteoSAFe/status.jsp?task=e4391891497b469b8b049c954136bdd5 |
| **Additional Data Analysis** |
| https://gnps-tableviewer.ucsd.edu/?usi1=https%3A%2F%2Fproteomics2.ucsd.edu%2FProteoSAFe%2Fresult.jsp%3Ftask%3De4391891497b469b8b049c954136bdd5%26view%3Dquery_results&plot_type=heatmap&columnx=rt&columny=i&columnfacet=&columnboxgroup= |

The term 'mass defect' originates from the fact that only the mono isotopic element $^{12}C$ has an integer value for atomic weight (12.000000). The mass defect filtering (MDF) technique was developed for metabolite detection purposes based on a narrow and well-defined mass defect range between the parent drug and its metabolites in pharmaceutics. Secondary metabolites in natural products classified into several families and the components in the same family usually share the same carbon skeleton or substructures. Consideration of filter reference and corresponding substituents would be able to define a mass defect window for certain homologues components.

| Element | Nuclide | Nominal Mass | Exact Mass | Mass Defect |
|---|---|---|---|---|
| Hydrogen | H | 1 | 1.0078 | 0.0078 |
| Carbon | $^{12}C$ | 12 | 12.0000 | 0.0000 |
| Nitrogen | $^{14}N$ | 14 | 14.0031 | 0.0031 |
| Oxygen | $^{16}O$ | 16 | 15.9949 | -0.0051 |
| Fluorine | $^{19}F$ | 19 | 18.9984 | -0.0016 |
| Sulfur | $^{32}S$ | 32 | 31.9721 | -0.0279 |
| Chlorine | $^{35}Cl$ | 35 | 34.9689 | -0.0311 |

**SI Figure 3.9 - 1**: Exact mass and mass defect of common elements.

Pilosanols are novel acylphloroglucinolated catechins which are observed characteristically in the Rosaceae plant, *Agrimonia pilosa*. The structural diversity of pilosanols results from acyl groups on phloroglucinol units. In order to filter the pilosanol peaks from TIC chromatograms of *A. pilosa* extract, we set the filter reference and substituents of pilosanol derivatives for MDF application.

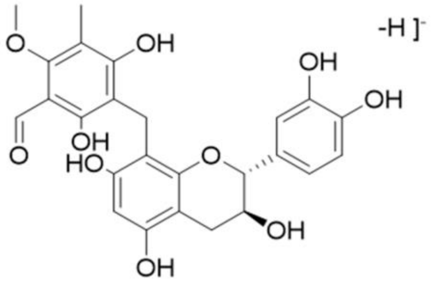


Chemical Formula: $C_{25}H_{23}O_{10}$
*m/z* 483.1291

| Substituent (R) | Formula change | Mass Defect Shift (Da) | min | max |
|---|---|---|---|---|
| H | +H | + 0.0078 | × 1 | |
| Methyl | +CH2 | +0.0157 | | |
| Ethyl | +C2H4 | +0.0313 | | |
| Propyl | +C3H7 | +0.0548 | | |
| Butyl | +C4H10 | +0.0783 | | × 1 |
| MDF setting | 0.1722 ± 39 mDa / M.W 480 ~ 550 | | | |

**SI Figure 3.9 - 2**: Filter reference and substituents of pilosanol derivatives for MDF

After processing the TIC chromatogram by MDF, most of the peaks were discarded and pilosanol derivative peaks remained in the filtered chromatogram (red box). The result was visualized by the “Table Viewer Dashboard”. From further isolation, these peaks were identified as pilosanols A-C and epipilosanols A-C and N.

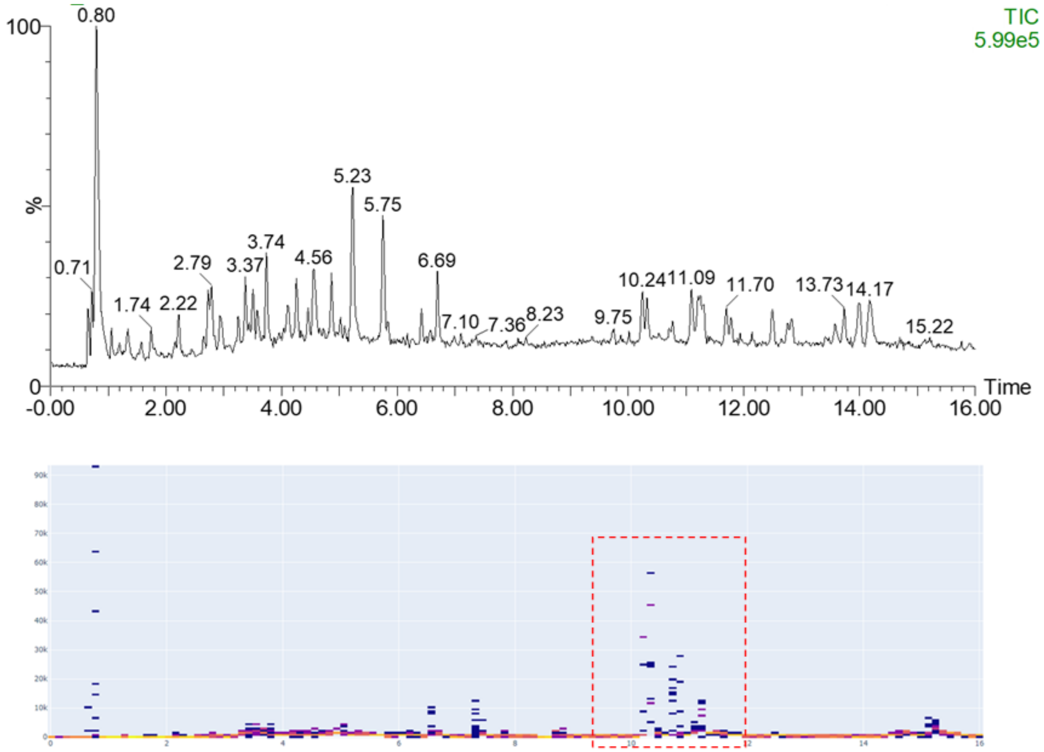


**SI Figure 3.9 - 3**: TIC chromatogram (above) and the filtered TIC chromatogram (below) of the extract of the aerial part of *A.pilosa*.

### 4 MassQL Query and Ion Mobility - Mass Spectrometry Data

#### 4.1 MassQL to Search for Perfluoroalkyl and Polyfluoroalkyl Substances (PFAS) with Ion Mobility in Parallel Accumulation and Serial Fragmentation (PASEF) Data

**Author/s: Steffen Heuckeroth**

| **MassQL Query** |
|---|
| `QUERY scaninfo(MS2DATA) WHERE MS2PREC=X AND MOBILITY=range(min=X*0.0006775+0.40557, max=X*0.00078231+0.48817) AND X=massdefect(min=0.9, max=0.99)` |
| **MassQL Translation** |
| `Returning the scan information on MS2.`<br>`The following conditions are applied to find scans in the mass spec data.`<br>`Finding MS2 spectra with a precursor m/z X.`<br>`Finding spectra with a range of ion mobility (X*0.0006775+0.40557, X*0.00078231+0.48817).`<br>`Enabling variable X with mass defect minimum of 0.9 and a maximum 0.99.` |
| **MassQL Query Link** |
| https://proteomics2.ucsd.edu/ProteoSAFe/status.jsp?task=ee10eddf76c24357b367d5af297a5436 |
| **Additional Data Analysis** |
| https://gnps.ucsd.edu/ProteoSAFe/result.jsp?task=535d077c5b384eab88a81e404713e5b2&view=group_by_compound |

Per- and polyfluorinated alkyl substances (PFAS) are widespread compounds, despite usage of some particular substances has been restricted by the Stockholm Convention (United Nations, *Treaty Series*, vol. 2256, p. 119;) Since production and use cases of perfluorooctanoic acid (PFOA) and perfluorooctanesulfonic acid (PFOS) and their derivatives have been limited, the industry replaced them by short chained acids such as perfluorohexanesulfonic acid (PFHxS), perfluoroether acids (GenX), perfluorotelomer alcohols and sulfides and others[65]. Some of these replacements serve as biological precursors to the restricted PFOA[66,67]. For example, 8:2 fluorotelomer alcohol is degraded to PFOA and other PFAS during aerobic and anaerobic digestion by sewage sludge in wastewater treatment plants (WWTP) and thus causes a continued release of PFOA into the environment[68]. Analysis and annotation of these compounds serving as precursors or replacements is challenging, because they cannot be unambiguously identified via their isotopic pattern, since fluorine is monoisotopic. Therefore, multiple approaches have been used for the untargeted identification of PFAS in environmental samples, firefighting foams and more[69]. These approaches include filtering for negative or just slightly positive mass defects due to the negative mass defect of fluorine (18.9984 Da), and filtering MS2 spectra for specific fragments, e.g., $[CF_3]^-$, $[C_2F_5]^-$ [70,71]. Furthermore, Kendrick Mass Defect based approaches by setting CF2 as a repeating unit have shown to be useful for homologue analysis[71,72]. Ion mobility spectrometry (IMS) has been shown to be a valuable addition to mass spectrometry in recent years, for example by increasing data dependant MS2 rates in the form of PASEF or by mobility resolved data independent fragmentation such as MSe[73–75]. The mobility of molecules can be used to vaguely group them by their substance class, such as glycans, peptides and lipids, which form different trajectories in the mobility-*m/z* plane. While natural products show only slight differences in their trajectories, PFAS show an immensly different trajectory[76], due to fluorine's high difference in mass but comparably lower difference in size, compared to hydrogen. Therefore, MS2 data of suspected PFAS can be specifically queried via MassQL by specifying a range of mobility depending on the molecular weight of the precursor. This query will form a parallelogram in the mobility-*m/z* plane and filter out most natural products, leaving PFAS and other potential xenobiotics behind. If the compounds are also filtered by precursor mass defect, the query becomes even more specific, providing a way of non-targeted tentative annotation of potential PFAS substances for further investigation. Additionally, the mass defect filter could also be applied to fragment ions, or fragments could be filtered for specific ions such as $[F(CF_2)_n]^-$.

#### Discussion

The dataset consists of a 50 nM PFAS standard mix, a SPE enriched WWTP effluent (according to US EPA 537.1) and a spiked sample of the WWTP effluent. The acquired data was subject to internal recalibration for *m*/*z* (NaFormate clusters) and mobility (Agilent P/N G1969-85000) in DataAnalysis 5.3 (Bruker Daltonics GmbH, Bremen, GER).

**SI Figure 4.1 - 1** plots the determined mobility versus the *m/z* for detected features in the WWTP effluent extract. Every dot represents a feature, whereas the size of the dot represents its intensity. Most of the compounds form a trajectory from a mobility of 0.65 Vs/cm² at *m/z* to 1.5 Vs/cm² at *m/z* 1500. However, the mobilities of some compounds differ from the expected trajectory, which can be used to screen these compounds for PFAS. PFAS show a lower-than-expected mobility for their *m/z* due to their high fluorine content. This could be used to screen complex mixtures such as environmental samples for unknown PFAS substances. Since the timsTOF fleX supports precursor isolation in mobility and *m/z* dimension, the MS2 information can be queried for precursors in a specific mobility range and a certain mass defect. The MassQL query forms a parallelogram in the mobility vs. *m/z* plot (**SI Fig. 4.1 - 1**, orange) and extracts the scan info. Since the acquisition of MS/MS spectra within this region was enforced for demonstration purposes, the amount of 5700 remaining spectra is rather large. Nevertheless, the results can be directly submitted to a workflow on the GNPS website to search for library hits and perform molecular networking. The library matching and MN workflow groups the spiked PFCAs into a single cluster due to their spectral similarity (**SI Fig. 4.1 - 2**). In the future, this workflow could be adapted to filter, group, and identify currently unknown PFAS posing a potential environmental and health hazard.

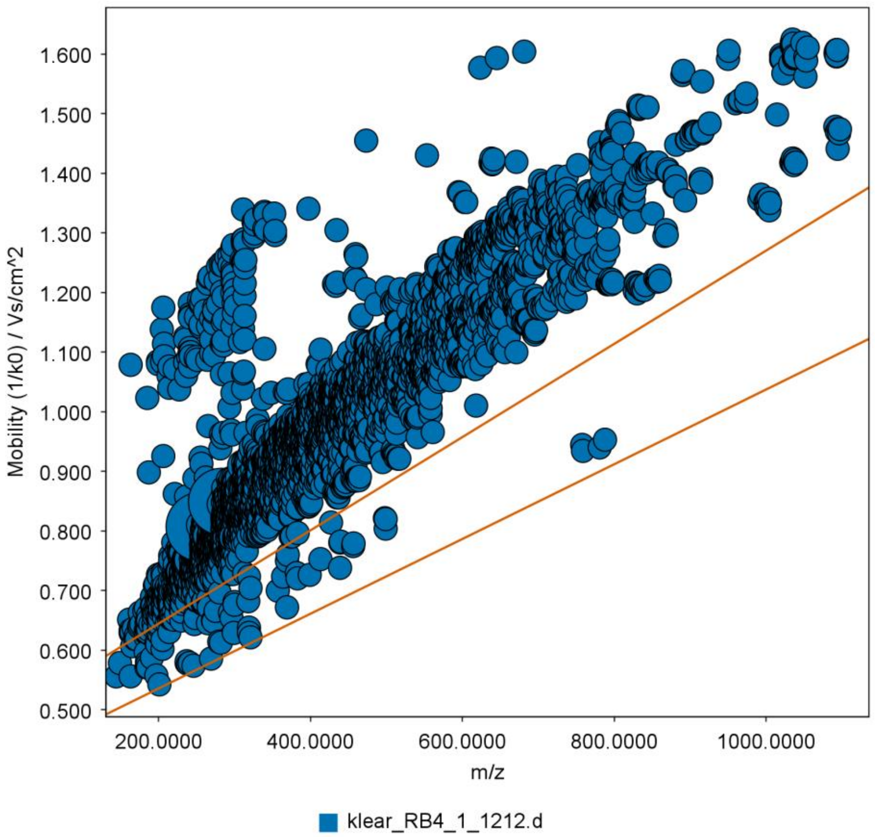


**SI Figure 4.1 - 1**: Plot of the mobility vs. m/z for a detected feature. Most features are located on a diagonal trajectory. However, some features possess a mobility value lower than their *m/z* would suggest, indicated by the orange parallelogram. This region can be specifically queried.

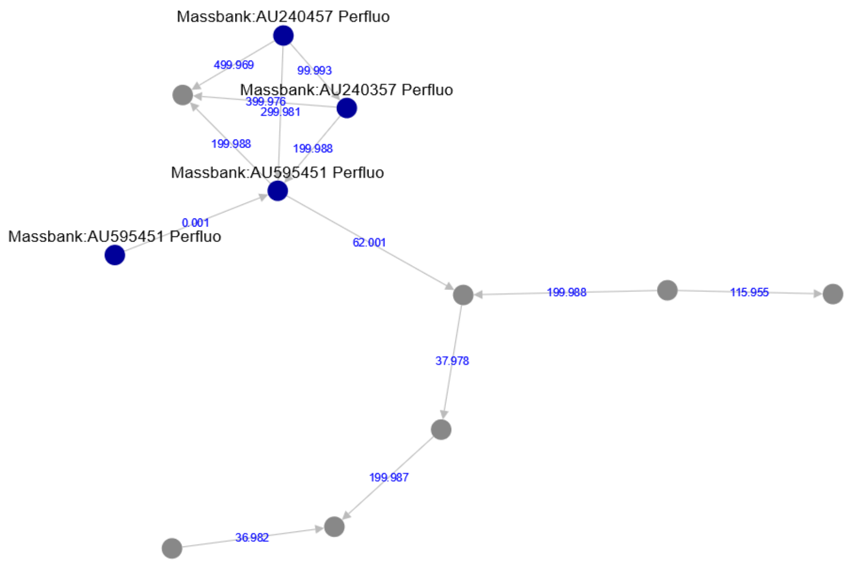


**SI Figure 4.1 - 2**: Molecular networking and library matching result. The spiked PFCAs form a molecular network due to their spectral similarity, some of which have been annotated by library matching.

#### 4.2 Detecting Fungal Metabolites of Interest in Microbial Extracts from Isobars Separated by Trapped Ion Mobility Spectrometry (TIMS)

**Author/s: Gordon T. Luu, Itzel Lizama-Chamu, and Laura M. Sanchez**

| **MassQL Query**<br>`QUERY scaninfo(MS2DATA) WHERE`<br>`MS2PREC=403.15:TOLERANCEMZ=0.1 AND`<br>`MOBILITY=range(min=0.910, max=0.940)` |
|---|
| **MassQL Translation**<br>`Returning the scan information on MS2.`<br>`The following conditions are applied to find scans in the mass spec data.`<br>`Finding MS2 spectra with a precursor m/z 403.15 with a 0.1 m/z tolerance.`<br>`Finding spectra with a range of ion mobility (0.91, 0.94).` |
| **MassQL Query Link**<br>https://proteomics2.ucsd.edu/ProteoSAFe/status.jsp?task=16537d79f72d43c89ec46349925ebd1a |
| **Additional Data Analysis**<br>https://proteomics2.ucsd.edu/ProteoSAFe/status.jsp?task=d0fe853234dc454d8d9d9f0169d8f40c |
| **Data Availability**<br>MSV000088153 |

Isobars are molecules that have identical nominal masses but different fractional masses. Even using high resolution mass spectrometers, isobars become increasingly difficult to distinguish as the difference in their fractional masses converge. Oftentimes liquid chromatography (LC) is able to separate isobaric molecules, and when coupled to tandem mass spectrometry (MS/MS), isobaric precursor molecules can be distinguished via their retention time. However, in certain applications, such as screening hundreds to thousands of chemical extracts for a target compound, chromatography free workflows are desired to allow for high throughput capabilities (i.e. matrix-assisted laser desorption/ionization time-of-flight mass spectrometry; direct infusion mass spectrometry; MALDI-TOF MS; DI-MS). An alternative to distinguishing isobars via retention time is to use collisional cross section obtained from ion mobility spectrometry. The introduction of trapped ion mobility spectrometry (TIMS) to mass spectrometers (i.e. Bruker timsTOF flex MS) reduces the amount of time required for isobaric separation from minutes to milliseconds, which allows for high throughput separation.

Discussion

Here, LC-TIMS-MS/MS data were used to differentiate between two isobaric features: *m/z* 403.16 and *m/z* 403.13. Data were acquired on a Bruker timsTOF flex MS in data dependent acquisition mode using parallel accumulation-serial fragmentation (ddaPASEF) and converted to mzML using TIMSCONVERT. **SI Figure 4.2 - 1A** shows the mobilogram for *Penicillium atramentosum* str. RS17. GNPS analysis revealed that *m/z* 403.16 may be an analogue of meleagrin (**SI Figure 4.2 - 1B**), while *m/z* 403.13 is a potential contaminant introduced into the extract during resuspension. We suspect *m/z* 403.16 could be the result of the loss of a methoxy group from meleagrin. GNPS provided a putative annotation for *m/z* 403.13 (tumanoic acid F), but no structural information could be found for the compound, and comparison of the fragmentation patterns in the sample and library spectra did not reveal well matched fragments (**SI Figure 4.2 - 1C**). A query was formulated to detect the presence of our compound of interest in a large number of spectra in future acquisitions following successive rounds of fractionation.

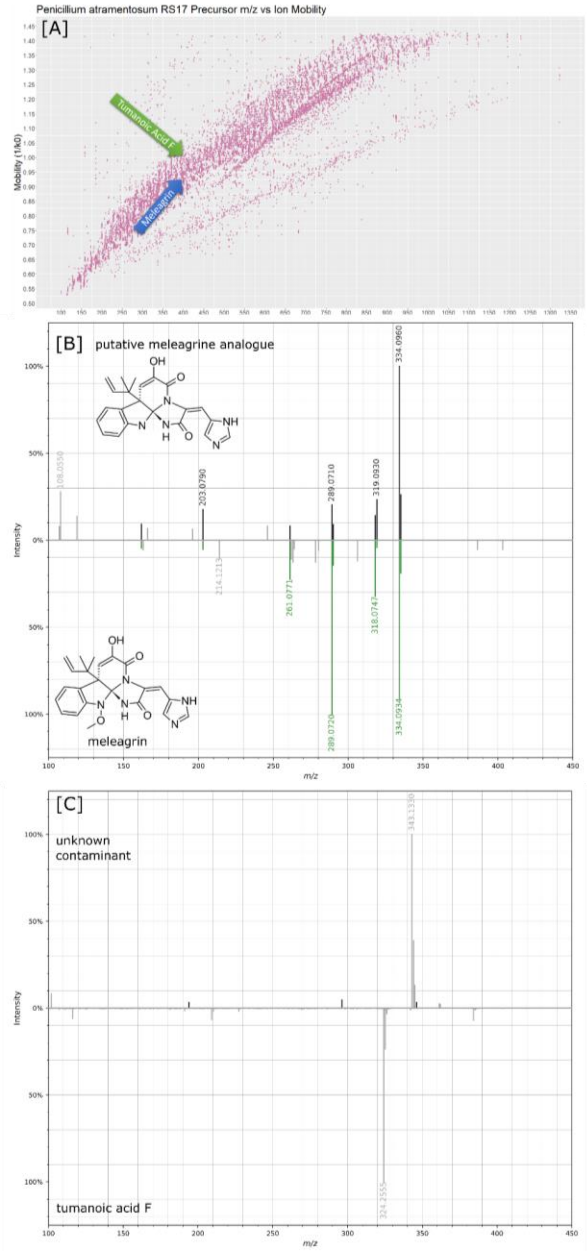

**SI Figure 4.2 - 1**: [A] Plot of mobility ($1/K_0$) vs mass-to-charge ratio (*m/z*). Features *m/z* 403.16 (putative meleagrin analogue) and 403.13 (tumanoic acid F) can be found with most other features located on a diagonal trajectory. Blue and green arrows denote the location of a putative meleagrin analogue and tumanoic acid F, respectively. [B] Mirror plot comparing sample MS/MS spectrum for putative meleagrin analogue (black) to library MS/MS spectrum for meleagrin (green) and their respective (putative) structures. [C] Mirror plot comparing sample MS/MS spectrum for *m/z* 403.13 (black) to tumanoic acid F (green).

### 5 MassQL Query and Gas Chromatography Mass Spectrometry

#### 5.1 Detection of fatty acid ethyl esters (FAEEs), biomarkers for excessive alcohol consumption, using MassQL to query GC-Orbitrap-EI data

**Author/s: Xin Hu**

**MassQL Query**

```
QUERY scaninfo(MS1DATA) WHERE
MS1MZ=88.0473:TOLERANCEMZ=0.01:INTENSITYPERCENT=2 AND
MS1MZ=101.0595:TOLERANCEPPM=5:INTENSITYPERCENT=15 AND
MS1MZ=X:INTENSITYPERCENT=20 AND
MS1MZ=X+14.0157:TOLERANCEPPM=5:INTENSITYPERCENT=5 AND
MS1MZ=X+28.0314:TOLERANCEPPM=5:INTENSITYPERCENT=5 AND
MS1MZ=X+42.0471:TOLERANCEPPM=5:INTENSITYPERCENT=5 AND
MS1MZ=X+56.0628:TOLERANCEPPM=5:INTENSITYPERCENT=5 AND
MS1MZ=X+70.0785:TOLERANCEPPM=5:INTENSITYPERCENT=5 AND
MS1MZ=X+84.0942:TOLERANCEPPM=5:INTENSITYPERCENT=5 AND
MS1MZ=X+98.1099:TOLERANCEPPM=5:INTENSITYPERCENT=5
```

##### MassQL Translation

Returning the scan information on MS1.

The following conditions are applied to find scans in the mass spec data.

Finding MS1 peak at m/z 88.0473 a 0.01 m/z tolerance and a minimum percent intensity relative to base peak of 2.0%.

Finding MS1 peak at m/z 101.0595 a 5.0 PPM tolerance and a minimum percent intensity relative to base peak of 15.0%.

Finding MS1 peak at m/z X a minimum percent intensity relative to base peak of 20.0%.

Finding MS1 peak at m/z X+14.0157 a 5.0 PPM tolerance and a minimum percent intensity relative to base peak of 5.0%.

Finding MS1 peak at m/z X+28.0314 a 5.0 PPM tolerance and a minimum percent intensity relative to base peak of 5.0%.

Finding MS1 peak at m/z X+42.0471 a 5.0 PPM tolerance and a minimum percent intensity relative to base peak of 5.0%.

Finding MS1 peak at m/z X+56.0628 a 5.0 PPM tolerance and a minimum percent intensity relative to base peak of 5.0%.

Finding MS1 peak at m/z X+70.0785 a 5.0 PPM tolerance and a minimum percent intensity relative to base peak of 5.0%.

Finding MS1 peak at m/z X+84.0942 a 5.0 PPM tolerance and a minimum percent intensity relative to base peak of 5.0%.

Finding MS1 peak at m/z X+98.1099 a 5.0 PPM tolerance and a minimum percent intensity relative to base peak of 5.0%.

##### MassQL Query Link

https://proteomics2.ucsd.edu/ProteoSAFe/result.jsp?task=fd0ab40176dd4f7a8ac6cbf7523a1e1a&view=query_results

##### Data Availability

MSV000088046

High resolution mass spectrometry enabled by the Orbitrap technique revolutionizes the accuracy in measurements of *m/z* and coverage of chemicals in complex biological matrices. In addition, collection of all spectral features in full-scan mode provides the greatest benefit by preserving information for unidentified MS features. In this experiment, we used MassQL to query the existing dataset of human plasma collected from GC-Orbitrap to test whether unidentified MS features can be annotated based on spectrum characteristics.

Fatty acid ethyl esters (FAEEs) are ethanol metabolites that remain detectable long after alcohol consumption and after ethanol disappears in the circulation. On that basis, FAEEs in blood samples are used as short-term confirmatory test for ethanol intake. In addition, deposition of FAEEs in tissue organs such as human liver, adipose tissue and hair can be measured as long-term markers of chronically elevated alcohol consumption as well as implications of alcohol-induced organ damage. FAEEs are readily detected by GC-MS. The spectra of straight-chain fatty acid under electron ionization (EI) include characteristic base peaks of *m/z*=88 (formed by McLafferty rearrangement) and long homologous series of related ions that are 14 amu apart (formed by $CH_2$ loss) such as m/z of 87, 101, 115, 129, 143, 157, 199 etc. Loss of ethoxide ion ($[M\text{-}45]^+$) is also characteristic and confirms that the chemical is an ethyl ester.

FAEEs contain ethyl esters of many various saturated and unsaturated, straight chain and branched chain fatty acids. By using the base peaks (*m/z*= 101.0595 and 88.0473) and the loss of ethoxide ion (M- 45.0340) from molecule ion (*m/z* = 239.2369 for ethyl palmitate, *m/z*=211.2062 for ethyl myristate, *m/z*=265.2531 for ethyl oleate), we identify the potential peaks for the three FAEEs in a human plasma sample (**SI Fig. 5.1 - 1**). We then used MassQLto query the characteristic pattern of the spectrum in 80 samples of human plasma as well as standard reference material (SRM) 1958. We obtained clusters of potential hits between retention time of 6 min and 11 min (**SI Fig. 5.1 - 1b**). Using the intensity criteria in MassQL to set the intensity of base peak of m/z 101 at 15% and above is advantageous in differentiating other similar spectrum from FAEEs that may represent fatty acid methyl esters and other interferences. For example, two potential peaks were identified for ethyl palmitate (RT=7.72 min and 8.12 min, Fig Xa and c). At RT=8.12 min, although the intensities of *m/z* 88 and 101 were higher compared to the peak at 7.72 min, *m/z* 101 was at much lower intensity percentage compared to other *m/z* in the spectrum (**SI Fig. 5.1 - 1b**) and provides less similarity compared to the mass spectrum of ethyl palmitate in authentic standard (**SI Fig. 5.1 - 1d**, provided by the NIST Chemistry WebBook, SRD 69). Of note, *m/z* 101 was found to be the most consistently high abundance ion for mid- to long-chain FAEEs and used as a quantifying ion in multiple studies[77–79]. Similar strategies were applied to query omega-3 and omega-6 with base peaks of 108 and 150, respectively. No hits were found among the 80 human plasma samples, possibly due to the low abundance of unsaturated FAEEs. Further studies are of interest to detect FAEEs in samples from alcohol abuse.

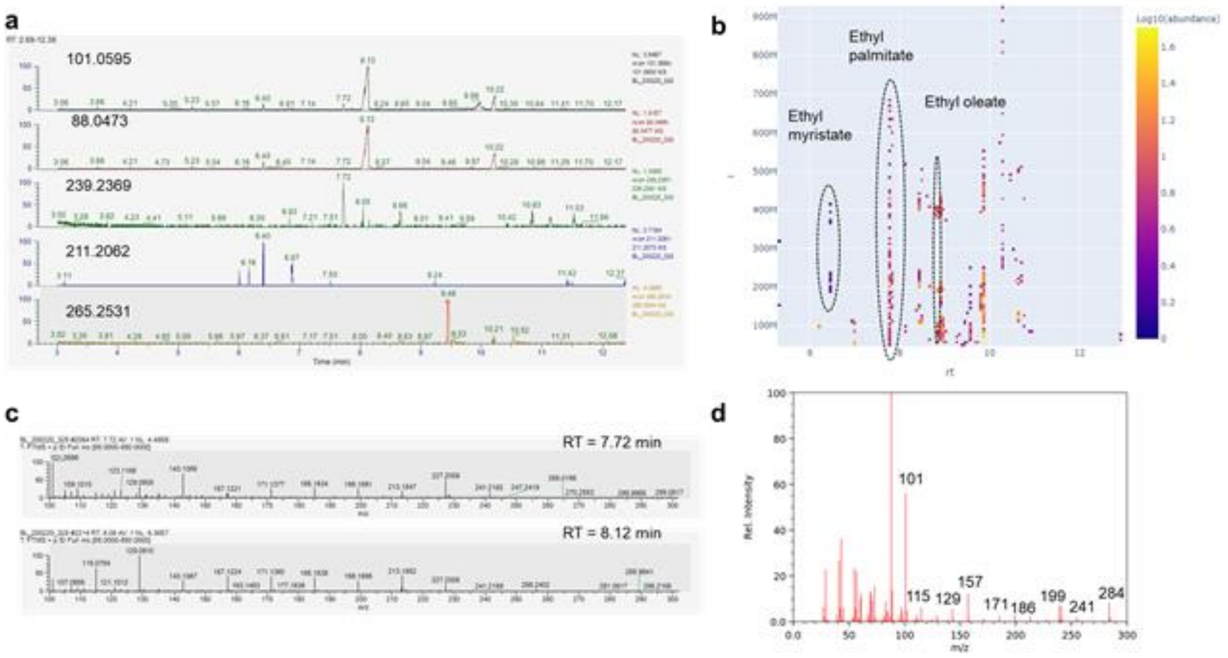


**SI Figure 5.1 - 1**. Detection of ethyl myristate, ethyl palmitate, and ethyl oleate as examples of fatty acid ethyl esters (FAEEs) in human plasma samples (n=80) from an archival repository. **a** Extracted ion chromatography (EIC) showed potential peaks of the three FAEEs by using the characteristic base ions (101.0595, 88.0473) and loss of ethoxide ions from the molecule ions (239.2369, 211.2062 and 265. 2531). **b** MassQL using the presence of base peaks and serial loss of $CH_2$ in fragmentation showed clusters of potential matches of FAEEs. **c** One of the two potential peaks for ethyl palmitate shown by EIC was not identified by MassQL at RT=8.12 min had less percentage of the base peak 101.0595, indicating that MassQL can be used to filter out similar interfering spectrum that had less resemblance of the authentic standard spectrum (**d**, ethyl palmitate EI spectrum from the NIST Chemistry WebBook).

26. Antimicrobial multidrug resistance in the era of COVID-19: a forgotten plight? | Antimicrobial Resistance & Infection Control | Full Text. https://aricjournal.biomedcentral.com/articles/10.1186/s13756-021-00893-z.

27. Natural products in drug discovery: advances and opportunities | Nature Reviews Drug Discovery. https://www.nature.com/articles/s41573-020-00114-z.

40. Martha Perez Gutierrez, R., Maria Neira Gonzalez, A. & Hoyo-Vadillo, C. Alkaloids from Piper: A Review of its Phytochemistry and Pharmacology. *Mini Rev. Med. Chem.* **13**, 163–

193 (2013).

59. Baars, O., Morel, F. M. M. & Perlman, D. H. ChelomEx: Isotope-Assisted Discovery of

Metal Chelates in Complex Media Using High-Resolution LC-MS. *Anal. Chem.* **86**, 11298–11305 (2014).

79. Politi, P., Rocchetti, M., Emanuele, E., Rondanelli, M. & Barale, F. Randomized placebo-controlled trials of omega-3 polyunsaturated fatty acids in psychiatric disorders: a review of

the current literature. *Curr. Drug Discov. Technol.* **10**, 245–253 (2013).

80. Mojumdar, E. H. *et al.* Stratum corneum lipid matrix: Location of acyl ceramide and cholesterol in the unit cell of the long periodicity phase. *Biochim. Biophys. Acta* **1858**, 1926–1934 (2016).
